# Variations in bacterial community at Chlorophyll Maximum (C-Max) depths along the west coast of India due to seasonal changes in the primary productivity

**DOI:** 10.1101/2023.07.05.547789

**Authors:** Ashutosh Shankar Parab, Mayukhmita Prithunandan Ghose, Cathrine Sumathi Manohar

**Author notes:** Corresponding author, ORCID: 0000-0003-0178-7784.

## Abstract

An increase in primary productivity is recorded annually during the southwest monsoon season along the west coast of India, an important upwelling zone. The influence of the seasonal variations in the *in situ* primary productivity on the bacterial dynamics and community structure was explored during the non-monsoon and productive monsoon seasons. In the monsoon season, shallower mixed layer depth, increased nutrient concentration and a significant, positive correlation of bacterial carbon (p < 0.01) with primary productivity was observed. Bacterial diversity was assessed in the chlorophyll maxima depths during both seasons based on next-generation, metagenomic analysis. In the non-monsoon season, genera such as *Idiomarina*, *Salinimonas*, *Marinobacter* of Proteobacteria and *Bacillus* and *Lactobacillus* of Firmicutes were dominant. These major bacterial genera are shown through canonical correspondence analysis to play an important ecological role. They could be responsible for the increased heterotrophic activity recorded through predicted functional gene profiles in this season. In the monsoon season, increased abundance in the autotrophic Cyanobacteria community and its photosynthetic activity was recorded in the gene profiles. Higher diversity of heterotrophic phyla such as Bacteroidetes, Actinobacteria and a few Candidatus phyla and changes in the diversity of Proteobacteria with a representation of *Alteromonas*, SAR86 clade and OM60 (NOR5) was observed. These results highlight the bacterial dynamics associated with seasonal variations in primary productivity along the west coast of India.

## 1. Introduction

Upwelling plays a crucial role in the biogeochemical processes, as it provides essential nutrients for phytoplankton growth, which in turn supports the marine food chain (Capone et al., 2013). Along the west coast of India (WCI), upwelling occurs annually from June to September, during the southwest monsoon season (Smitha et al., 2008; Gupta et al., 2016). The upwelling in this region is driven by the strong, persistent winds that blow parallel to the coast, which create a coastal upwelling front (Schott et al., 2009). These winds push the surface waters away from the coast, allowing the colder, nutrient-rich waters from the deeper ocean to rise to the surface (Vinayachandran et al., 2021; Gupta et al., 2021). This brings in nutrient-rich water to the surface that is abundant in nutrients such as nitrates and phosphates, which leads to an increase in phytoplankton productivity. Subsequently, this results in the generation of substantial amounts of organic matter along the WCI during the southwest monsoon season (Kumar et al., 2000; Ramaiah et al., 1996; Habeebrehman et al., 2011). Bacteria, which are referred to as the “Unseen Majority” play crucial roles in the biogeochemical cycling of organic matter and are therefore fundamental for the operation of marine ecosystems (Whitman et al., 1998). The heterotrophic bacteria through the “microbial loop” break down the particulate organic matter into the dissolved organic matter that sustains the marine food web and nutrient cycling (Pomeroy et al., 2007). Thus providing necessary nutrients for the primary production of phytoplankton, which forms the basis of the aquatic food web (Wiggert et al., 2005; Azam and Malfatti, 2007). Evaluating the processes that influence the structure of bacterial communities on spatial and temporal scales is essential for understanding the recycling of organic matter in marine ecosystems (Danovaro et al., 2017).

Bacteria play critical roles in regulating global nutrient cycles and primary production (Baltar et al., 2015). Therefore, any changes in their abundance or activity as well as the surrounding environmental changes can have significant effects on the functioning of marine ecosystems. Initial studies of the bacterial community analysis from major upwelling regions across the globe, was carried out based on conventional cloning and PCR-based methods, has studied specific bacterial groups and diversity in the various upwelling regions have shown seasonal variations in dominant bacterial communities, including Alphaproteobacteria, Gammaproteobacteria, Bacteroidetes and certain Planctomycetes (Kuypers et al., 2005; Cai et al., 2007; Alonso-Gutiérrez et al., 2009). This signified their involvement in biogeochemical cycling and highlighted the impact of environmental variations on the bacterial diversity. However, studies carried out based on recent Next-generation sequencing (NGS) technologies have provided valuable advancements in understanding the bacterial communities in upwelling regions, allowing in-depth examination of the bacterial dynamics (Jing et al., 2013; Bergen et al., 2015; Thiele et al., 2019; Liu et al., 2022). These studies have found bacterial communities including Proteobacteria, Cyanobacteria, Bacteroidota, Firmicutes, Actinobacteria and certain previously unidentified Candidatus phyla prevalent in diverse upwelling regions, highlighting their ecological significance. NGS technologies have significantly enhanced the understanding of the active members within bacterial communities, revealing the ecological significance of rare phylotypes such as SAR11, Alteromonadales and Vibrionales in their response to environmental changes (Gutiérrez-Barral et al., 2021; Mena et al., 2022). Additionally, the impact of upwelling on microbial community composition has also been observed at C-Max depths (Gong et al., 2022). Collectively, these studies underscore the variability and complexity of bacterial community structure and abundance within upwelling zones shaped by distinct environmental factors and seasons.

Studies on bacterial communities along the WCI have primarily focused on specific regions, such as the oxygen minimum zones (Bandekar et al., 2018; Fernandes et al., 2020; Lincy and Manohar, 2021), the mud bank regions (Anas et al., 2021; Vijayan et al., 2022) and estuarine habitats (Khandeparker et al., 2017; Kuchi et al., 2021). Studies based on culturable approaches have shown the influence of monsoonal upwelling and higher productivity on the bacterial community structure and functional diversity in the estuarine, coastal and open ocean waters along the WCI (Anas et al., 2021; Vijayan et al., 2022; Parab et al., 2022). Our prior investigation, on the culturable bacterial diversity along the WCI, showed higher bacterial abundance during the monsoon season and with the highest colony-forming units (CFUs) from the C-Max depths (Parab et al., 2022). The C-Max, is a prominent feature of tropical marine environments occurring between 20-100 m water depth with maximum primary productivity, due to the optimal physicochemical conditions for the growth of phytoplankton (Cullen, 2015). These C-Max depths have very high a high availability of organic substrate for bacteria. However, there are limited research from the upwelling regions from the WCI, on the impact of varying primary productivity on the bacterial communities from C-Max zones utilising high-throughput sequencing techniques. To address this knowledge gap, this study aimed to comprehensively analyse the impact of increased productivity and surrounding environmental factors on the bacterial dynamics along one of the most productive zone WCI, during non-monsoon and monsoon seasons, using high-throughput sequencing of 16S rRNA amplicons. This approach allows an in-depth understanding of the regional bacterial community structure and its potential seasonal variations. This study provides the first detailed analysis of the bacterial community and functional diversity associated with seasonal changes in the primary productivity along the WCI.

## 2. Methodology

### 2.1 Study location, sample collection and assessment of physicochemical parameters

Sampling was conducted during oceangraphic cruises onboard RV Sindhu Sankalp during the non-monsoon season in February 2019, Cruise # SSK-125 and in the southwest monsoon season in September 2019, Cruise # SSK-131 along the WCI. Samples were collected between 8°N and 16°N and 72°N to 76°E, at three stations: off Goa, Mangalore and Trivandrum (Fig. S1). At each station, samples were collected from a coastal location at 30 m water depth and an off-shore location at 600 m depth. The sampling locations in the coastal and off-shore regions of Goa are designated as G30 and G600 respectively. Similar to this, sampling locations off Mangalore and Trivandrum stations are designated as M30, M600 and T30, T600 respectively (Fig. S1). At each of the six sampling locations, the conductivity-temperature-depth (CTD) underwater profiler, Sea-Bird, USA, model: SBE-911 plus with sensor accuracy of ± 0.0003 S/m, ± 0.001°C and ± 0.015 % for conductivity, temperature and depth respectively was employed to quantify the physicochemical characteristics. The water temperature, density, salinity, dissolved oxygen (DO) and fluorescence, which represents the concentration of chlorophyll in the water column was measured. In all coastal locations (30 m contour), samples were collected from the (i) surface at about 2 m below the sea surface, (ii) the depth at which maximum concentration of chlorophyll was recorded (C-Max) and (iii) bottom depth at about 3 m above the seafloor. In the off-shore region along the 600 m contour, samples were collected from (i) surface, (ii) C-Max, (iii) 100 m, (iv) 200 m (v) 400 m and (vi) bottom depth. The depth of C-Max was defined based on the readings obtained from the fluorescence sensor during the downward haul of the CTD underwater profiler and samples were collected at desired depths during the upward haul. Water samples collected from all the depths were immediately sub-sampled and appropriately stored for subsequent examination. Nutrient analysis was carried out in an automated continuous flow analyzer (SKALAR 5000, Netherlands) to measure the levels of nitrate, nitrite, phosphate and silicate, present in water samples.

### 2.2. Estimation of chlorophyll and primary productivity

Chlorophyll-a (Chl-a) concentrations in the water column at discreet depths were determined fluorometrically following a standard protocol of the Joint Global Ocean Flux Study (JGOFS, UNESCO 1994). For this, 1 L of seawater sample from each depth was filtered using a 0.4 µm, 47 mm GF/F filter (Whatman, USA) in triplicates. Chl-a was extracted from the filters using 10 mL of 90 % acetone through overnight incubation at 20°C in the dark. After the incubation, the samples were vortexed and centrifuged to remove cellular debris. The clear supernatant was collected and used for the estimation of chl-a concentration on a Trilogy Laboratory Fluorometer (Turner Designs, USA). The water column chl-a concentration from the euphotic zone was determined through the integration of chl-a values from surface to 100 m depth and were expressed as mg m^-2^. Gross primary productivity (PP) was assessed in water samples from coastal and off-shore locations in both seasons based on a ^14^C-labelled method using labeled inorganic C, NaH^14^CO_3_ following the standard protocol of the JGOFS (UNESCO 1994). For this, water samples were collected and incubated without any pre-filtration in three light and one dark, 250 mL polycarbonate bottle (Nalgene, Germany), from all the sampling depths of the coastal (3 depths) and surface, C-Max, 100 and 200 m depths from the off-shore locations. Each bottle was supplemented with one ampoule of NaH^14^CO_3_, obtained from the

Board of Radiation and Isotope Technology in Mumbai, with a specific activity of 185 kBq. The bottles were then incubated at the respective water depths using an *in situ* mooring system for 12 daylight hours. At the end of the incubation, water samples were filtered using GF/F 47 mm diameter, 0.7 m pore size filters (Whatman, USA) and stored in scintillation vials in the dark till further analysis. For estimation of the radioactivity, filter papers were dissolved overnight using concentrated HCl followed by incubation in recommended volume of liquid scintillation cocktail (Sisco Research Laboratory, Mumbai, India). The levels of radioactivity were assessed on a scintillation counter (PerkinElmer Wallac 1409 DSA, USA) and the values were expressed as mg C m^-3^ d^-1^. To determine depth-integrated PP, productivity measurements up to 100 m depth were taken and integrated to obtain water column productivity and expressed as mg C m^-2^ d^-1^.

### 2.3. Estimation of bacterial productivity and dynamics

To estimate the rate of bacterial productivity (BP) at coastal and off-shore stations, water samples from all the sampling depths were collected during both seasons and subjected to the ^3^H-thymidine incorporation method as outlined in the JGOFS protocols (UNESCO 1994). Following the protocol, water samples (20 mL) in triplicates from each depth were incubated in the dark with ^3^H-thymidine at final concentration of aboute 0.6 nM (specific activity 18 Ci mmole^-1^, Bhabha Atomic Research Center, Mumbai) for 2 hours at ambient temperature. The incubation was halted by adding filtered formaldehyde and the samples were filtered using 0.22 µm cellulose acetate, 25 mm diameter filters (Millipore, India). The filters were washed with cold trichloroacetic acid (10 % w/v) followed by ethanol (70 % v/v) and were stored in scintillation vials for further analysis. The filter papers were soaked overnight in ethyl acetate and then a 4 mL of liquid scintillation cocktail was added and radioactivity was quantified using a scintillation counter (PerkinElmer Wallac 1409 DSA, USA). BP was calculated utilizing an average oceanic conversion rate of 2.17 × 10^18^ cells mol^-1^ of thymidine incorporated (Ducklow in 1993) and the results were expressed as mg C m^-3^ d^-1^.

Bacterial abundance was estimated using epifluorescence microscopy (Porter and Feig 1980). For this, 20 mL water subsamples from each depth were preserved with buffered formaldehyde to achieve a 2 % final concentration. Triplicate aliquots of the preserved samples, each measuring 2 - 5 mL, were stained with 4’6-diamidino-2-phenylindole (DAPI) to reach a final concentration of 1 μg mL ^-1^. These samples were then filtered using black polycarbonate membrane filters with a pore size of 0.22 μm, (Millipore, U.S.A.). The bacterial counts were carried out using an epifluorescence microscope (Olympus BX-51, Japan) and expressed in terms of the number of bacterial cells L^-1^. The bacterial counts were also converted to bacterial carbon (BC) using the conversion factor of 20 fg cell ^-1^ (Ducklow et al., 1995) and reported as mg C m^-3^. The total plates counts (TPC) of heterotrophic bacteria present in the water samples were also determined based on the spread plate technique on Zobell Marine Agar (ZMA) expressed as colony-forming units (CFUs mL^-1^) (Parab et al., 2022).

### 2.4. Bacterial community and functional diversity analysis

#### 2.4.1. Extraction of metagenomic DNA, library preparation and Illumina sequencing

For extraction of metagenomic DNA, 20 L of water samples from C-Max depths at each location were filtered through 0.22 mm Durapore filters (Millipore, USA) using a Peristaltic pump (Millipore, USA) on board the research vessel. The filters were immediately stored in cryovials at −80°C in a DNA storage buffer containing 0.75M sucrose, 40 mM EDTA and 50 mM Tris (pH 8.3). DNA extraction was carried out using a FastDNA^TM^ SPIN kit (MP Biomedical, USA) according to the manufacturer’s protocol. The quality of the metagenomics DNA extracted was tested and a sufficient quantity of DNA was utilized for library preparation, which targeted the 16S rRNA gene V3-V4 region using specific proprietary primers (16S amplicon PCR forward primer: 5’ TCGTCGGCAGCGTCAGATGTGTATAAGAGACAGCC TACGGGNGGCWGCAG 3’ and 16S amplicon PCR reverse primer: 5’ GTCTCGTGGGCTC GGAGATGTGTATAAGAGACAGGACTACHVGGGTATCTAATCC 3’). Illumina PCR sequencing adapter sequence used were, Index 1 Read 5′ CAAGCAGAAGACGGCATA CGAGAT[i7]GTCTCGTGGGCTCGG and Index 2 Read 5′ AATGATACGGCGACCACCGAGATCTACAC [i5] TCGTCGGCAGCGTC. (Qu et al., 2018; Bora et al., 2022)The 16S rRNA gene libraries generated were sequenced on NovaSeq 6000 v1.5 Illumina sequencing platform (Illumina Inc., USA) at the National Institute of Biomedical Genomics, West Bengal, India. The sequences of the amplicons obtained from the C-Max depths from all the locations during both seasons were processed with a standard pipeline using the QIIME2 tool (Bolyen et al., 2019). The raw reads were initially quality filtered and primers were removed using Cutadapt v4.2. After quality filtration, the ASV file was generated using the DADA2 tool v3.16 and taxonomic profiling of the operational taxonomic units (OTU) was carried out against the V3-V4 classifier.

#### 2.4.1. Functional diversity analysis

Functional diversity of bacterial communities was derived from marker gene sequences using the PICRUSt2 software, version 2.4.1. These acquired gene profiles were subsequently categorized into KEGG pathways (Douglas et al. 2020). The PICRUSt tool facilitated the prediction of potential functionalities from 16S rRNA sequence data. Employing the KEGG database, closed reference OTU picking was performed against the Greengenes taxonomy. The predicted functional profiles with gene profiles further grouped into KEGG pathways, following to PICRUSt protocols. Additionally, to measure the variance between reference genomes and the inferred metagenome, NSTI (Nearest Sequenced Taxon Index) scores were determined.

### 2.5. Data analysis

Statistical analyses were performed using SPSS, to investigate the significance of the distribution between physicochemical and biological characteristics in the two seasons using a single-factor analysis of variance (ANOVA). Pearson’s correlation analysis was employed to interpret relationships within the physicochemical and biological parameters between the two seasons, at a significance level of 5 % (P < 0.05). Alpha diversity indices and Beta diversity between two seasons at OUT level were calculated using Non-metric Multidimensional Scaling Analysis (NMDS) based on Bray–Curtis distance using the R package Phyloseq v1.22.3 (McMurdie and Holmes, 2013). A significance in the distribution of bacterial community between non-monsoon and monsoon season were estimated using STAMP bioinformatics software v2.1.3 (Parks et al., 2014). Heatmap of bacterial communities at a class level was plotted using hierarchical clustering analysis based on the Euclidean similarity index in R package Pheatmap v1.0.12. The R packages: ggplots2 and were used for the visualization of the data. All these analyses were performed in R studio v3.5.1 (R Core Team, 2015). Bacterial community structure at the genera level, along with environmental variables of the non-monsoon and monsoon season, were analysed based on canonical correspondence analysis (CCA) using PAST software v3.23 (Hammer et al., 2006).

## 3. Results

### 3.1. Physicochemical parameters

The physicochemical parameters varied significantly between the non-monsoon and monsoon seasons. During the non-monsoon season, the temperature in the water column remained constant at coastal stations. In the monsoon season, the temperature decreased by 2 to 4 ℃ from the surface to the bottom of the water column at coastal stations. In the off-shore stations, the thermocline depth was observed between 80 and 100 m during the non-monsoon season and it was shallower at 25 to 50 m, during the monsoon season (Fig. 1). During the non-monsoon season, halocline depth along the off-shore stations was observed at 80 ± 20 m, which decreased to 25 to 50 m during the monsoon season similar to the thermocline depth (Fig. 1). The influence of salinity on the water column characteristicts was also studied based on BVF from both the seasons, the influence of reduced salinity on the BVF was evident during the monsoon season (Fig. S2). The average sea surface DO concentration at the coastal and off-shore locations was 4 ± 2 mg L^-1^ during the non-monsoon and monsoon season. In the coastal stations at 30 m, DO values were consistent throughout the water column during the non-monsoon season and in the monsoon season, DO values dropped to < 1 mg L^-1^ at mid-depth and nearly reached anoxic conditions at the bottom depths. In the off-shore stations (600 m depth) DO concentration reached close to anoxic conditions at the mixed layer depth (MLD) at about 80-100 m in the non-monsoon and monsoon seasons (Fig. 1).

**Fig. 1.**
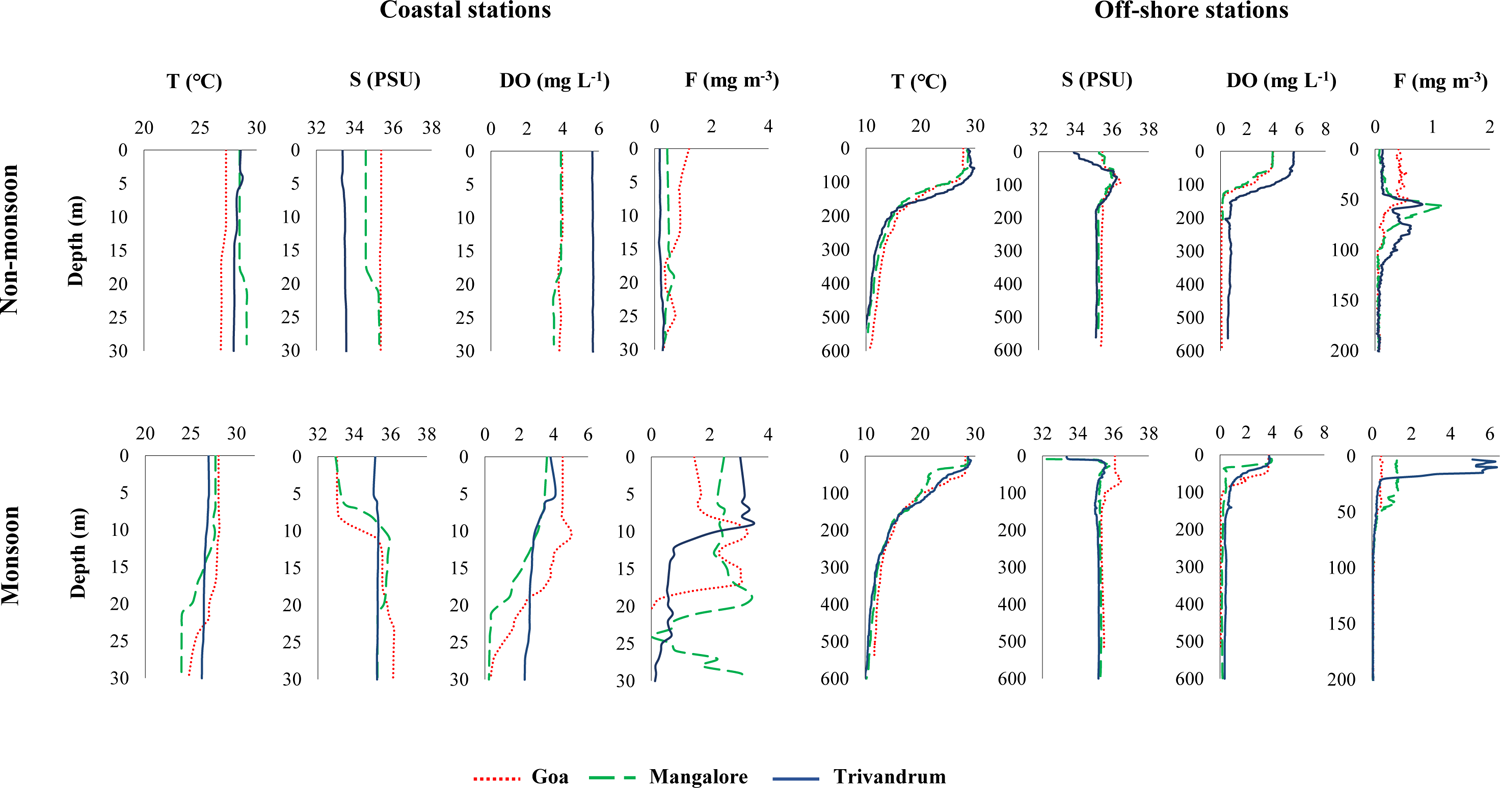
Profiles of physiochemical variables temperature (T), salinity (S), dissolved oxygen (DO) and fluorescence (F) in the water column of the coastal and off-shore stations during non-monsoon and monsoon season along the west coast of India.

Fluorescence values recorded during the non-monsoon season throughout the water column was an average ⁓ 0.4 ± 0.3 mg m^-3^ and it reached a maximum of around 1 mg m^-3^ at the C-Max depth. In the monsoon season, the average value of fluorescence in the water column was ⁓ 0.9 ± 1 mg m^-3^ and it reached a maximum of ⁓ 6 mg m^-3^ at the C-Max depth (Fig. 1). C-Max depths in the water column varied during both seasons, which were about 15 m and 55 m during the non-monsoon season in the coastal and off-shore stations respectively. In the monsoon season, C-Max in coastal and off-shore stations was observed at approximately 10 m and 30 m respectively. Higher nutrient concentration was observed in the monsoon season in both the coastal and off-shore stations. The nitrate concentration, measured at all stations during the non-monsoon season, had an average of around 2.55 ± 2.47 µM, which increased to ⁓ 17.37 ± 16.14 µM in the monsoon season (Fig. 2). The average nitrite concentration along all stations was ⁓ 0.16 ± 0.2 µM during the non-monsoon season and increased in the monsoon season to ⁓ 0.53 ± 0.85 µM. The average phosphate concentration in the water column during the non-monsoon and monsoon seasons was ⁓ 3 ± 2.8 µM and 0.92 ± 0.73 µM respectively (Fig. 2). The average silicate values detected during the non-monsoon and monsoon seasons were ⁓10.83 ± 7.30 µM and ⁓18.93 µM, respectively.

**Fig. 2.**
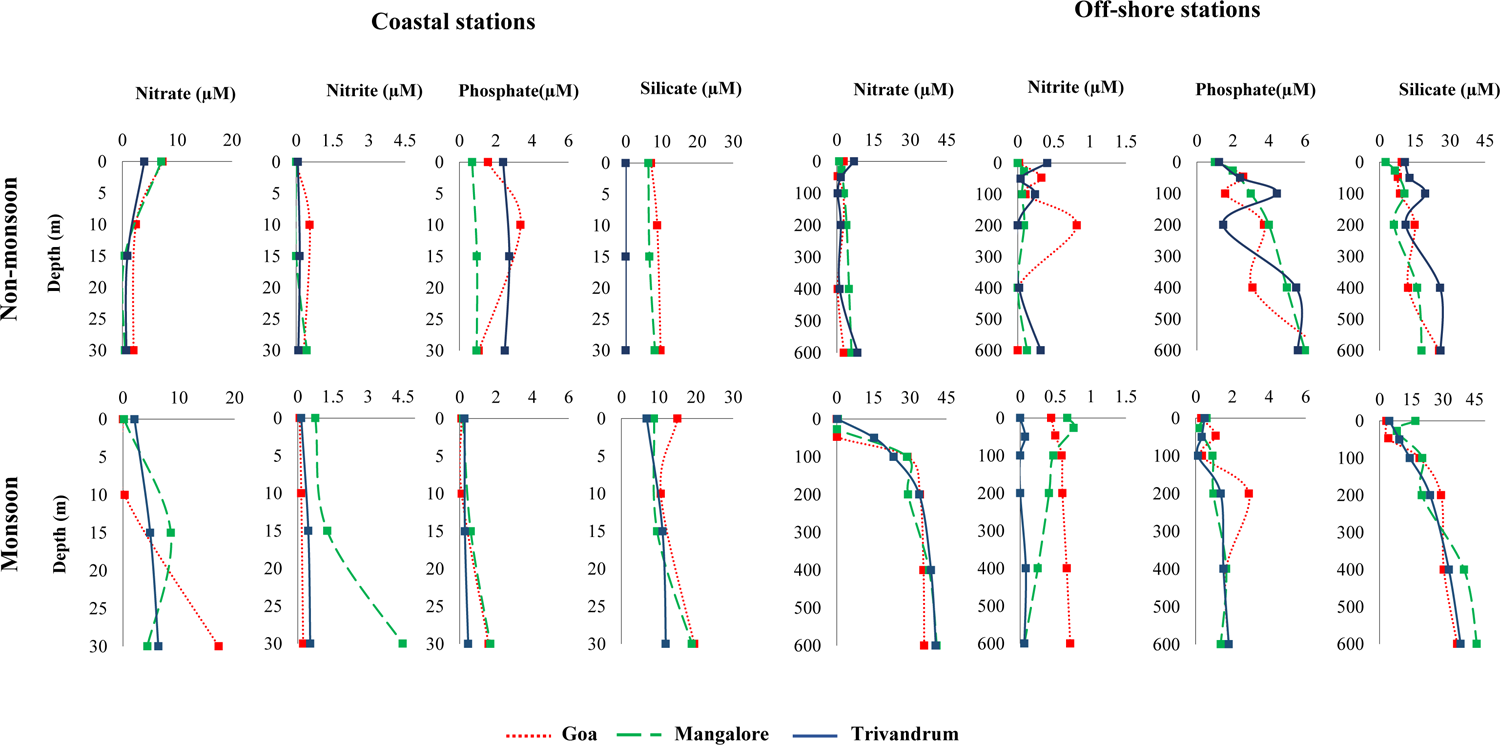
Profiles of nitrate, nitrite, phosphate and silicate in the water column of the coastal and off-shore stations during non-monsoon and monsoon season along the west coast of India.

### 3.2. Primary productivity and bacterial dynamics

Chl-a concentration estimated fluorometrically from discreet water depths was ranging from 0.002 to 0.16 mg m^-3^ in the non-monsoon and monsoon season it was estimated to be 0.003 to 1.05 mg m^-3^ (Table S1) Column integrated chl-a in the euphotic zone calculated for coastal and off-shore stations was observed to be 5.64 ± 2 mg m^-2^ in the non-monsoon and it increased to 11.42 ± 6 mg m^-2^ during the monsoon seasons (Fig. 3a). The PP in the water column during the non-monsoon season varied between 1.20 to 145.38 mg C m^-3^ d^-1^, while during the monsoon season, it ranged from 29.62 to 266.24 mg C m^-3^ d^-1^ (Table S1). The column-integrated PP in the euphotic zone was 800.31 ± 538 mg C m^-2^ during the non-monsoon season and the PP estimated during the monsoon season was higher and estimated to be 3355.60 ± 1608 mg C m^-2^ (Fig. 3b). This indicated an upwelling-induced increase in the primary productivity along the WCI during the monsoon season.

**Fig. 3.**
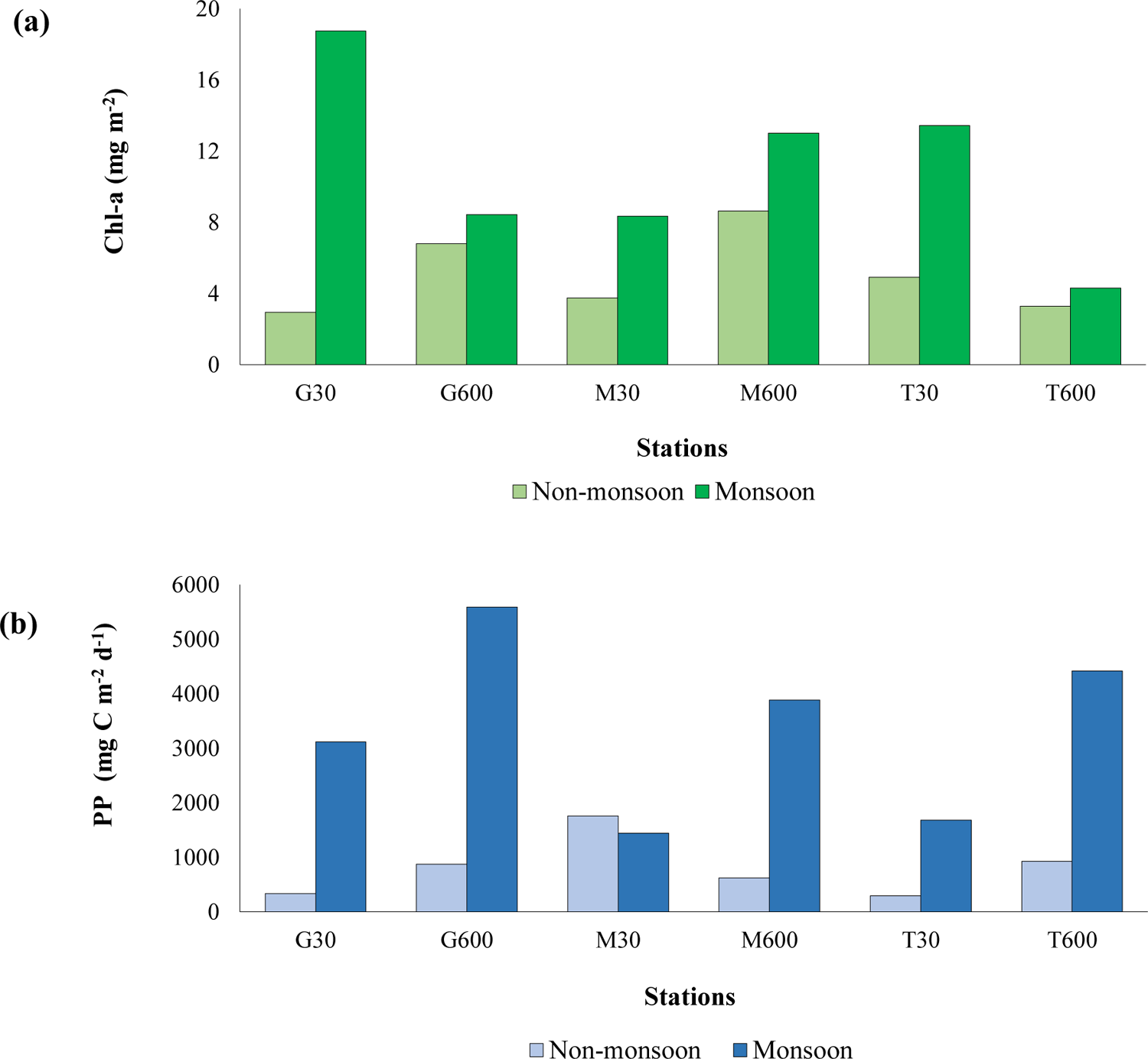
Column integrated (a) chlorophyll-a (Chl-a) concentration and (b) primary productivity (PP) in the water column of the coastal and off-shore stations during non-monsoon and monsoon season along the west coast of India.

The bacterial activity in both seasons was estimated by quantifying the bacterial abundance and productivity. The estimated BP did not vary much between the seasons which was about 10 ± 2 mg C m^-3^ d^-1^ in the coastal and off-shore stations (Table 1). Bacterial counts were determined microscopically and were expressed as bacterial carbon, which was estimated to be 209.88 ± 47 mg C m^-3^ during the non-monsoon season and it increased to an average of 1017.62 ± 132 mg C m^-3^ in monsoon season (Table 1).

**Table 1.** Average bacterial productivity (BP), bacterial abundance (BA) and bacterial carbon biomass (BC) in the water column of the coastal and off-shore stations during non-monsoon and monsoon season.

| Stations | BP<br>(mg C m<sup>-3</sup> d<sup>-1</sup>) | BA<br>(Cells × 10<sup>9</sup> L<sup>-1</sup>) | BC<br>(mg C m<sup>-3</sup>) |
| --- | --- | --- | --- |
| **Non-monsoon season** |  |  |  |
| G30 | 4.30 ± 5.19 | 10.64 ± 2.68 | 212.87 ± 53.52 |
| G600 | 2.95 ± 1.33 | 11.31 ± 4.37 | 226.11 ± 87.48 |
| M30 | 3.79 ± 2.72 | 13.65 ± 0.78 | 272.99 ± 15.65 |
| M600 | 25.92 ± 10.99 | 6.36 ± 2.06 | 127.23 ± 41.14 |
| T30 | 6.75 ± 0.93 | 10.97 ± 0.09 | 219.30 ± 1.73 |
| T600 | 17.80 ± 9.53 | 10.04 ± 4.10 | 200.78 ± 82 |
| **Monsoon season** |  |  |  |
| G30 | 13.89 ± 0.79 | 52.94 ± 14.27 | 2657.00 ± 1436.22 |
| G600 | 7.34 ± 2.26 | 57.09 ± 40.05 | 1682.00 ± 293.67 |
| M30 | 7.13 ± 0.44 | 55.51 ± 10.33 | 1923.33 ± 158.31 |
| M600 | 7.91 ± 2.04 | 41.25 ± 11.19 | 1580.00 ± 394.08 |
| T30 | 14.51 ± 2.35 | 54.45 ± 26.72 | 2226.67 ± 156.86 |
| T600 | 10.93 ± 2.27 | 44.05 ± 26.75 | 1756.50 ± 399.50 |

TPC was enumerated to estimate the viable bacterial counts in the water column from all the stations. An average of 4162.30 CFU mL^-1^ in the non-monsoon season was recorded and it increased to 11392.59 CFU mL^-1^ during the monsoon season (Fig. 4). The highest bacterial CFUs were obtained from the C-Max depths from almost all the locations sampled along the WCI (Fig. 4). An increasing trend of the bacterial carbon biomass and CFUs was in accordance with the higher primary productivity observed during the monsoon season.

**Fig. 4.**
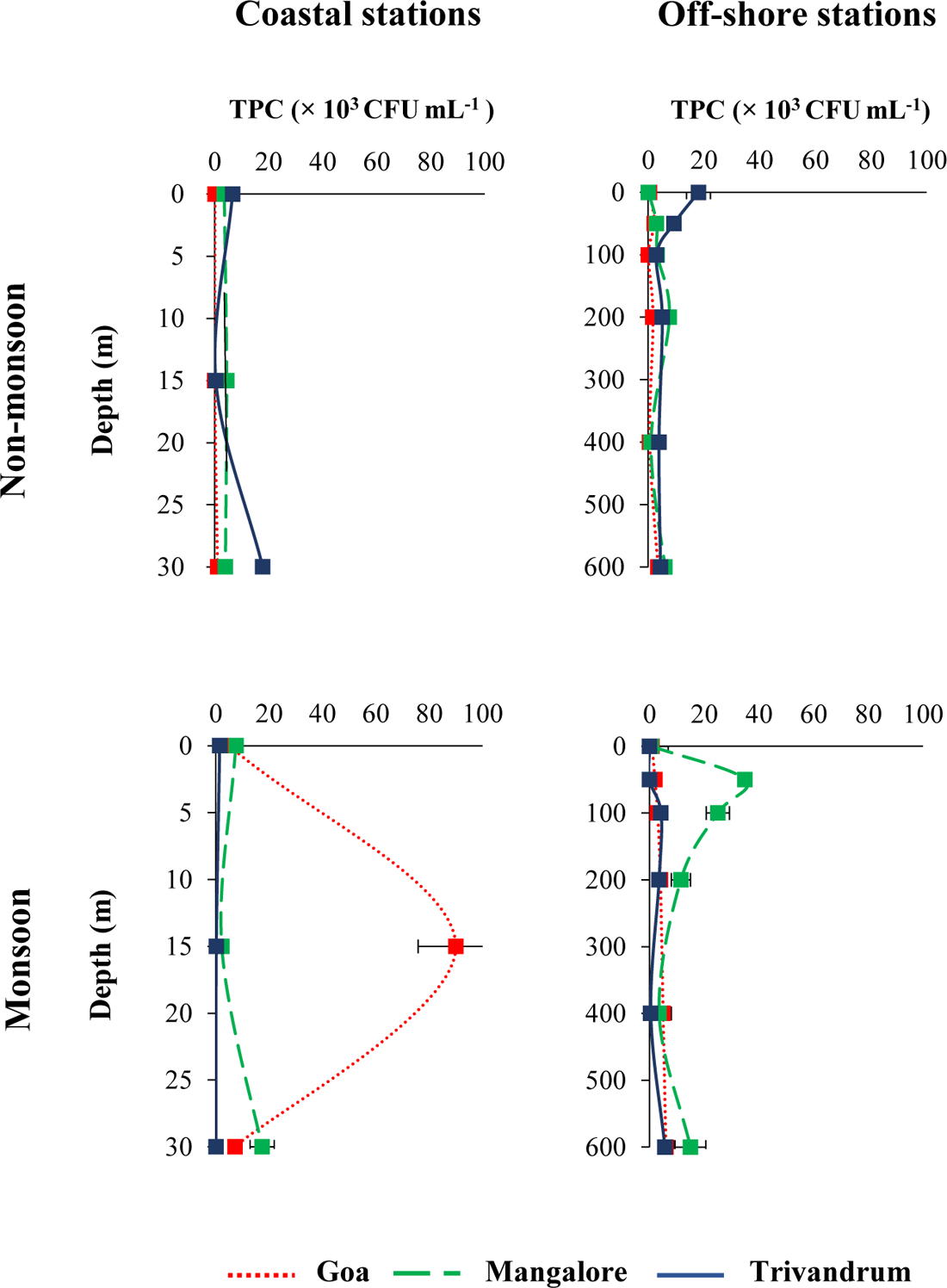
Total plate counts (TPC) of culturable bacteria in the water column of the coastal and off-shore stations during non-monsoon and monsoon season along the west coast of India.

Statistical analysis based on the ANOVA test was carried out to evaluate the significant changes in different parameters during both seasons (Table 2). Estimated environmental parameters including DO, fluorescence, nitrate, nitrite, phosphate, silicate, chl-a, PP, TPC and BC showed significant variations between seasons (p < 0.05). Correlation between environmental variables was also determined using Pearson’s correlation matrix for the physicochemical and biological variables measured from water samples during non-monsoon and monsoon seasons (Table 3). A significant correlation was observed between the different environmental variables estimated during both seasons. Temperature and DO had a significant positive correlation with fluorescence. *In situ* fluorescence concentration determined showed a significant positive correlation with PP and BC. Temperature and DO showed a significant negative correlation with nitrate and silicate. Among the nutrients, nitrate and silicate were positively correlated and Chl-a and PP showed a positive correlation with TPC and BC respectively (Table 3). These results indicate temperature has a significant impact on the fluorescence values which is a measure of the phytoplankton abundance. A significant positive correlation between the primary productivity variables and bacterial abundance and counts shows the influence of the former on the latter in the study area.

**Table 2.** Analysis of variance (ANOVA: single factor) assessing the seasonal difference in the physicochemical and biological parameters: temperature, salinity, dissolved oxygen (DO), fluorescence, nutrients, chlorophyll-a (chl-a), primary productivity (PP), bacterial productivity (BP), bacterial abundance (BA), bacterial carbon (BC) and total plates counts (TPC) between non-monsoon and monsoon season.

|  | df | F | F crit | P-value |
| --- | --- | --- | --- | --- |
| Temperature | 1.00 | 0.57 | 4.03 | 0.45 |
| Salinity | 1.00 | 0.02 | 4.03 | 0.88 |
| DO | 1.00 | 8.68 | 4.03 | 0.001** |
| Fluorescence | 1.00 | 4.58 | 4.03 | 0.04* |
| Nitrate | 1.00 | 22.34 | 4.03 | < 0.001** |
| Nitrite | 1.00 | 4.86 | 4.03 | 0.03* |
| Phosphate | 1.00 | 14.39 | 4.03 | < 0.001** |
| Silicate | 1.00 | 9.14 | 4.03 | < 0.001** |
| Chl-a | 1.00 | 5.41 | 4.04 | 0.02* |
| PP | 1.00 | 11.45 | 4.08 | < 0.001** |
| BP | 1.00 | 0.96 | 4.03 | 0.33 |
| BA | 1.00 | 70.05 | 4.03 | < 0.001** |
| BC | 1.00 | 70.05 | 4.03 | < 0.001** |
| TPC | 1.00 | 3.78 | 4.03 | 0.06* |

Df: degree of freedom, F value greater than F critical value indicates statistical significance.
\*, Significant at 0.1%., \*\*: Significant at 1%.

**Table 3.**
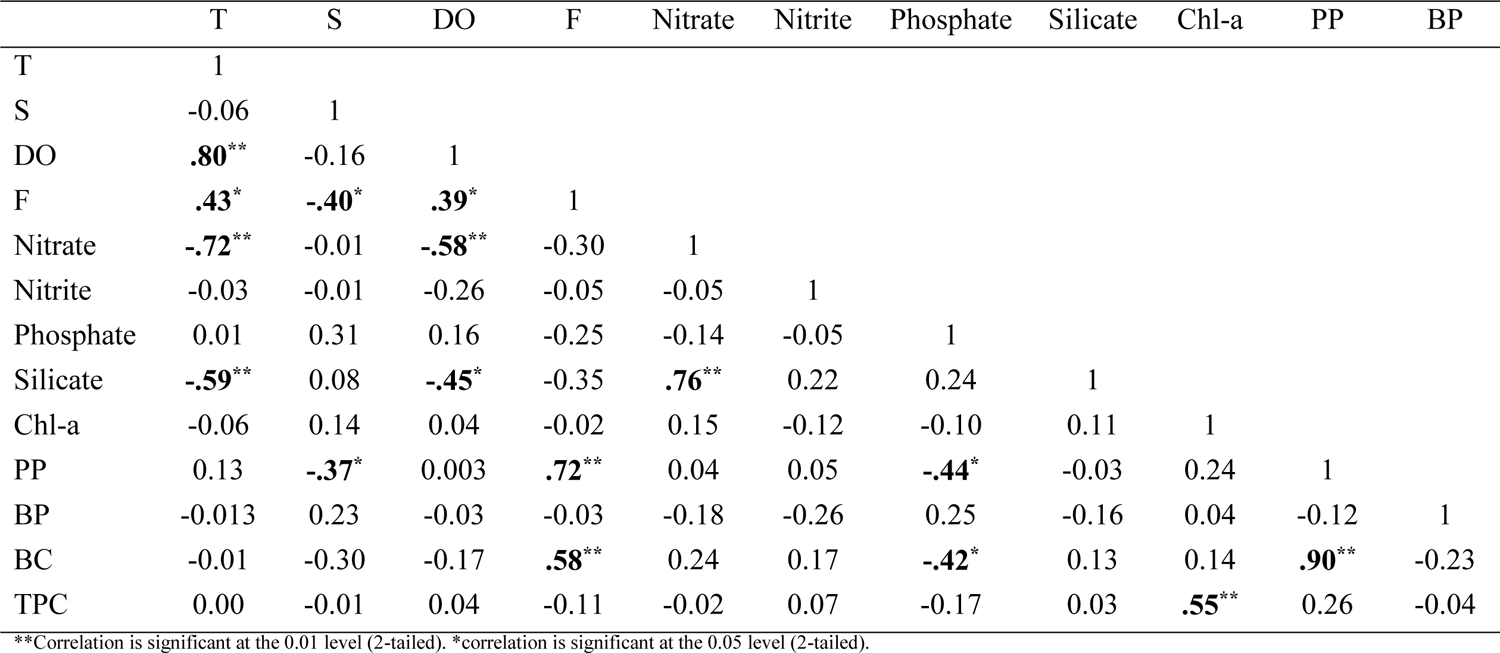
Pearson’s correlation matrix of physicochemical and biological parameters, temperature (T), salinity (S), dissolved oxygen (DO), fluorescence (F), nutrients, Chlorophyll-a (Chl-a), primary production (PP), bacterial production (BP), bacterial carbon biomass (BC) and total plates counts (TPC) during non-monsoon and monsoon season.

### 3.4. Alpha and Beta diversity of bacteria from the C-Max along the WCI

The bacterial diversity from the stations along the WCI during the non-monsoon and monsoon season was assessed from the C-Max depths based on metagenomic analysis, using next-generation Illumina sequencing techniques. Sequence statistics show that the number of reads generated was nearly similar during both seasons, however, OTUs generated after denoising the sequence reads were higher in the monsoon season (Table S2). The rarefaction curves, based on observable features, were curved toward the saturation plateau at the 97 % cut-off value, indicating that the sampling sizes were enough for the community analysis (Fig. S3). The average OTU abundance in the non-monsoon season was 646 OTUs and it increased in the monsoon season, to an average of 2286 OTUs. In comparison to other stations, higher OTU abundance was seen in the Trivandrum coastal and off-shore locations during the monsoon season (Table S2). For both non-monsoon and monsoon seasons, various alpha diversity indices for diversity and richness were calculated. Shannon, Chao1 and ACE indices were significantly higher during the monsoon season. The coastal stations showed considerably higher Shannon, Chao-1 and ACE diversity indices than the off-shore stations (Fig. 5a). Statistical analysis of the diversity indices calculated shows significant seasonal differences (Table S3).

**Fig. 5.**
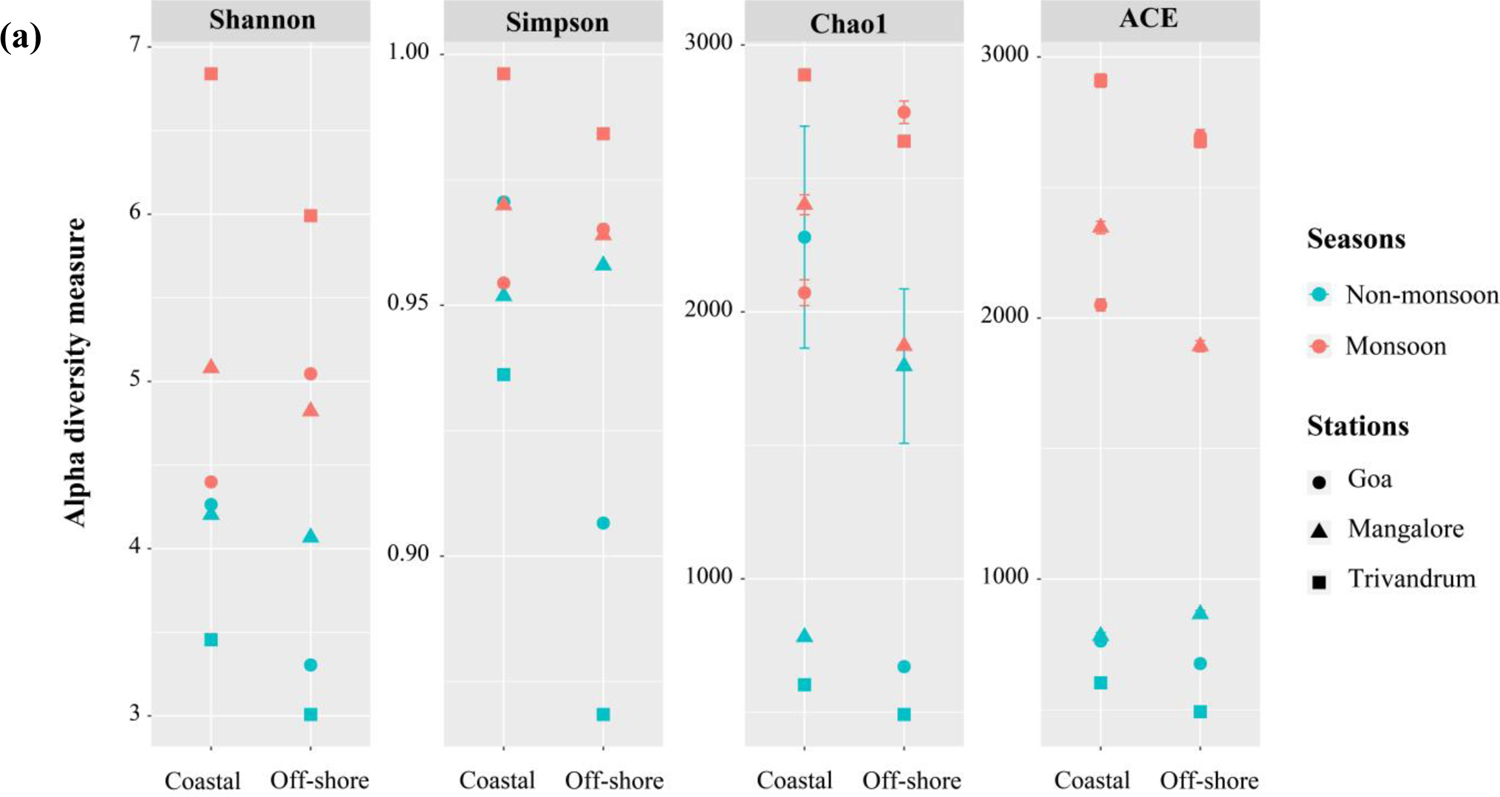
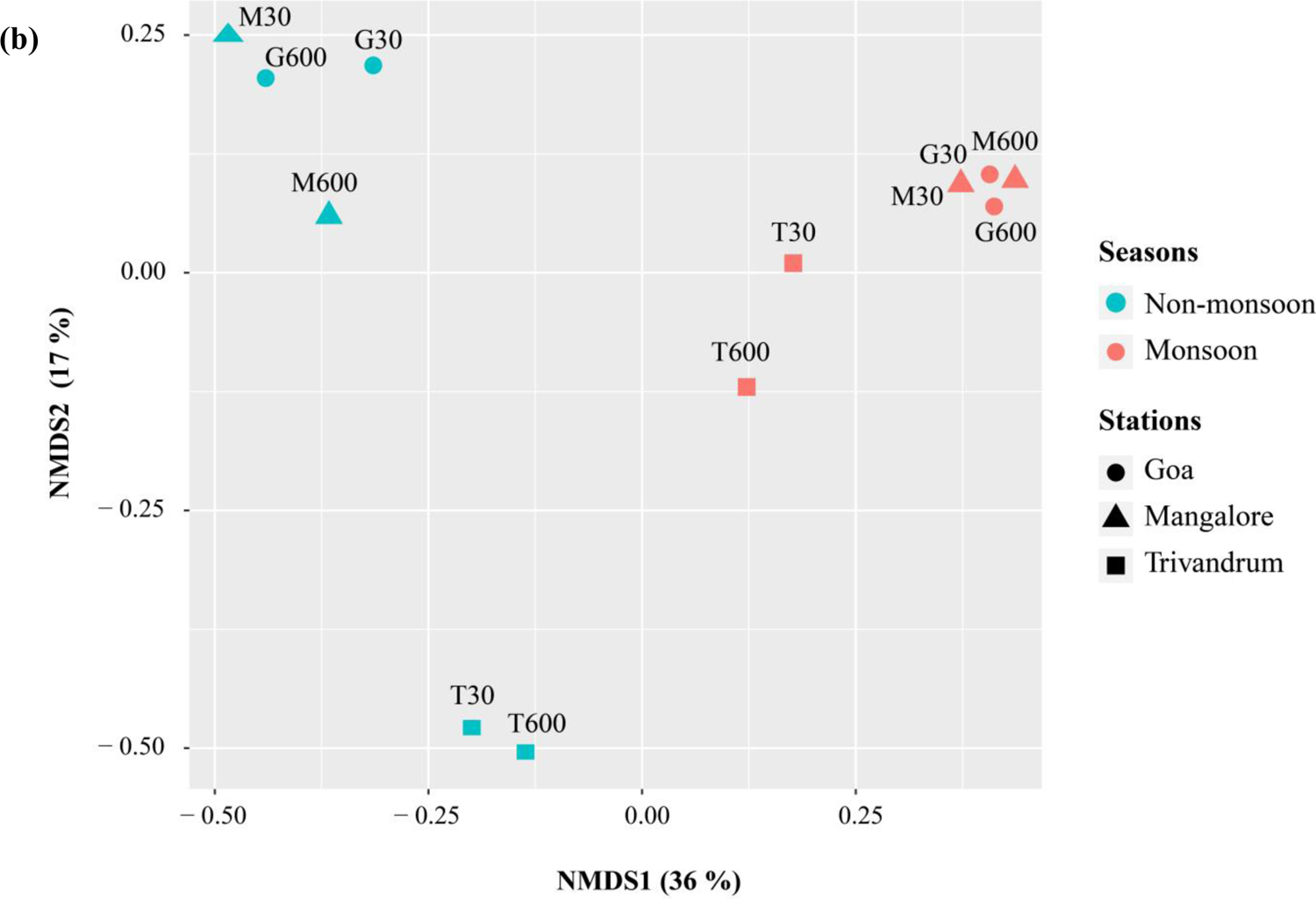
(a) Alpha diversity measure and (b) Non-metric multidimensional scaling (NMDS) plot (axis defines 2D space that allows the best spatial representation of sample distance, based on Bray–Curtis distance with stress = 0.08) depicting variations in the bacterial operational taxonomic units (OTUs) in the coastal and off-shore stations during monsoon and non-monsoon season along the west coast of India.

Beta diversity analysis of bacterial communities during non-monsoon and monsoon seasons was performed based on the NMDS method. NMDS ordination differentiates the samples into abundance-weighted community compositions based on the Bray-Curtis distance. The bacterial communities were grouped based on the seasons and locations and NMDS analysis showed that sampling locations during the non-monsoon season were divided into two apparent groups (Fig. 5b). The bacterial communities from the non-monsoon season of the Goa and Mangalore locations clustered together and a similar grouping was evident in the Trivandrum coastal and off-shore communities. A similar trend was observed in the monsoon season, where the Trivandrum station exhibited a different level of diversity than the Goa and Mangalore coastal and off-shore stations (Fig. 5b).

### 3.5. Bacterial community structure at C-Max along the WCI

Bacterial OTUs generated in Illumina sequencing were classified based on the Silva V3-V4 classifier for taxonomic profiling. For classification in this study, the sklearn classifier within QIIME was utilized, which operated on a confidence score range of 0.7 to 1. Based on taxonomic annotation, OTUs generated during both seasons belonged to 36 bacterial phyla in total, of which 26 phyla were present during the non-monsoon season and the number of phyla increased during the monsoon season to 35 (Fig. 6a). In the non-monsoon season, 18 of the 26 phyla belonged to well-defined (Based on List of Prokaryotic names with Standing in Nomenclature (LPSN), 7 of it were of candidatus status and 1 unknown or uncharacterized phyla were identified. During the monsoon season, the number of well-defined phyla increased to 22, candidatus to 12 and one unknown phyla were reported (Fig. 6a). The dominant phyla observed in the C-Max depth in the non-monsoon season were Proteobacteria (59 %) and Firmicutes (33 %), while in the monsoon season, it was Cyanobacteria (49 %) followed by Proteobacteria (32 %) (Fig. 6b). The proportion of Cyanobacteria communities increased significantly from 3 % in the non-monsoon season to 49 % during the monsoon season. Though the relative abundance of phylum Proteobacteria decreased during the monsoon season, it remained the second most dominant phyla. The relative abundance of phylum Firmicutes decreased considerably from 33 to 1 % during monsoon season. Bacteroidetes, Actinobacteria, Planctomycetes and Verrucomicrobia were also among the top 10 phyla observed during both the seasons and their abundance increased by two to three folds during the monsoon season. The temporal differences observed in dominant bacterial communities during non-monsoon and monsoon seasons were statistically significant (p < 0.001) (Fig. 6c). Among the well-defined phyla, only one phylum, Halanaerobiaeota was specific to the non-monsoon season.

**Fig. 6.**
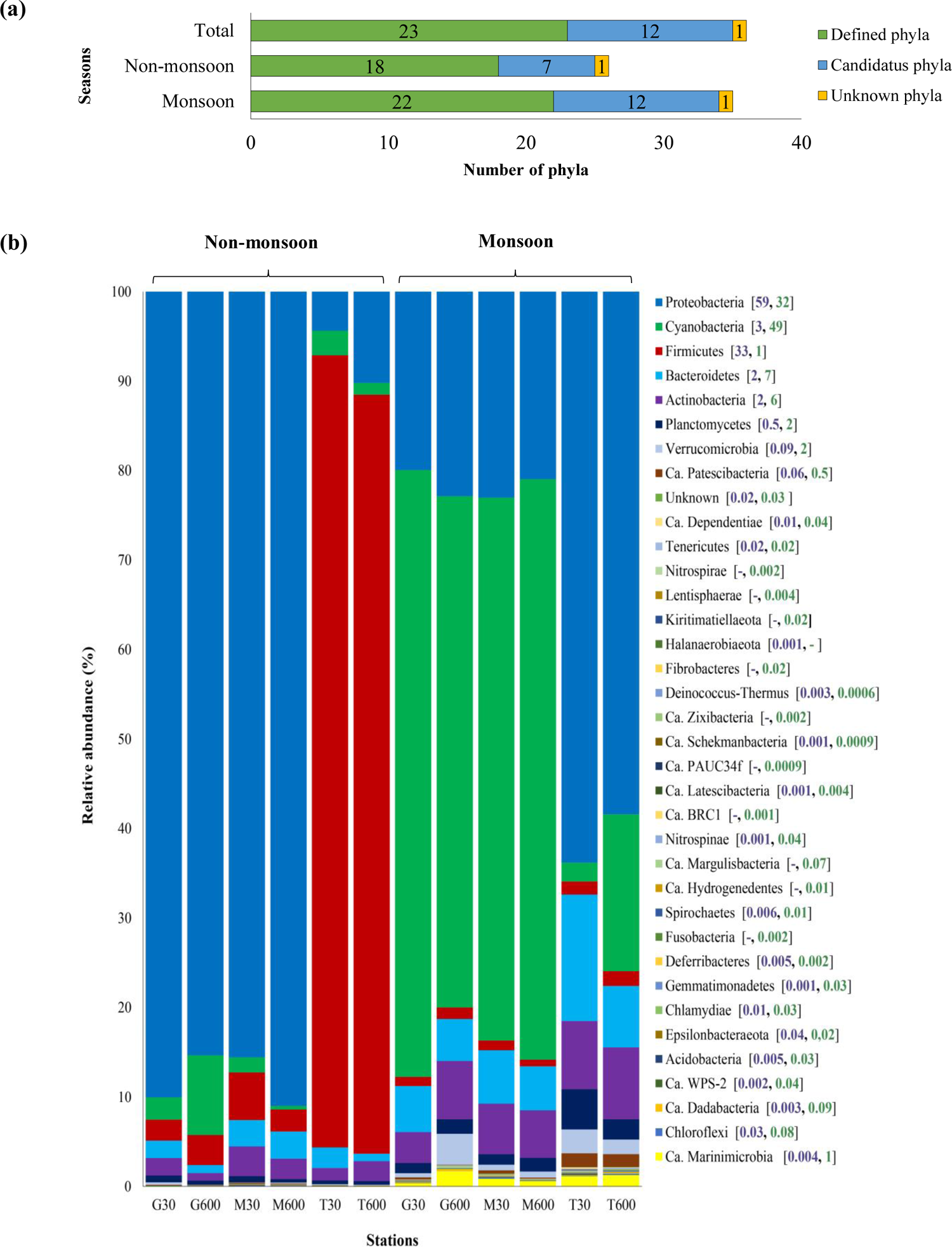
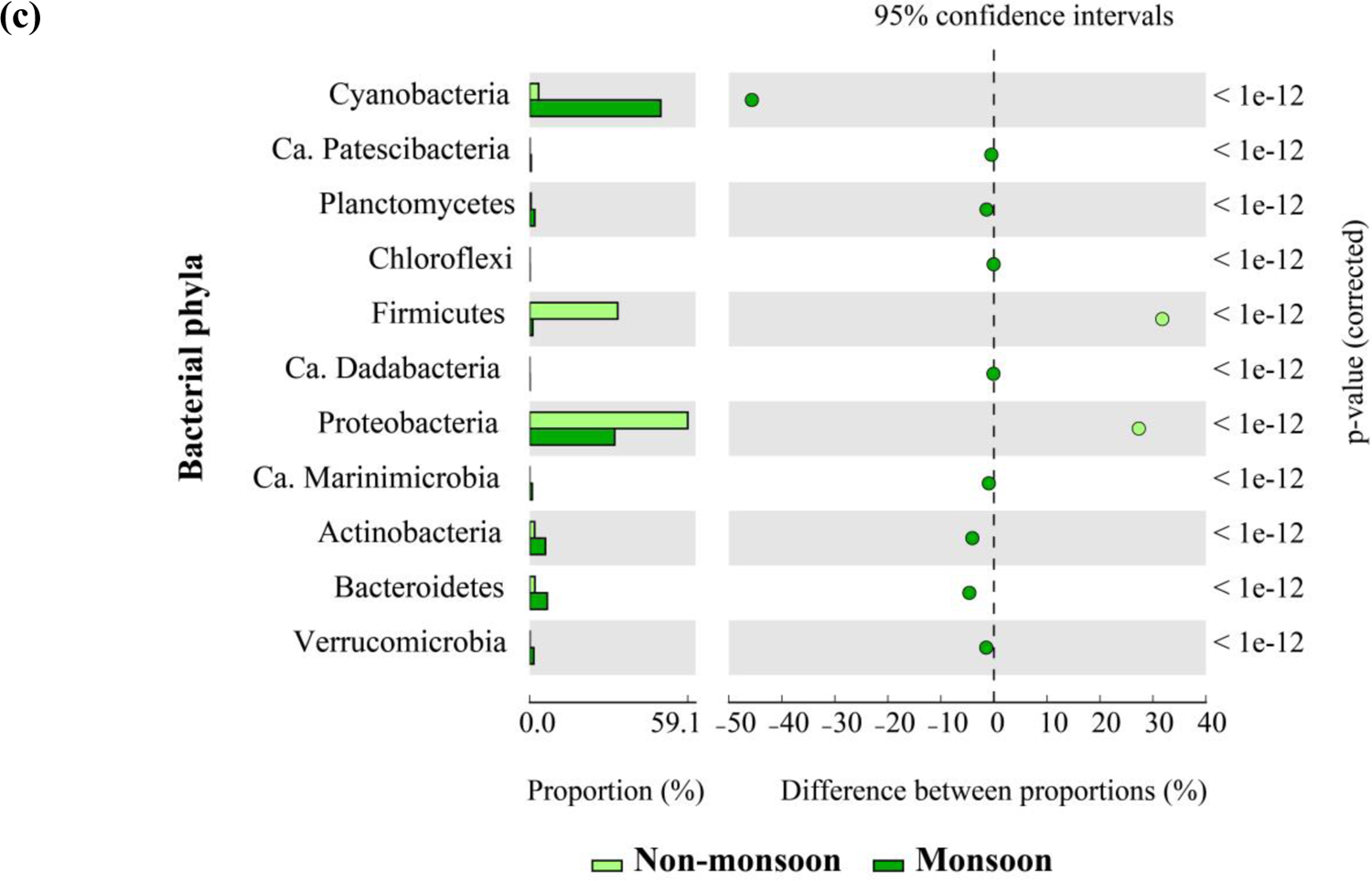
(a) Distribution of total bacterial phyla as Defined, Candidatus and Unknown taxa, (b) Relative abundance of bacterial phyla (number in parentheses indicates the relative abundance in the non-monsoon and monsoon season) and (c) Distribution of bacterial phyla showing significant statistical difference during the non-monsoon and monsoon season.

However, about 5 bacterial phyla including Nitrospirae, Lentisphaerae, Kiritimatiellaeota, Fibrobacteres and Fusobacteria were specific only to the monsoon season. There was a marginal increase in the relative abundance of six bacterial phyla, namely Nitrospinae, Spirochaetes, Gemmatimonadetes, Chlamydiae, Acidobacteria and Chloroflexi, during the monsoon season, despite their low abundance levels of less than 0.01 %. The relative abundance of a few phyla, Dianococcus-Thermus, Deferribacters and Epsilonbacteroeota showed a decreasing trend during the monsoon season. However, the diversity was higher in the monsoon with an increase in well-defined and candidatus phyla, which accounted for about 7 and 12 phyla from the non-monsoon and monsoon seasons respectively. The candidatus phyla including Ca. Patescibacteria, Ca. Dependencies, Ca. Schekmanbacteria, Ca. Latescibacteria, Ca. WPS-2, Ca. Dadabacteria, Ca. Marinimicrobia was found during both seasons. Most of the candidatus phyla that were commonly identified in both seasons also showed an increase in their relative abundance during the monsoon season. Ca. Marinimicrobia and Ca. Patescibacteria had a significantly (p < 0.001) higher contribution in diversity, ranging from 0.5 to 1 %, during the monsoon season (Fig. 6c). Candidatus phyla including Ca. Zixibacteria, Ca. PAUC34f, Ca. BRC1, Ca. Margulisbacteria and Ca. Hydrogenedentes were exclusively present during the monsoon season (Fig. 6b). These results suggest that the monsoon season had a higher diversity of both defined and Candidatus phyla (Fig. 6).

Distribution of the bacterial OTUs at the class level was also investigated using hierarchical clustering of the heatmap, based on the Euclidian similarity index for the top 5 phyla (Fig. S4a). Within the Proteobacteria phylum, the bacterial class Gammaproteobacteria was dominant during both seasons, followed by Alphaproteobacteria and Deltaproteobacteria. Although Gammaproteobacteria was dominant during both seasons, during the monsoon season, their relative abundance reduced from 48 to 17 % and there was a marginal increase in the abundance of Alphaproteobacteria and Deltaproteobacteria. The monsoon season was marked by a higher abundance of phylum Cyanobacteria, which was represented by class Melainabacteria, Oxyphotobacteria and Sericytochromatia (Fig. S4a). The diversity of phylum Firmicutes decreased during the monsoon season and distinct differences in the class level were also observed with the representation of class Bacilli and Clostridia during the non-monsoon and Erysipelotrichia and Negativicutes during the monsoon season.

The dominant classes of phyla that increased during the monsoon season are represented by Acidimicrobiia, Actinobacteria, Coriobacteriia, Nitriliruptoria, Rubrobacteria, Thermoleophilia and an unknown class of Actinobacteria; and class Bacteroidia, Ignavibacteria and Rhodothermia of phylum Bacteroidetes (Fig. S4a). The other predominant representatives were classes Planctomycetacia and Verrucomicrobiae of phylum Planctomycetes and Verrucomicrobia respectively in both the seasons (Fig. S4b). A few classes belong to the phyla Planctomycetes and Candidatus phyla Marinimicrobia (SAR406 clade), namely Ca. Patescibacteria and Ca. Margulisbacteria were exclusively present during the monsoon season. A significant difference in the distribution of the bacterial classes was observed between both seasons, with higher diversity in the monsoon season (Fig. S4b). To further investigate the community structure of bacterial populations, bacterial communities were categorized at order and family levels (Table S4). During the non-monsoon season, the most abundant bacterial family was Moraxellaceae, belonging to the order Pseudomonadales of Gammaproteobacteria. During the monsoon season, order Synechococcales of the family Cyanobiaceae belonging to phylum Cyanobacteria was the predominant autotrophic community. The most abundant heterotrophic bacterial community at order level in this season was Flavobacteriales from phylum Bacteroidetes and SAR86 clade from Gammaproteobacteria (Table S4).

### 3.6. Correlation between environmental variables and bacterial communities

The influence of environmental and primary productivity parameters on bacterial community structure at the genera level was evaluated for both the non-monsoon and monsoon seasons using CCA (Fig. 7). For the analysis, bacterial genera with a relative abundance of 0.5% and greater were plotted along with temperature, salinity, DO, fluorescence, chl-a and PP measured during both seasons. Based on the CCA ordination results, the environmental factors considered in this study accounted for 94.21 % of the variation in the community during the non-monsoon season. In this season, the bacterial genera *Cyanobium PCC-6307*, belonging to Cyanobacteria; *Bacillus* from phylum Firmicutes, showed a significant correlation with temperature, DO and chl-a. The primary productivity parameters fluorescence and PP had a major influence on the bacterial genera largely belonging to Proteobacteria, including, *Idiomarina*, *Alcanivorax*, *Alteromonas*, *Salinimonas*, *Marinobacter*, *Acinetobacter* from Gammaproteobacteria; Alphaproteobacteria including *Rhodobacteraceae_uncultured, Sphingomonadaceae_uncultured* and *Sulfitobacter* (Fig. 7a). One of the genera *Lactobacillus* from phylum Firmicutes was also positively associated with fluorescence concentration during the non-monsoon season. Environmental variables account for 87.01 % of the variation in the bacterial community during the monsoon season. PP and fluorescence were the most significant contributors to the community variation during this season (Fig. 7b). Majority of bacterial genera belonging to Proteobacteria including Ectothiorhodospiraceae_uncultured, SAR86 clade, sOM60 (NOR5) clade from class Gammaproteobacteria, SAR11 clade_Clade I, Rhodobacteraceae_uncultured, AEGEAN-169 marine group, *Euzebya* from class Alphaproteobacteria were highly correlated with the primary productivity parameters during the monsoon season. PP and fluorescence had also shown a positive influence on bacterial taxa from phylum Bacteroidetes including NS4 and NS5 marine groups and Candidatus Marinimicrobia (SAR406 clade) during the monsoon season. The bacterial genera Sva0996 marine group Ca. *Actinomarina*, from phylum Actinobacteria; OCS116 clade, *Sphingomonadaceae*_uncultured from Gammaproteobacteria and *Alteromonas* from Alphaproteobacteria, was strongly correlated to temperature (Fig. 7b).

**Fig. 7.**
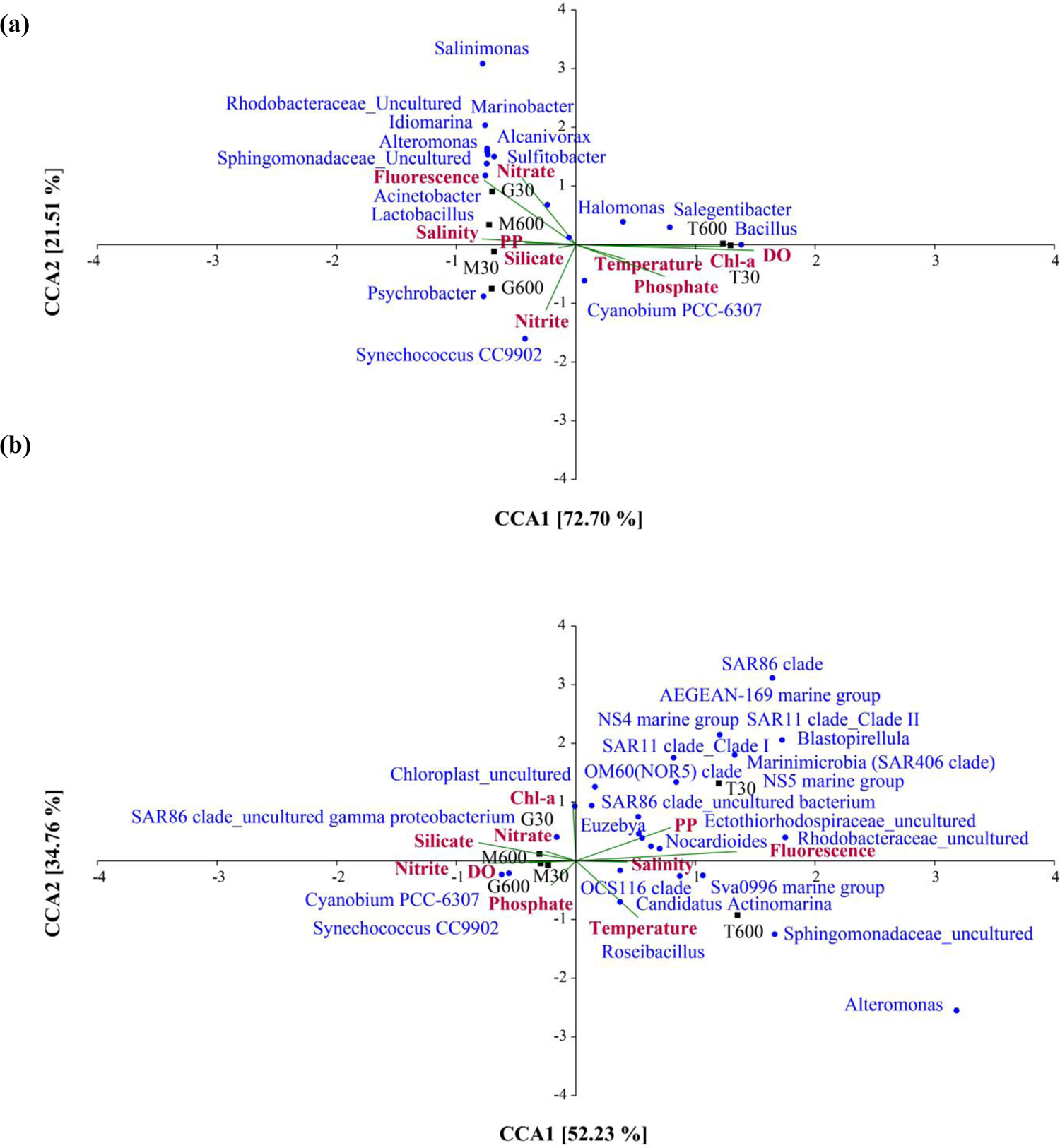
Canonical correspondence analysis (CCA) ordination plot of dominant bacterial genera and environmental variables temperature, salinity, dissolved oxygen (DO), fluorescence, chlorophyll-a (chl-a), primary productivity (PP) and nutrients during (a) non-monsoon and (b) monsoon season.

Bacterial genera *Synechococcus CC9902*, *Cyanobium PCC-6307* and *Chloroplast* from Cyanobacteria were associated with the DO and chl-a concentrations in the C-Max depth. Overall, the CCA results suggested that DO and chl-a influenced the distribution of the Cyanobacterial community. The primary productivity parameters fluorescence and PP are seen to influence the bacterial populations belonging to Proteobacteria which included a large majority of the Gammaproteobacteria and candidatus taxa. This shows that the variations in the primary productivity associated with upwelling events have a significant impact on the bacterial community composition in WCI.

### 3.7. Functional diversity at C-Max along the WCI

The functional profiles of the bacterial communities showed that the predicted pathways belonged to a variety of vital classes of functions like metabolism, processing of genetic and environmental information and pathways for cellular processes during both seasons (Fig. 8a). Significant variations in the abundance of functional genes were also evident between the non-monsoon and monsoon seasons. Further investigations into seasonal changes in the gene profiles show that, during the non-monsoon season, there was a significant increase in the abundance of metabolic pathways related to carbohydrate metabolism, lipid and amino acid metabolism, glycan biosynthesis and xenobiotic degradation. Pathways related to energy metabolism including, carbon fixation by prokaryotes, photosynthesis and membrane transport were abundant in the monsoon season (Fig. 8b and c). Functional profiling of the pathways related to the cellular processes revealed that signalling pathways, biofilm formation and pathways related to cellular prokaryotic communities were abundant during the non-monsoon season. Pathways related to cell growth and death, cell motility and prokaryotic quorum sensing were uniformly distributed in both seasons (Fig. 8c). Based on the functional gene profiles, it is evident that the non-monsoon season was associated with an increased representation of pathways related to heterotrophic metabolism, however during monsoon season autotrophic activities were higher.

**Fig. 8.**
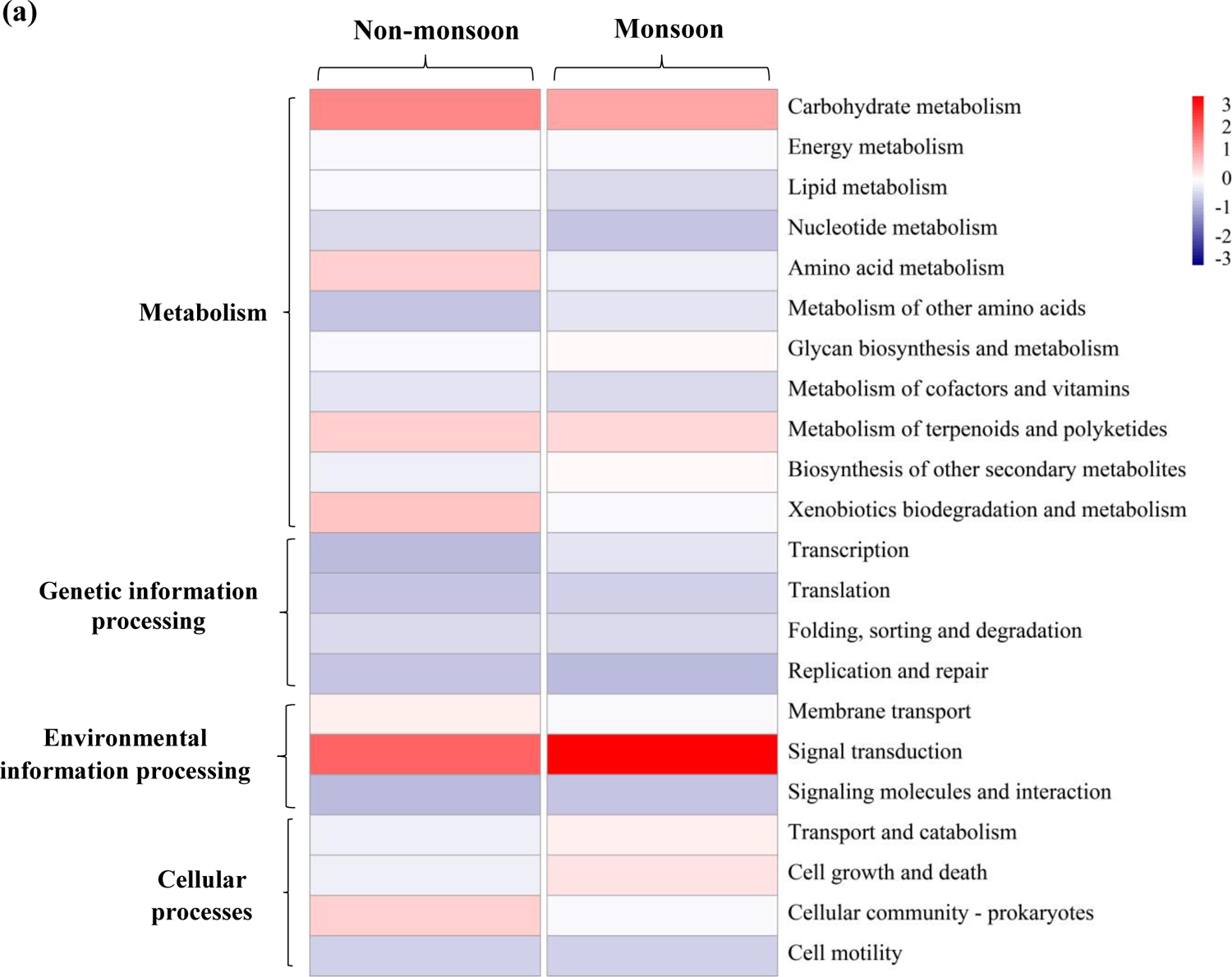
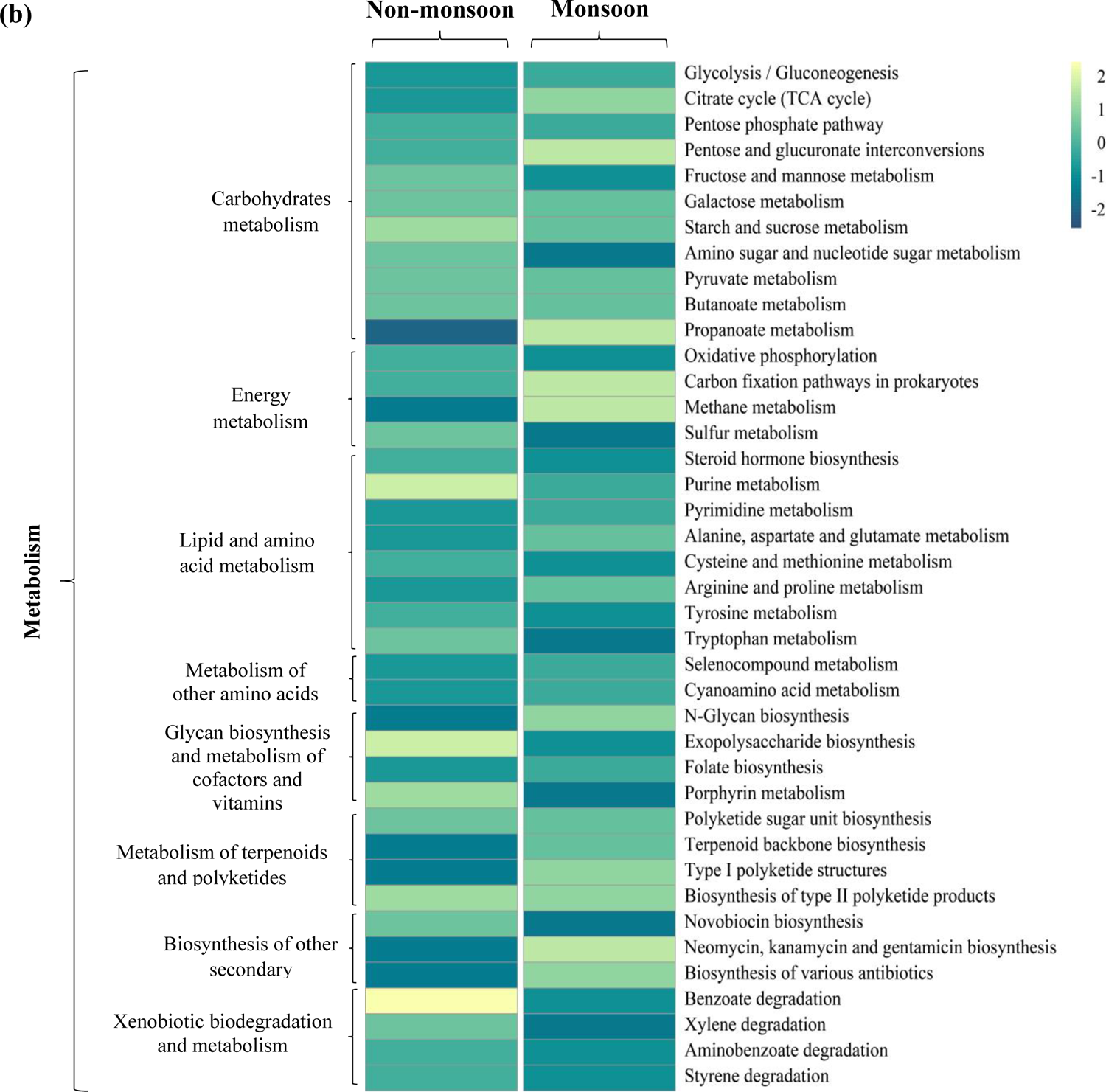
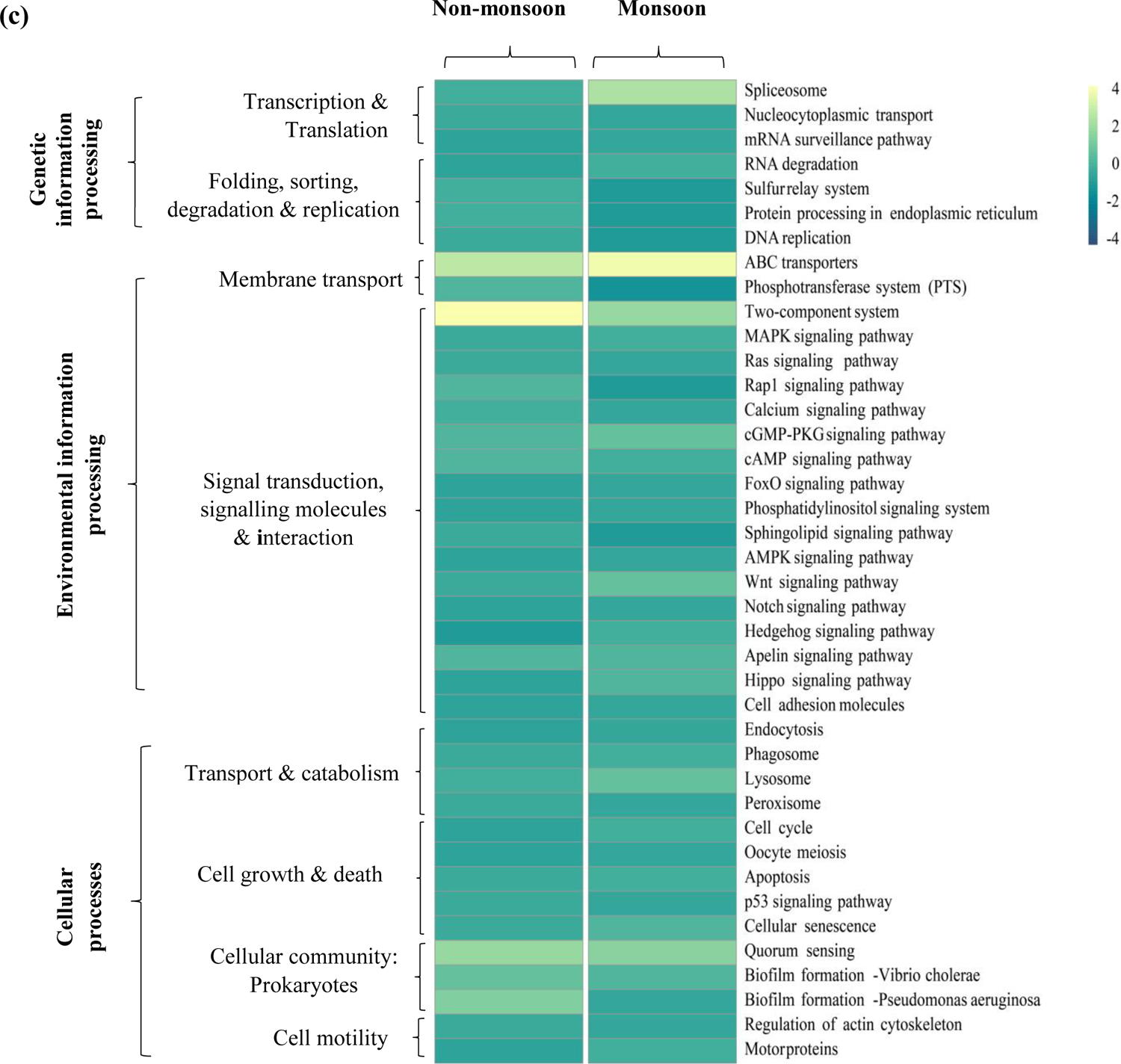
Heatmap showing functional pathways during the non-monsoon and monsoon season, (a) predicted based on KEGG database (all categories), (b) depicts pathways related to metabolism and (c) pathways related to genetic, environmental information processes and cellular processing. Carbohydrate-Active Enzymes. Front. Microbiol. 9, 1864. https://doi.org/10.3389/fmicb.2018.01864

## 4. Discussion

### 4.1. Influence of seasonal primary productivity on the bacterial dynamics

Seasonal upwelling occurs along the WCI during the southwest monsoon season which leads to increased phytoplankton productivity and accumulation of organic matter in this region (Smitha et al., 2008; Gupta et al., 2021). The availability of organic matter resulting from phytoplankton blooms during upwelling fosters the proliferation of various bacterial communities (Rocke et al., 2020). In our study, we investigated the impact of seasonal changes in primary productivity along the WCI on the bacterial dynamics to understand the differences in the bacterial communities and their functional profiles during the non-monsoon and monsoon season. The assessment of water column physicochemical parameters shows upwelling signatures, characterized by shallow MLD and increased nutrients (p < 0.01) associated with higher concentration of chl-a (p < 0.05) (Table 2) in the subsurface waters during the southwest monsoon season. This season also exhibited a significant increase in fluorescence (p < 0.05) values, indicating higher primary productivity compared to the non-monsoon season (Table 2). Throughout the monsoon season, nitrate (p < 0.01) and phosphate (p < 0.01) levels in the water column were significantly high and this causes an increase in phytoplankton productivity. *In situ* estimation of PP, as well as chl-a concentration from the study region, also indicate the presence of higher phytoplankton productivity during monsoon season. The organic matter produced by primary producers drives the microbial carbon pump and contributes to the overall energy flow and nutrient cycling (Jiao et al., 2010; Buchan et al., 2014). Increased organic load also causes a decline in the DO levels and a significant increase in nitrite concentration (P < 0.01). This was observed during the monsoon season, which may be attributed to denitrification activities of the bacterial community, which are known to occur along the WCI (Naqvi et al., 2010; Gonsalves et al., 2011). High primary productivity results in the generation of organic matter, leading to an increase in microbial activity for sustaining the food web and ecosystem functioning (Habeebrehman et al., 2008; Zingone et al., 2019; Rixen et al. 2019). Studies have suggested that high primary production is typically promoted by elevated nutrient levels associated with recently upwelled shelf waters, which are usually accompanied by a surge in bacterial abundance (Gauns et al., 2005; Lamont et al., 2014; Reji et al., 2020).

Results from this study region on the bacterial dynamics associated with changes in the primary productivity along the WCI showed an increase in viable bacterial counts during the monsoon season in the C-Max depths (Fig. 4). This suggests strong dependence of the bacterial community on organic matter derived from phytoplankton (Molina et al., 2020). Bacterial carbon estimated based on abundance value increased significantly during monsoon season and showed a positive correlation with primary productivity measurement (Table 1). These findings align with previous research conducted in the Arabian Sea, which also indicates that the higher levels of phytoplankton growth during the monsoon season have an impact on the bacterial abundance and culturable bacterial communities (Ramaiah et al., 2005; Parab et al., 2022). These studies on the bacterial dynamics in upwelling regions have also shown that a correlation between bacterial productivity and primary productivity exists, underlining the crucial role played by bacteria. However, the characterization of the relationship between seasonal variations in primary productivity levels and the structure of bacterial communities using metagenomics tools along the WCI remains unexplored.

### 4.2. Bacterial community structure and functional profiles at C-Max depths in the non-monsoon season

The intricate relationship between primary production and bacterial communities has garnered substantial attention, as bacteria are essential players in carbon cycling and organic matter recycling within the marine ecosystem. Hence, in this study, we employed metagenomic approaches to investigate the bacterial diversity associated with seasonal changes in primary production along the WCI in the C-Max depth. Our results revealed substantial variations in bacterial diversity and community structure in response to seasonal changes in primary production. Higher OTU richness and diversity (H) were observed at all the stations sampled during the monsoon season. Temporal differences in alpha diversity were also evident between the two seasons and within the locations studied along the WCI (Fig. 5).

Metagenomcc analysis of the bacaterial diversity showed that, during the non-monsoon season, only 26 of the total 36 phyla were represented and the dominance of Proteobacteria and Firmicutes followed by Bacteroidetes and Actinobacteria was observed. Within Proteobacteria, Gammaproteobacteria were the most dominant class represented by major genera such as *Psychrobacter*, *Idiomarina* and *Alcanivorax,* followed by a small proportion of Alphaproteobacteria and Deltaproteobacteria (Fig. 6). Gammaproteobacteria are reported from various marine ecosystems and the observed increase in their relative abundance can be attributed to their remarkable ability to adapt and thrive under conditions of limited organic matter availability (Cho and Giovannoni, 2004). Additionally, these bacteria possess the capacity to utilize alternative sources of energy for growth (Cho et al., 2007; Chen et al., 2021). Moreover, Gammaproteobacteria exhibit a unique capability of efficiently utilizing organic matter through the secretion of extracellular hydrolysis enzymes, enabling them to access and metabolize complex substrates present in the environment (Spring et al., 2015; Liu and Liu, 2020). Firmicutes were the second dominant phyla in the non-monsoon season represented by major genera *Bacillus* and *Lactobacillus* and their total abundance decreased significantly in the monsoon. They are known to be metabolically versatile and can adapt to different environmental conditions (Siezen and van Hylckama Vlieg, 2011; Kadnikov et al., 2020). The dominance of this group during the non-monsoon season could be due to their ability to efficiently utilize and grow in the presence of limited organic matter. These bacteria have evolved strategies to thrive well in these conditions by employing various mechanisms such as sporulation, biofilm formation and metabolic versatility (Kadnikov et al., 2020).

During the non-monsoon season, the presence of bacterial phyla such as Actinobacteria, Bacteroidetes, Planctomycetes, Verrucomicrobia and various candidatus phyla was observed to be in minor proportions (Fig. 6b). Actinobacteria and Bacteroidetes are known to encompass diverse microbial taxa with versatile metabolic capabilities (Bunse et al., 2016). However, their reduced abundance during the non-monsoon season suggests a potential limitation in their ability to thrive under nutrient-limited conditions (Ho et al., 2017). These bacterial groups may rely on higher nutrient availability, which is typically associated with periods of increased primary production, such as during the monsoon season. Furthermore, the presence of various candidate phyla in low percentages further supports the notion that these bacterial groups may struggle to flourish under conditions of limited nutrient availability. The minor representation of these candidate phyla during the non-monsoon season could suggest that they possess specialized ecological niches or physiological traits that restrict their growth and proliferation in low-nutrient environments (Yilmaz et al., 2016). This finding exhibits interesting implications regarding the adaptability of minor bacterial groups to conditions of low productivity during this season.

To understand the influence of environmental variables and primary productivity on the bacterial communities, CCA was carried out, which showed two significant correlations in the non-monsoon season. This analysis showed, that in this relatively less productive season, the dominant bacterial genera from Gammaproteobacteria and Alphaproteobacteria exhibited a significant correlation with the fluorescence concentration. The dominant bacterial genera, from the phylum Cyanobacteria, Firmicutes, Gammaproteobacteria and Bacteroidetes obtained from this season, exhibited a significant correlation with temperature, dissolved oxygen (DO) and chl-a levels (Fig. 7a). Environmental variables such as temperature influence the metabolic rates and growth of these organisms, with optimal ranges facilitating their proliferation (Mohanty et al., 2022). These metabolic capabilities of the bacterial communities enable them to thrive in diverse environments and outcompete other bacterial taxa. The physicochemical factors play an important role in shaping the structure of marine microbial communities and cause variations at spatial scales (Wang et al., 2019). Recent studies have also shown that variations in the community composition of marine bacteria as well as patterns of worldwide bacterial (16S) diversity and community structure are strongly correlated with temperature and DO (Sun et al., 2015; James et al., 2022).

Seasonal variations in primary productivity exert a significant influence on the predicted functional profiles of the bacterial communities. The functional diversity of the bacterial assemblages is known to vary, with evidence suggesting that the upwelling seasons may exhibit higher levels of functional diversity than other periods. This indicates that during the upwelling season, the metabolism of microbial communities is particularly active in breaking down complex organic compounds such as carbohydrates, proteins and polysaccharides (Caroppo et al., 2022). However, our previous study based on culturable bacterial populations found that functional activities in the water column during the monsoon season were reduced comparatively than in the non-monsoon season (Parab et al., 2022). Similarly, this study based on metagenomics approaches and predictive gene profiling showed that the heterotrophic metabolic activities were higher during the non-monsoon season (Fig. 8). The abundance of genes involved in membrane transport, carbohydrate metabolism, xenobiotic degradation and amino acid metabolism shows that metabolic exchange and nutrient transformation through extracellular heterotrophic activities were significantly higher during non-monsoon could be due to limited nutrient availability. Genes related to two-component signal transduction systems were also significantly higher during the non-monsoon season when productivity was low. Studies have also reported that bacteria can sense and adapt to diverse environments, stressors and growth conditions and also respond to stimuli like low nutrient and quorum signals (Mitrophanov and Groisman, 2008; Sebastián et al., 2013; Prüß, 2017). Limited nutrient availability can indeed increase heterotrophic activities in bacterial communities. When nutrients are scarce, heterotrophic bacteria like Alphaproteobacteria SAR11 and Flavobacteria are known to adapt by maximizing their resource utilization and metabolic efficiency (Sebastián et al., 2013, 2016). Bacteria may exhibit increased enzymatic activity and cellular activity such as quorum sensing and biofilm formation allowing them to efficiently break down available organic matter and obtain essential nutrients for growth (Lever et al., 2015 Prüß, 2017), which was also evident from our study (Fig. 8c). An increase in functional genes of pathways associated with heterotrophic metabolic activities were found to be more prevalent during the non-monsoon season. This can be linked to the contribution of the dominant Gammaproteobacteria and Firmicutes communities. Their higher representation during the non-monsoon season suggests their active involvement in heterotrophic metabolic activities during the non-monsoon season.

### 4.3. Bacterial community structure and functional profiles at C-Max depths in the monsoon season

During the monsoon season, the bacterial diversity was higher with representatives from 35 phyla belonging to various defined and candidatus phyla. The dominant bacterial groups during this season were from the autotrophic bacterial phyla, the Cyanobacteria which increased by three folds during the monsoon season. The significant increase in the abundance of Cyanobacteria from 3 % during the non-monsoon season to 49 % during the monsoon season implies that this group of bacteria may have a competitive advantage over other bacterial groups during the upwelling season. Cyanobacteria are known to form blooms in nutrient-rich waters and thus increase in their abundance as compared to non-monsoon season (James et al., 2022). During both seasons, *Synechococcus* and *Cyanobium* were the dominant genera of Cyanobacteria observed. *Synechococcus* is considered a significant Cyanobacterial genus due to its important ecological role in oceanic upwelling regions, specifically in carbon cycling and trophic networks (Dore et al., 2022). Though there was an increase in the abundance and diversity of bacterial communities in the monsoon season; there was a decrease in the abundance (from 59 to 32 %) and community shift in the diversity of the most prevalent heterotrophic bacterial group Proteobacteria. The decrease in the abundance of this major group was obtained largely due to the reduction in the class Gammaproteobacteria. There was also a change in the diversity within this group which was represented by *Alteromonas*, SAR86 clade and OM60 (NOR5) in this season. These genera could be of ecological importance in the upwelling regions as this class of bacteria is reported from regions of high primary productivity (González et al., 2000; Alonso-Sáez et al., 2007). Within this phyla, Alphaproteobacteria and Deltaproteobacteria abundance also increased significantly during the monsoon season (Fig. S4). Alphaproteobacteria have been identified as particle-degrading microorganisms, which suggests their potential significance as key contributors to the biogeochemical cycling of carbon and other nutrients during upwelling (Bachmann et al., 2018). Deltaproteobacterial abundance has also been shown to be higher in upwelling regions. They exchange nutrients with Alphaproteobacterial and other bacterial communities through mutualism interactions, providing inorganic compounds in exchange for organic carbon compounds and thus play important roles in linking the nutrient cycles (Sheik et al., 2013; Aldunate et al., 2018; Sun et al., 2022).

The abundance of Bacteroidetes, another prominent phylum observed in this study and increases significantly during the monsoon season and has been commonly found associated in upwelling systems with algal blooms (Alonso et al., 2006; Schattenhofer et al., 2009). They are also important players in the breakdown of particulate organic materials in the water column (Kirchman, 2002; Pinhassi et al., 2004; Bergen et al., 2015). It was found that Actinobacteria is a prevalent and widely distributed phylum and its abundance increased from 2 % during the non-monsoon season to 6 % during the monsoon season. The diversity of these phyla is reported from C-Max depths in other upwelling regions (Morris et al., 2012; Montes et al., 2020); which are possibly increased in numbers due to the accumulation of phytoplankton biomass at these depths. Other dominant phyla observed were Verrucomicrobia and Planctomycetes; known to be frequently prevalent and numerous in marine bacterial assemblages (Freitas et al., 2012). Thus, during upwelling events, Cyanobacteria and

Proteobacteria tend to dominate, indicating their ecological significance, while a significant increase in other bacterial groups during upwelling, such as Actinobacteria, Bacteroidetes, Verrucomicrobia and Planctomycetes contribute to organic matter recycling (Rocke et al., 2020; Sun et al., 2020; Reji et al., 2020). In addition to the well-defined phyla observed in the study region, the presence of various candidatus phyla was also detected. Notably, during the monsoon season characterized by high primary productivity, candidatus phyla including Ca. Marinimicrobia, Ca. Patescibacteria and Ca. Dadabacteria exhibited a significant increase in their relative abundance. Their potential roles in the decomposition and utilization of the abundant organic matter generated during this period highlight their ecological significance in nutrient cycling and carbon metabolism within the study region (Mestre et al., 2018; Frank et al., 2016). Further investigaions into the functional potential and ecological contributions of these candidatus phyla are warranted to elucidate their specific roles in microbial community dynamics and ecosystem functioning during the monsoon season.

Based on CCA analysis, an association of Cyanobacterial genera including *Cyanobium and Synecoccoccus* with temperature, DO and chl-a concentrations during monsoon season was evident, this highlights their sensitivity to environmental conditions (Fig. 7b). Dissolved oxygen availability is crucial for the survival of aerobic bacteria, including Cyanobacteria (Berg et al., 2022). The structure of the bacterial assemblages in the marine environment is subject to temporal and regional change due to oceanographic phenomena like upwelling (Alonso-Gutiérrez et al., 2009) and also due to environmental factors induced by upwelling including physicochemical changes and organic matter generation. Physicochemical factors such as temperature and DO availability play a crucial role in shaping the diversity, abundance and activity of bacterial communities in the marine ecosystem, particularly in upwelling regions (Sun et al., 2022). A positive correlation of fluorescence and PP with the majority of bacterial communities including Gammaproteobacteria SAR66, OM60 (NOR5), Bacteroidetes NS4 and NS5 marine groups, Actinobacterial genera Sva0996 marine groups, Ca. *Actinomarina*, Ca. *Marinimicrobia* and other candidatus groups during the monsoon season were observed. These bacterial groups have previously been shown to be associated with high-productivity environments and their presence may reflect an increase in nutrient availability due to upwelling (Bachmann et al., 2018; Reji et al., 2020; James et al., 2022). This surge in the abundance of candidatus phyla can be attributed to their adaptation to thrive in environments rich in organic matter derived from primary productivity (Mestre et al., 2018; Frank et al., 2016). The enhanced availability of nutrients and labile carbon sources during the productive monsoon season likely stimulates the growth and metabolic activities of these bacterial phyla (Hug et al., 2015; Reji et al., 2020; Graham and Tully, 2021). Seasonal variations in primary productivity have been recognized as a crucial driver of microbial community composition and abundance. As primary productivity variations occur, bacterial communities respond by adjusting their composition and functional capacities to adapt to changing environmental conditions (Nguyen et al., 2020). The relationship between primary productivity-derived organic matter and bacterial diversity existed at both local and global scales (Phoma et al., 2021). Understanding the influence of seasonal primary productivity variations on increased bacterial diversity is essential for unraveling the intricate relationships between primary producers and bacterial populations, as well as their roles in nutrient cycling and ecosystem functioning. The correlation of bacterial communities with fluorescence and PP indicated that changes in these parameters in the seawater column could impact the abundance and composition of bacterial community variations. Studies had shown that the interactions between phytoplankton and accompanying bacteria could have important effects on both communities (Jasti et al., 2005; Rooney-Varga et al., 2005).

Predictive functional profiling showed autotrophic activities including, prokaryotic carbon fixation activities and cellular catabolism increased during the monsoon season. Autotrophic activities, as well as the abundance of prokaryotic carbon fixation pathways, could be due to the significant increase in Cyanobacterial populations during the upwelling season, which may have impacted the results of our metagenomics comparisons. Studies suggest that the differences in functional profiles observed between different seasons using metagenomics approaches may be influenced by the high abundance of genes associated with photosynthesis from Cyanobacteria during the upwelling season (Ward et al., 2017; Matsumoto et al., 2021). Upwelling circumstances are linked to an increase in energy metabolic processes (Coelho-Souza et al., 2012; Gregoracci et al., 2015; Parvathi et al., 2019), such as prokaryotic carbon fixation, nitrogen metabolism and methane metabolism, which was also evident from our study (Fig.8a). Understanding the metabolic interactions between primary producers, such as photoautotrophic species like phytoplanktons and Cyanobacteria and heterotrophic organisms, such as bacteria and archaea, is essential for understanding how ecosystems work (Caroppo et al., 2022). The results of our study indicate that upwelling has the potential to increase the metabolic capacity of the majority of the bacterial community, particularly in the areas of prokaryotic carbon fixation and various cellular catabolic processes (Fig. 8b). Compounds such as dissolved organic matter can enhance the growth of bacterial species such as Roseobacter, Oceanospirillales and Flavobacteriales (Osterholz et al., 2016), leading to alterations in the microbial community and functional diversity. Overall during the monsoon season, the dominant pathways related to autotrophic carbon fixation and cellular processing were abundant. The dominance of these pathways during monsoonal upwelling season indicates that bacterial communities are adapting to the changes in nutrient availability and environmental conditions during monsoon season and are prioritizing biomass production and maintenance of cellular metabolism as key functions. This observation underscores the significant impact of primary productivity variations, particularly those associated with upwelling events, on the composition of bacterial communities within the WCI region.

## 5. Conclusion

This study presents a detailed account of the bacterial community structure in C-Max depths during the non-monsoon and monsoon seasons along WCI. This study has improved the understanding of microbial functioning along the WCI, an important upwelling zone and a significant carbon sequestration region. Major implications from this study is based on the observed taxonomic and functional diversity patterns, combined with physicochemical factors, due to primary productivity variations and relationships among bacterial communities. This study using Next-generation sequencing methods, has provided a deeper understanding of the community structure and diversity associated with seasonal changes in primary productivity, along the WCI. The level of taxonomic resolution provided by Illumina sequencing allowed us to build upon previous research and uncover that, seasonal gradients of the bacterial taxonomic makeup, revealing spatiotemporal variations at genera level, that was previously unexplored from this region. Additionally, Illumina sequencing has enabled us to understand the presence of candidatus phylotypes which are metabolically active, which are not characterised by conventional culture-based methods. The seasonal transitions in WCI, and the dynamics of the bacterial community composition are most likely the outcome of increased primary productivity and environmental factors. Further studies are required to understand the basis of microbial mechanisms connected to organic matter recycling and export to the deeper depths of this region. This study has improved the understanding of the seasonal changes in microbial community structure along the WCI in the Arabian Sea.

## Data availability

Data will be made available on request. The metagenomics data generated in this study have been deposited in the National Centre for Biotechnology Information (NCBI) Sequence Read Archive (SRA) under the bioproject number PRJNA873935. The data can be accessed from the following link: [https://www.ncbi.nlm.nih.gov/bioproject/PRJNA873935].

## Acknowledgments

The authors wish to extend their appreciation to the captain, crew and ship cell staff, as well as our colleagues on board RV Sindhu Sankalp, for their constant support during the SSK-125 and SSK-131 cruises. Ashutosh S Parab expresses gratitude to UGC, India, for providing the research fellowship, Grant No. F.16-6 (Dec.2016)/2017 (NET). This is NIO contribution number xxxx.

Appendix A. Supplementary data

Appendix B. Supplementary file

