## Supplemental data for "Variations in bacterial community at Chlorophyll Maximum (C-Max) depths along the west coast of India due to seasonal changes in the primary productivity"

**Contents**

**Fig. S1.** Map illustrating the study area and sampling stations off Goa, Mangalore and Trivandrum along the west coast of India……………………………………………………...3

**Fig. S2.** Depth profiles of density and Brunt-Vaisala frequency (BVF) in the water column of the coastal and off-shore stations during non-monsoon and monsoon season……………………………………………………………………………………….....4

**Fig. S3.** Rarefactions curve at various sequencing depths for (a) non-monsoon and monsoon season………………………………………………………………………………………….5

**Fig. S4.** Hierarchal clustering heatmap based on the Euclidian similarity index, showing the relative abundance of bacteria at class level (a) from top five phyla and (b) from other than top five phyla during non-monsoon and monsoon season……………………………………….....6

**Table S1** Depth wise values of chlorophyll-a (Chl-a), primary productivity (PP), bacterial productivity (BP), bacterial abundance (BA), bacterial carbon (BC), and total plate count (TPC) in the water column of the coastal and off-shore stations during non-monsoon (NMS) and monsoon season (MS)…………………………………………………………………………7

**Table S2** Read statistics of the amplicon sequence variants (ASVs) and operational taxonomic units (OTUs) generated……………………………………………………………………….11

**Table S3** Analysis of variance (ANOVA: single factor) assessing the seasonal difference in the bacterial OTUs generated and diversity indices between non-monsoon and monsoon season………………...………………………………………………………………………12

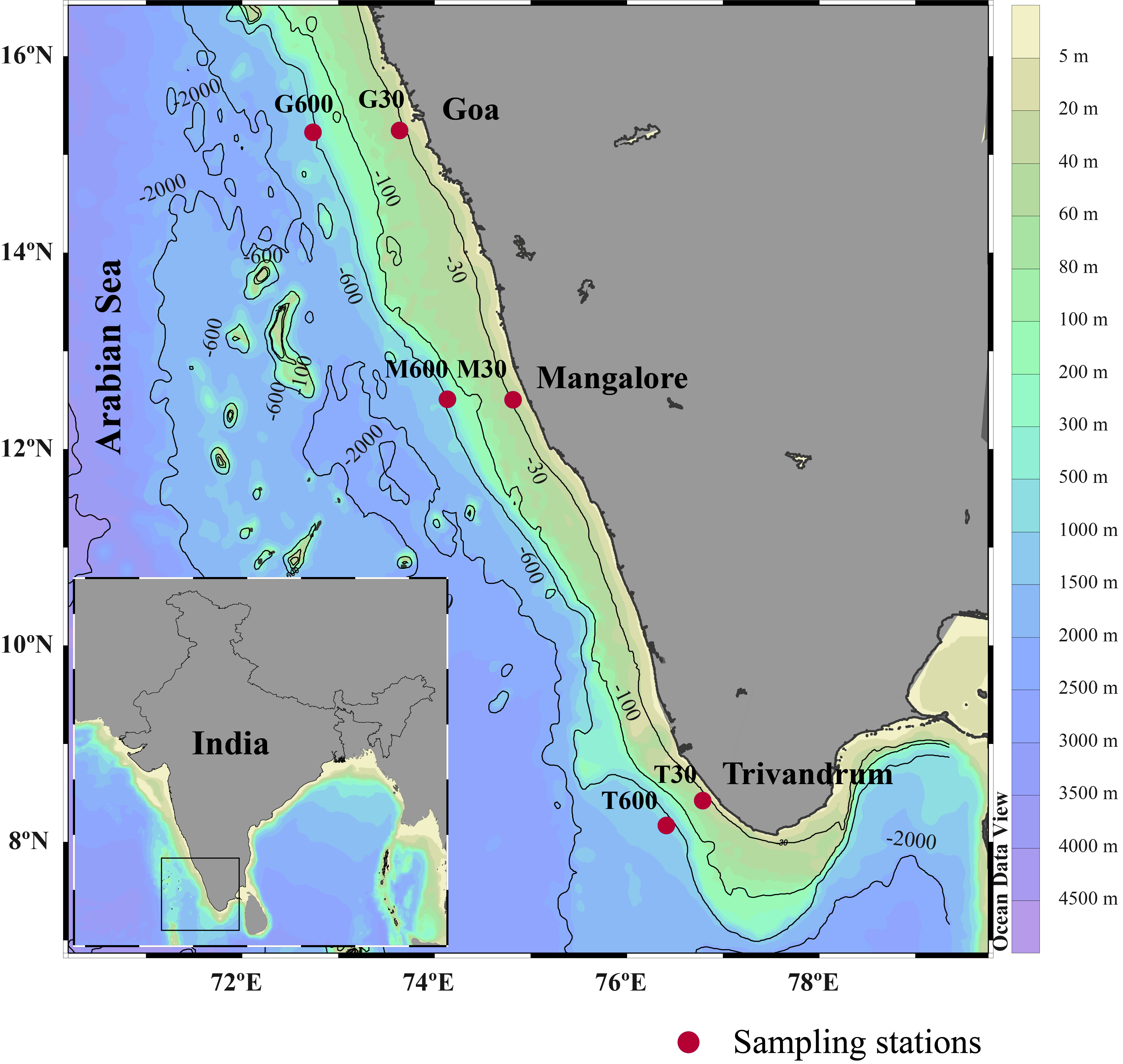

Fig. S1. Map illustrating the study area and sampling stations off Goa, Mangalore and Trivandrum along the west coast of India.

**Off-shore stations**

**Coastal stations**

| Density (kg m-3) | G600 |
| --- | --- |
| 1022.83 | 3 |
| 1022.83 | 4 |
| 1022.83 | 5 |
| 1022.84 | 6 |
| 1022.84 | 7 |
| 1022.84 | 8 |
| 1022.85 | 9 |
| 1022.85 | 10 |
| 1022.86 | 11 |
| 1022.86 | 12 |
| 1022.86 | 13 |
| 1022.87 | 14 |
| 1022.87 | 15 |
| 1022.88 | 16 |
| 1022.88 | 17 |
| 1022.89 | 18 |
| 1022.89 | 19 |
| 1022.89 | 20 |
| 1022.9 | 21 |
| 1022.9 | 22 |
| 1022.91 | 23 |
| 1022.91 | 24 |
| 1022.92 | 25 |
| 1022.92 | 26 |
| 1022.92 | 27 |
| 1022.93 | 28 |
| 1022.93 | 29 |
| 1022.94 | 30 |
| 1022.94 | 31 |
| 1022.95 | 32 |
| 1022.96 | 33 |
| 1022.97 | 34 |
| 1023 | 35 |
| 1023.05 | 36 |
| 1023.09 | 37 |
| 1023.11 | 38 |
| 1023.12 | 39 |
| 1023.13 | 40 |
| 1023.13 | 41 |
| 1023.14 | 42 |
| 1023.15 | 43 |
| 1023.16 | 44 |
| 1023.17 | 45 |
| 1023.18 | 46 |
| 1023.18 | 47 |
| 1023.2 | 48 |
| 1023.21 | 49 |
| 1023.21 | 50 |
| 1023.22 | 51 |
| 1023.24 | 52 |
| 1023.26 | 53 |
| 1023.27 | 54 |
| 1023.28 | 55 |
| 1023.3 | 56 |
| 1023.31 | 57 |
| 1023.32 | 58 |
| 1023.34 | 59 |
| 1023.37 | 60 |
| 1023.38 | 61 |
| 1023.42 | 62 |
| 1023.41 | 63 |
| 1023.45 | 64 |
| 1023.47 | 65 |
| 1023.48 | 66 |
| 1023.51 | 67 |
| 1023.55 | 68 |
| 1023.61 | 69 |
| 1023.65 | 70 |
| 1023.68 | 71 |
| 1023.73 | 72 |
| 1023.75 | 73 |
| 1023.77 | 74 |
| 1023.79 | 75 |
| 1023.86 | 76 |
| 1023.88 | 77 |
| 1023.91 | 78 |
| 1023.94 | 79 |
| 1023.95 | 80 |
| 1023.97 | 81 |
| 1023.99 | 82 |
| 1024 | 83 |
| 1024.02 | 84 |
| 1024.03 | 85 |
| 1024.04 | 86 |
| 1024.06 | 87 |
| 1024.1 | 88 |
| 1024.13 | 89 |
| 1024.14 | 90 |
| 1024.18 | 91 |
| 1024.2 | 92 |
| 1024.22 | 93 |
| 1024.24 | 94 |
| 1024.26 | 95 |
| 1024.27 | 96 |
| 1024.28 | 97 |
| 1024.29 | 98 |
| 1024.29 | 99 |
| 1024.35 | 100 |
| 1024.44 | 101 |
| 1024.51 | 102 |
| 1024.56 | 103 |
| 1024.58 | 104 |
| 1024.62 | 105 |
| 1024.65 | 106 |
| 1024.74 | 107 |
| 1024.83 | 108 |
| 1024.89 | 109 |
| 1024.94 | 110 |
| 1025 | 111 |
| 1025.09 | 112 |
| 1025.13 | 113 |
| 1025.14 | 114 |
| 1025.15 | 115 |
| 1025.16 | 116 |
| 1025.17 | 117 |
| 1025.19 | 118 |
| 1025.21 | 119 |
| 1025.24 | 120 |
| 1025.32 | 121 |
| 1025.34 | 122 |
| 1025.36 | 123 |
| 1025.4 | 124 |
| 1025.42 | 125 |
| 1025.46 | 126 |
| 1025.5 | 127 |
| 1025.54 | 128 |
| 1025.58 | 129 |
| 1025.61 | 130 |
| 1025.61 | 131 |
| 1025.62 | 132 |
| 1025.63 | 133 |
| 1025.64 | 134 |
| 1025.65 | 135 |
| 1025.68 | 136 |
| 1025.7 | 137 |
| 1025.74 | 138 |
| 1025.77 | 139 |
| 1025.82 | 140 |
| 1025.85 | 141 |
| 1025.87 | 142 |
| 1025.9 | 143 |
| 1025.93 | 144 |
| 1025.94 | 145 |
| 1025.95 | 146 |
| 1025.96 | 147 |
| 1025.96 | 148 |
| 1025.97 | 149 |
| 1025.99 | 150 |
| 1026.02 | 151 |
| 1026.03 | 152 |
| 1026.05 | 153 |
| 1026.06 | 154 |
| 1026.07 | 155 |
| 1026.09 | 156 |
| 1026.11 | 157 |
| 1026.14 | 158 |
| 1026.16 | 159 |
| 1026.22 | 160 |
| 1026.27 | 161 |
| 1026.28 | 162 |
| 1026.31 | 163 |
| 1026.38 | 164 |
| 1026.4 | 165 |
| 1026.43 | 166 |
| 1026.44 | 167 |
| 1026.45 | 168 |
| 1026.47 | 169 |
| 1026.49 | 170 |
| 1026.51 | 171 |
| 1026.55 | 172 |
| 1026.57 | 173 |
| 1026.58 | 174 |
| 1026.6 | 175 |
| 1026.62 | 176 |
| 1026.65 | 177 |
| 1026.69 | 178 |
| 1026.72 | 179 |
| 1026.73 | 180 |
| 1026.74 | 181 |
| 1026.76 | 182 |
| 1026.79 | 183 |
| 1026.81 | 184 |
| 1026.89 | 185 |
| 1026.92 | 186 |
| 1026.94 | 187 |
| 1026.95 | 188 |
| 1026.97 | 189 |
| 1027 | 190 |
| 1027.02 | 191 |
| 1027.03 | 192 |
| 1027.05 | 193 |
| 1027.05 | 194 |
| 1027.06 | 195 |
| 1027.06 | 196 |
| 1027.07 | 197 |
| 1027.07 | 198 |
| 1027.08 | 199 |
| 1027.08 | 200 |
| 1027.09 | 201 |
| 1027.09 | 202 |

| Density (kg m-3) | M600 |
| --- | --- |
| 1022.38 | 3 |
| 1022.41 | 4 |
| 1022.46 | 5 |
| 1022.49 | 6 |
| 1022.53 | 7 |
| 1022.54 | 8 |
| 1022.56 | 9 |
| 1022.58 | 10 |
| 1022.6 | 11 |
| 1022.61 | 12 |
| 1022.63 | 13 |
| 1022.64 | 14 |
| 1022.65 | 15 |
| 1022.66 | 16 |
| 1022.67 | 17 |
| 1022.67 | 18 |
| 1022.68 | 19 |
| 1022.69 | 20 |
| 1022.69 | 21 |
| 1022.7 | 22 |
| 1022.7 | 23 |
| 1022.71 | 24 |
| 1022.71 | 25 |
| 1022.72 | 26 |
| 1022.72 | 27 |
| 1022.73 | 28 |
| 1022.73 | 29 |
| 1022.74 | 30 |
| 1022.74 | 31 |
| 1022.75 | 32 |
| 1022.75 | 33 |
| 1022.76 | 34 |
| 1022.76 | 35 |
| 1022.77 | 36 |
| 1022.77 | 37 |
| 1022.78 | 38 |
| 1022.78 | 39 |
| 1022.78 | 40 |
| 1022.79 | 41 |
| 1022.8 | 42 |
| 1022.81 | 43 |
| 1022.83 | 44 |
| 1022.83 | 45 |
| 1022.84 | 46 |
| 1022.85 | 47 |
| 1022.88 | 48 |
| 1022.91 | 49 |
| 1022.95 | 50 |
| 1022.99 | 51 |
| 1023.02 | 52 |
| 1023.05 | 53 |
| 1023.08 | 54 |
| 1023.1 | 55 |
| 1023.12 | 56 |
| 1023.13 | 57 |
| 1023.16 | 58 |
| 1023.18 | 59 |
| 1023.2 | 60 |
| 1023.23 | 61 |
| 1023.29 | 62 |
| 1023.31 | 63 |
| 1023.35 | 64 |
| 1023.4 | 65 |
| 1023.42 | 66 |
| 1023.45 | 67 |
| 1023.48 | 68 |
| 1023.49 | 69 |
| 1023.5 | 70 |
| 1023.52 | 71 |
| 1023.53 | 72 |
| 1023.54 | 73 |
| 1023.56 | 74 |
| 1023.6 | 75 |
| 1023.64 | 76 |
| 1023.68 | 77 |
| 1023.7 | 78 |
| 1023.74 | 79 |
| 1023.77 | 80 |
| 1023.78 | 81 |
| 1023.83 | 82 |
| 1023.85 | 83 |
| 1023.88 | 84 |
| 1023.9 | 85 |
| 1023.94 | 86 |
| 1024.02 | 87 |
| 1024.06 | 88 |
| 1024.07 | 89 |
| 1024.07 | 90 |
| 1024.08 | 91 |
| 1024.09 | 92 |
| 1024.1 | 93 |
| 1024.13 | 94 |
| 1024.15 | 95 |
| 1024.19 | 96 |
| 1024.23 | 97 |
| 1024.28 | 98 |
| 1024.34 | 99 |
| 1024.37 | 100 |
| 1024.38 | 101 |
| 1024.39 | 102 |
| 1024.4 | 103 |
| 1024.42 | 104 |
| 1024.45 | 105 |
| 1024.59 | 106 |
| 1024.72 | 107 |
| 1024.79 | 108 |
| 1024.83 | 109 |
| 1024.91 | 110 |
| 1024.97 | 111 |
| 1025.09 | 112 |
| 1025.16 | 113 |
| 1025.2 | 114 |
| 1025.27 | 115 |
| 1025.3 | 116 |
| 1025.33 | 117 |
| 1025.36 | 118 |
| 1025.41 | 119 |
| 1025.45 | 120 |
| 1025.5 | 121 |
| 1025.55 | 122 |
| 1025.57 | 123 |
| 1025.59 | 124 |
| 1025.62 | 125 |
| 1025.64 | 126 |
| 1025.66 | 127 |
| 1025.69 | 128 |
| 1025.74 | 129 |
| 1025.78 | 130 |
| 1025.8 | 131 |
| 1025.81 | 132 |
| 1025.84 | 133 |
| 1025.89 | 134 |
| 1025.92 | 135 |
| 1025.96 | 136 |
| 1025.99 | 137 |
| 1026.02 | 138 |
| 1026.1 | 139 |
| 1026.13 | 140 |
| 1026.15 | 141 |
| 1026.18 | 142 |
| 1026.2 | 143 |
| 1026.21 | 144 |
| 1026.22 | 145 |
| 1026.22 | 146 |
| 1026.24 | 147 |
| 1026.27 | 148 |
| 1026.29 | 149 |
| 1026.32 | 150 |
| 1026.34 | 151 |
| 1026.36 | 152 |
| 1026.37 | 153 |
| 1026.4 | 154 |
| 1026.42 | 155 |
| 1026.45 | 156 |
| 1026.48 | 157 |
| 1026.49 | 158 |
| 1026.5 | 159 |
| 1026.54 | 160 |
| 1026.56 | 161 |
| 1026.56 | 162 |
| 1026.57 | 163 |
| 1026.58 | 164 |
| 1026.58 | 165 |
| 1026.59 | 166 |
| 1026.6 | 167 |
| 1026.63 | 168 |
| 1026.64 | 169 |
| 1026.66 | 170 |
| 1026.68 | 171 |
| 1026.71 | 172 |
| 1026.72 | 173 |
| 1026.73 | 174 |
| 1026.75 | 175 |
| 1026.79 | 176 |
| 1026.81 | 177 |
| 1026.81 | 178 |
| 1026.83 | 179 |
| 1026.84 | 180 |
| 1026.85 | 181 |
| 1026.86 | 182 |
| 1026.87 | 183 |
| 1026.88 | 184 |
| 1026.88 | 185 |
| 1026.9 | 186 |
| 1026.91 | 187 |
| 1026.92 | 188 |
| 1026.93 | 189 |
| 1026.94 | 190 |
| 1026.96 | 191 |
| 1026.97 | 192 |
| 1026.99 | 193 |
| 1027 | 194 |
| 1027.01 | 195 |
| 1027.02 | 196 |
| 1027.03 | 197 |
| 1027.04 | 198 |
| 1027.07 | 199 |
| 1027.08 | 200 |
| 1027.09 | 201 |
| 1027.12 | 202 |

| Density (kg m-3) | T600 |
| --- | --- |
| 1021.35 | 3 |
| 1021.38 | 4 |
| 1021.41 | 5 |
| 1021.44 | 6 |
| 1021.46 | 7 |
| 1021.47 | 8 |
| 1021.48 | 9 |
| 1021.49 | 10 |
| 1021.5 | 11 |
| 1021.51 | 12 |
| 1021.53 | 13 |
| 1021.54 | 14 |
| 1021.56 | 15 |
| 1021.57 | 16 |
| 1021.59 | 17 |
| 1021.6 | 18 |
| 1021.61 | 19 |
| 1021.61 | 20 |
| 1021.63 | 21 |
| 1021.66 | 22 |
| 1021.7 | 23 |
| 1021.78 | 24 |
| 1021.8 | 25 |
| 1021.82 | 26 |
| 1021.88 | 27 |
| 1021.92 | 28 |
| 1021.95 | 29 |
| 1021.96 | 30 |
| 1021.99 | 31 |
| 1022.05 | 32 |
| 1022.09 | 33 |
| 1022.1 | 34 |
| 1022.12 | 35 |
| 1022.12 | 36 |
| 1022.13 | 37 |
| 1022.14 | 38 |
| 1022.21 | 39 |
| 1022.3 | 40 |
| 1022.33 | 41 |
| 1022.36 | 42 |
| 1022.37 | 43 |
| 1022.41 | 44 |
| 1022.45 | 45 |
| 1022.47 | 46 |
| 1022.5 | 47 |
| 1022.51 | 48 |
| 1022.54 | 49 |
| 1022.57 | 50 |
| 1022.58 | 51 |
| 1022.6 | 52 |
| 1022.65 | 53 |
| 1022.68 | 54 |
| 1022.7 | 55 |
| 1022.74 | 56 |
| 1022.77 | 57 |
| 1022.83 | 58 |
| 1022.84 | 59 |
| 1022.86 | 60 |
| 1022.88 | 61 |
| 1022.91 | 62 |
| 1022.93 | 63 |
| 1022.95 | 64 |
| 1022.97 | 65 |
| 1022.98 | 66 |
| 1022.98 | 67 |
| 1022.99 | 68 |
| 1023 | 69 |
| 1023.01 | 70 |
| 1023.02 | 71 |
| 1023.04 | 72 |
| 1023.07 | 73 |
| 1023.13 | 74 |
| 1023.18 | 75 |
| 1023.21 | 76 |
| 1023.24 | 77 |
| 1023.25 | 78 |
| 1023.27 | 79 |
| 1023.28 | 80 |
| 1023.3 | 81 |
| 1023.32 | 82 |
| 1023.35 | 83 |
| 1023.37 | 84 |
| 1023.39 | 85 |
| 1023.4 | 86 |
| 1023.42 | 87 |
| 1023.43 | 88 |
| 1023.44 | 89 |
| 1023.47 | 90 |
| 1023.5 | 91 |
| 1023.51 | 92 |
| 1023.53 | 93 |
| 1023.54 | 94 |
| 1023.56 | 95 |
| 1023.6 | 96 |
| 1023.62 | 97 |
| 1023.63 | 98 |
| 1023.64 | 99 |
| 1023.66 | 100 |
| 1023.67 | 101 |
| 1023.68 | 102 |
| 1023.72 | 103 |
| 1023.75 | 104 |
| 1023.77 | 105 |
| 1023.8 | 106 |
| 1023.84 | 107 |
| 1023.93 | 108 |
| 1023.96 | 109 |
| 1023.99 | 110 |
| 1024.03 | 111 |
| 1024.1 | 112 |
| 1024.14 | 113 |
| 1024.17 | 114 |
| 1024.19 | 115 |
| 1024.21 | 116 |
| 1024.22 | 117 |
| 1024.22 | 118 |
| 1024.23 | 119 |
| 1024.24 | 120 |
| 1024.27 | 121 |
| 1024.32 | 122 |
| 1024.38 | 123 |
| 1024.46 | 124 |
| 1024.52 | 125 |
| 1024.53 | 126 |
| 1024.55 | 127 |
| 1024.56 | 128 |
| 1024.57 | 129 |
| 1024.65 | 130 |
| 1024.69 | 131 |
| 1024.71 | 132 |
| 1024.8 | 133 |
| 1024.89 | 134 |
| 1024.97 | 135 |
| 1025.05 | 136 |
| 1025.12 | 137 |
| 1025.13 | 138 |
| 1025.14 | 139 |
| 1025.15 | 140 |
| 1025.18 | 141 |
| 1025.23 | 142 |
| 1025.26 | 143 |
| 1025.29 | 144 |
| 1025.31 | 145 |
| 1025.35 | 146 |
| 1025.41 | 147 |
| 1025.46 | 148 |
| 1025.51 | 149 |
| 1025.55 | 150 |
| 1025.56 | 151 |
| 1025.61 | 152 |
| 1025.67 | 153 |
| 1025.74 | 154 |
| 1025.84 | 155 |
| 1025.89 | 156 |
| 1025.93 | 157 |
| 1025.95 | 158 |
| 1025.96 | 159 |
| 1025.99 | 160 |
| 1026.06 | 161 |
| 1026.12 | 162 |
| 1026.21 | 163 |
| 1026.24 | 164 |
| 1026.25 | 165 |
| 1026.29 | 166 |
| 1026.33 | 167 |
| 1026.35 | 168 |
| 1026.39 | 169 |
| 1026.42 | 170 |
| 1026.43 | 171 |
| 1026.44 | 172 |
| 1026.46 | 173 |
| 1026.5 | 174 |
| 1026.54 | 175 |
| 1026.64 | 176 |
| 1026.67 | 177 |
| 1026.68 | 178 |
| 1026.7 | 179 |
| 1026.71 | 180 |
| 1026.71 | 181 |
| 1026.72 | 182 |
| 1026.72 | 183 |
| 1026.73 | 184 |
| 1026.76 | 185 |
| 1026.8 | 186 |
| 1026.83 | 187 |
| 1026.85 | 188 |
| 1026.9 | 189 |
| 1026.96 | 190 |
| 1026.99 | 191 |
| 1027 | 192 |
| 1027 | 193 |
| 1027.01 | 194 |
| 1027.01 | 195 |
| 1027.02 | 196 |
| 1027.02 | 197 |
| 1027.03 | 198 |
| 1027.03 | 199 |
| 1027.04 | 200 |
| 1027.05 | 201 |
| 1027.08 | 202 |

| BVF (cycle hr-1) | G30 |
| --- | --- |
| 4.45355 | 0 |
| 4.00701 | 4 |
| 3.41299 | 5 |
| 3.45879 | 6 |
| 3.59273 | 7 |
| 3.6793 | 8 |
| 3.80548 | 9 |
| 3.45877 | 10 |
| 4.82664 | 11 |
| 8.47217 | 12 |
| 7.87509 | 13 |
| 8.41607 | 14 |
| 8.95949 | 15 |
| 8.54578 | 16 |
| 5.20298 | 17 |
| 4.38194 | 18 |
| 3.84635 | 19 |
| 4.55796 | 20 |
| 4.23579 | 21 |
| 4.955 | 22 |
| 5.99028 | 23 |
| 5.74893 | 24 |
| 4.59227 | 25 |
| 4.04567 | 26 |
| 4.00657 | 27 |
| 4.12271 | 28 |
| 5.04926 | 29 |
| 6.51854 | 30 |

| BVF (cycle hr-1) | M30 |
| --- | --- |
| 2.917 | 0 |
| 3.41472 | 3 |
| 3.63813 | 4 |
| 3.59455 | 5 |
| 3.22484 | 6 |
| 3.63811 | 7 |
| 3.72371 | 8 |
| 3.8074 | 9 |
| 3.50575 | 10 |
| 3.84854 | 11 |
| 3.72368 | 12 |
| 3.68112 | 13 |
| 4.45569 | 14 |
| 4.27521 | 15 |
| 8.28839 | 16 |
| 16.643 | 17 |
| 19.4207 | 18 |
| 13.8857 | 19 |
| 10.4253 | 20 |
| 6.19944 | 21 |
| 4.49014 | 22 |
| 4.92509 | 23 |
| 4.31115 | 24 |
| 3.72299 | 25 |
| 4.34751 | 26 |
| 3.8478 | 27 |
| 3.72297 | 28 |

| BVF (cycle hr-1) | T30 |
| --- | --- |
| 6.7396 | 0 |
| 7.59758 | 4 |
| 19.7449 | 5 |
| 8.17688 | 6 |
| 7.96183 | 7 |
| 7.74083 | 8 |
| 7.0365 | 9 |
| 9.38006 | 10 |
| 5.2379 | 11 |
| 5.6436 | 12 |
| 5.53069 | 13 |
| 5.9694 | 14 |
| 6.35322 | 15 |
| 6.92322 | 16 |
| 6.76187 | 17 |
| 8.04001 | 18 |
| 10.5353 | 19 |
| 8.78923 | 20 |
| 4.04916 | 21 |
| 4.202 | 22 |
| 4.1643 | 23 |
| 3.84954 | 24 |
| 3.80835 | 25 |
| 9.10616 | 26 |
| 10.5497 | 27 |
| 8.2331 | 28 |
| 5.94222 | 29 |
| 5.23717 | 30 |

Depth (m)

| Density (kg m-3) | G30 |
| --- | --- |
| 1022.92 | 0 |
| 1022.93 | 4 |
| 1022.93 | 5 |
| 1022.94 | 6 |
| 1022.94 | 7 |
| 1022.94 | 8 |
| 1022.95 | 9 |
| 1022.95 | 10 |
| 1022.96 | 11 |
| 1022.96 | 12 |
| 1022.99 | 13 |
| 1023.01 | 14 |
| 1023.03 | 15 |
| 1023.05 | 16 |
| 1023.08 | 17 |
| 1023.09 | 18 |
| 1023.09 | 19 |
| 1023.1 | 20 |
| 1023.1 | 21 |
| 1023.11 | 22 |
| 1023.12 | 23 |
| 1023.13 | 24 |
| 1023.14 | 25 |
| 1023.15 | 26 |
| 1023.15 | 27 |
| 1023.16 | 28 |
| 1023.16 | 29 |
| 1023.17 | 30 |

| Density (kg m-3) | M30 |
| --- | --- |
| 1021.89 | 0 |
| 1021.89 | 3 |
| 1021.9 | 4 |
| 1021.9 | 5 |
| 1021.91 | 6 |
| 1021.91 | 7 |
| 1021.91 | 8 |
| 1021.92 | 9 |
| 1021.92 | 10 |
| 1021.93 | 11 |
| 1021.93 | 12 |
| 1021.94 | 13 |
| 1021.94 | 14 |
| 1021.95 | 15 |
| 1021.95 | 16 |
| 1021.97 | 17 |
| 1022.06 | 18 |
| 1022.18 | 19 |
| 1022.24 | 20 |
| 1022.28 | 21 |
| 1022.29 | 22 |
| 1022.3 | 23 |
| 1022.3 | 24 |
| 1022.31 | 25 |
| 1022.31 | 26 |
| 1022.32 | 27 |
| 1022.32 | 28 |

| Density (kg m-3) | T30 |
| --- | --- |
| 1020.96 | 0 |
| 1020.97 | 4 |
| 1020.99 | 5 |
| 1021.12 | 6 |
| 1021.14 | 7 |
| 1021.16 | 8 |
| 1021.18 | 9 |
| 1021.19 | 10 |
| 1021.22 | 11 |
| 1021.23 | 12 |
| 1021.24 | 13 |
| 1021.25 | 14 |
| 1021.26 | 15 |
| 1021.27 | 16 |
| 1021.29 | 17 |
| 1021.3 | 18 |
| 1021.32 | 19 |
| 1021.36 | 20 |
| 1021.38 | 21 |
| 1021.39 | 22 |
| 1021.39 | 23 |
| 1021.4 | 24 |
| 1021.4 | 25 |
| 1021.41 | 26 |
| 1021.43 | 27 |
| 1021.47 | 28 |
| 1021.49 | 29 |
| 1021.5 | 30 |

| BVF (cycle hr-1) | G600 |
| --- | --- |
| 3.76411 | 0 |
| 2.17321 | 4 |
| 3.36671 | 5 |
| 3.36671 | 6 |
| 3.88753 | 7 |
| 3.63645 | 8 |
| 3.72201 | 9 |
| 3.54879 | 10 |
| 3.80565 | 11 |
| 3.27181 | 12 |
| 3.80563 | 13 |
| 3.88748 | 14 |
| 3.41308 | 15 |
| 3.59283 | 16 |
| 3.8056 | 17 |
| 3.59282 | 18 |
| 3.6794 | 19 |
| 3.54872 | 20 |
| 3.5928 | 21 |
| 3.76397 | 22 |
| 3.76396 | 23 |
| 3.76395 | 24 |
| 3.72189 | 25 |
| 3.54867 | 26 |
| 3.59275 | 27 |
| 3.76392 | 28 |
| 3.67932 | 29 |
| 4.12315 | 30 |
| 3.80549 | 31 |
| 5.38176 | 32 |
| 5.74941 | 33 |
| 10.8798 | 34 |
| 12.4831 | 35 |
| 11.2071 | 36 |
| 7.46425 | 37 |
| 4.85877 | 38 |
| 4.3458 | 39 |
| 3.84629 | 40 |
| 4.85873 | 41 |
| 6.54274 | 42 |
| 5.49696 | 43 |
| 4.76049 | 44 |
| 5.43936 | 45 |
| 4.55778 | 46 |
| 7.22827 | 47 |
| 5.26281 | 48 |
| 3.96698 | 49 |
| 5.61014 | 50 |
| 6.825 | 51 |
| 7.46372 | 52 |
| 6.54236 | 53 |
| 6.42092 | 54 |
| 7.11823 | 55 |
| 5.72101 | 56 |
| 6.2469 | 57 |
| 6.82467 | 58 |
| 9.50361 | 59 |
| 6.49373 | 60 |
| 11.0924 | 61 |
| 11.0924 | 62 |
| 11.3586 | 63 |
| 7.18364 | 64 |
| 5.4674 | 65 |
| 9.02749 | 66 |
| 11.9124 | 67 |
| 13.1548 | 68 |
| 11.6447 | 69 |
| 10.4784 | 70 |
| 12.0688 | 71 |
| 7.16074 | 72 |
| 7.93187 | 73 |
| 8.9738 | 74 |
| 14.122 | 75 |
| 8.93829 | 76 |
| 9.36773 | 77 |
| 9.40112 | 78 |
| 5.5234 | 79 |
| 7.46114 | 80 |
| 8.59703 | 81 |
| 5.49471 | 82 |
| 7.24713 | 83 |
| 6.53993 | 84 |
| 6.0399 | 85 |
| 7.20345 | 86 |
| 11.6691 | 87 |
| 9.92106 | 88 |
| 6.29455 | 89 |
| 9.92085 | 90 |
| 8.8485 | 91 |
| 7.56485 | 92 |
| 7.35402 | 93 |
| 7.87028 | 94 |
| 6.44229 | 95 |
| 6.29413 | 96 |
| 4.27033 | 97 |
| 3.59036 | 98 |
| 13.6198 | 99 |
| 16.5763 | 100 |
| 15.0028 | 101 |
| 12.1806 | 102 |
| 8.72162 | 103 |
| 11.2408 | 104 |
| 9.41456 | 105 |
| 16.828 | 106 |
| 17.0867 | 107 |
| 13.23 | 108 |
| 12.9536 | 109 |
| 13.3945 | 110 |
| 16.7033 | 111 |
| 12.2287 | 112 |
| 5.40516 | 113 |
| 5.66063 | 114 |
| 5.4051 | 115 |
| 5.85159 | 116 |
| 6.74905 | 117 |
| 7.62327 | 118 |
| 10.1968 | 119 |
| 16.2726 | 120 |
| 6.65482 | 121 |
| 8.59127 | 122 |
| 10.1961 | 123 |
| 9.52709 | 124 |
| 10.4543 | 125 |
| 11.1799 | 126 |
| 11.0668 | 127 |
| 11.9795 | 128 |
| 8.51682 | 129 |
| 4.72167 | 130 |
| 5.60358 | 131 |
| 5.82338 | 132 |
| 5.22661 | 133 |
| 6.03515 | 134 |
| 9.10455 | 135 |
| 7.90456 | 136 |
| 11.1505 | 137 |
| 9.20705 | 138 |
| 13.2831 | 139 |
| 8.85912 | 140 |
| 7.15335 | 141 |
| 9.97566 | 142 |
| 10.748 | 143 |
| 5.19573 | 144 |
| 4.3035 | 145 |
| 5.4608 | 146 |
| 4.85203 | 147 |
| 5.2855 | 148 |
| 7.86361 | 149 |
| 9.18885 | 150 |
| 6.65256 | 151 |
| 5.98175 | 152 |
| 6.23856 | 153 |
| 6.26363 | 154 |
| 6.9523 | 155 |
| 8.80468 | 156 |
| 10.1615 | 157 |
| 7.43206 | 158 |
| 13.9153 | 159 |
| 12.2732 | 160 |
| 5.13412 | 161 |
| 10.1915 | 162 |
| 14.5428 | 163 |
| 8.40222 | 164 |
| 8.42078 | 165 |
| 7.19508 | 166 |
| 5.92787 | 167 |
| 7.21675 | 168 |
| 8.53155 | 169 |
| 8.11687 | 170 |
| 10.2364 | 171 |
| 8.80271 | 172 |
| 5.9275 | 173 |
| 7.30272 | 174 |
| 7.82121 | 175 |
| 10.0815 | 176 |
| 11.4644 | 177 |
| 8.67645 | 178 |
| 5.25379 | 179 |
| 5.22383 | 180 |
| 9.63549 | 181 |
| 9.5864 | 182 |
| 6.38539 | 183 |
| 15.7905 | 184 |
| 9.86 | 185 |
| 8.05702 | 186 |
| 6.64958 | 187 |
| 7.32285 | 188 |
| 9.35364 | 189 |
| 8.11488 | 190 |
| 6.99408 | 191 |
| 6.21037 | 192 |
| 4.51459 | 193 |
| 4.26456 | 194 |
| 3.71438 | 195 |
| 3.71437 | 196 |
| 3.8389 | 197 |
| 3.71435 | 198 |
| 3.9595 | 199 |
| 3.67188 | 200 |
| 3.71433 | 201 |
| 4.26449 | 202 |

| BVF (cycle hr-1) | M600 |
| --- | --- |
| 8.97989 | 0 |
| 12.6372 | 4 |
| 10.4091 | 5 |
| 11.1539 | 6 |
| 6.42325 | 7 |
| 7.52926 | 8 |
| 7.33855 | 9 |
| 7.18672 | 10 |
| 7.715 | 11 |
| 6.75749 | 12 |
| 6.71067 | 13 |
| 5.11251 | 14 |
| 5.38254 | 15 |
| 3.96804 | 16 |
| 4.1237 | 17 |
| 5.294 | 18 |
| 4.41858 | 19 |
| 3.84711 | 20 |
| 3.88781 | 21 |
| 4.12364 | 22 |
| 4.1993 | 23 |
| 4.00744 | 24 |
| 4.16161 | 25 |
| 4.04651 | 26 |
| 4.12359 | 27 |
| 4.0074 | 28 |
| 3.84702 | 29 |
| 4.16156 | 30 |
| 3.96788 | 31 |
| 4.08518 | 32 |
| 3.54896 | 33 |
| 3.84698 | 34 |
| 3.72217 | 35 |
| 3.67962 | 36 |
| 3.67961 | 37 |
| 3.72215 | 38 |
| 3.80579 | 39 |
| 4.00729 | 40 |
| 4.38257 | 41 |
| 6.3236 | 42 |
| 7.57 | 43 |
| 4.85945 | 44 |
| 4.55855 | 45 |
| 5.77705 | 46 |
| 9.68633 | 47 |
| 8.96014 | 48 |
| 11.1375 | 49 |
| 11.444 | 50 |
| 10.0994 | 51 |
| 9.79862 | 52 |
| 9.18493 | 53 |
| 8.79975 | 54 |
| 7.71309 | 55 |
| 4.694 | 56 |
| 10.0205 | 57 |
| 8.61869 | 58 |
| 7.69236 | 59 |
| 9.45444 | 60 |
| 13.3468 | 61 |
| 7.89389 | 62 |
| 11.8208 | 63 |
| 12.0316 | 64 |
| 9.09731 | 65 |
| 9.87667 | 66 |
| 9.23438 | 67 |
| 6.11914 | 68 |
| 5.93641 | 69 |
| 6.84708 | 70 |
| 5.17152 | 71 |
| 5.55291 | 72 |
| 8.56216 | 73 |
| 11.1198 | 74 |
| 10.9772 | 75 |
| 11.5904 | 76 |
| 8.39458 | 77 |
| 10.1114 | 78 |
| 9.73075 | 79 |
| 5.77444 | 80 |
| 12.4531 | 81 |
| 9.48478 | 82 |
| 8.84989 | 83 |
| 7.16022 | 84 |
| 12.3509 | 85 |
| 15.2352 | 86 |
| 11.9094 | 87 |
| 3.45689 | 88 |
| 4.41559 | 89 |
| 5.01576 | 90 |
| 5.55141 | 91 |
| 6.41835 | 92 |
| 9.02475 | 93 |
| 8.63284 | 94 |
| 9.96823 | 95 |
| 12.3619 | 96 |
| 11.8553 | 97 |
| 14.3396 | 98 |
| 9.29824 | 99 |
| 5.34871 | 100 |
| 5.10817 | 101 |
| 5.43611 | 102 |
| 7.94915 | 103 |
| 9.64638 | 104 |
| 21.4161 | 105 |
| 20.1127 | 106 |
| 14.4457 | 107 |
| 10.783 | 108 |
| 16.6293 | 109 |
| 12.6838 | 110 |
| 19.9286 | 111 |
| 14.3339 | 112 |
| 12.138 | 113 |
| 14.0677 | 114 |
| 10.7074 | 115 |
| 8.79019 | 116 |
| 10.3491 | 117 |
| 12.1885 | 118 |
| 11.2363 | 119 |
| 12.265 | 120 |
| 12.3921 | 121 |
| 9.24196 | 122 |
| 8.12051 | 123 |
| 8.91312 | 124 |
| 8.73508 | 125 |
| 6.65378 | 126 |
| 10.7054 | 127 |
| 11.9917 | 128 |
| 11.1924 | 129 |
| 6.90799 | 130 |
| 6.0347 | 131 |
| 10.0074 | 132 |
| 12.237 | 133 |
| 10.6306 | 134 |
| 11.0219 | 135 |
| 8.62513 | 136 |
| 10.1621 | 137 |
| 16.0235 | 138 |
| 9.54028 | 139 |
| 7.66085 | 140 |
| 9.58928 | 141 |
| 8.11812 | 142 |
| 6.28821 | 143 |
| 4.75341 | 144 |
| 4.58537 | 145 |
| 6.00738 | 146 |
| 10.6288 | 147 |
| 7.19561 | 148 |
| 10.0207 | 149 |
| 8.27073 | 150 |
| 6.83761 | 151 |
| 6.53247 | 152 |
| 9.27215 | 153 |
| 8.15581 | 154 |
| 10.3132 | 155 |
| 9.66931 | 156 |
| 5.84783 | 157 |
| 5.22443 | 158 |
| 10.7886 | 159 |
| 6.95072 | 160 |
| 5.31358 | 161 |
| 4.3385 | 162 |
| 4.4102 | 163 |
| 4.48076 | 164 |
| 5.54464 | 165 |
| 6.0583 | 166 |
| 9.42212 | 167 |
| 4.97809 | 168 |
| 8.21234 | 169 |
| 7.51415 | 170 |
| 9.30452 | 171 |
| 6.53135 | 172 |
| 6.2866 | 173 |
| 7.28067 | 174 |
| 10.5074 | 175 |
| 7.84053 | 176 |
| 5.13278 | 177 |
| 6.57886 | 178 |
| 5.57219 | 179 |
| 6.55491 | 180 |
| 5.65592 | 181 |
| 5.19338 | 182 |
| 4.94592 | 183 |
| 4.37384 | 184 |
| 6.50674 | 185 |
| 5.25333 | 186 |
| 6.48252 | 187 |
| 5.45822 | 188 |
| 6.81268 | 189 |
| 5.95282 | 190 |
| 7.19322 | 191 |
| 7.40793 | 192 |
| 5.31246 | 193 |
| 5.87312 | 194 |
| 5.31241 | 195 |
| 5.03977 | 196 |
| 6.05702 | 197 |
| 9.37007 | 198 |
| 7.25793 | 199 |
| 6.05686 | 200 |
| 8.19141 | 201 |
| 5.28253 | 202 |

| BVF (cycle hr-1) | T600 |
| --- | --- |
| 9.12371 | 0 |
| 9.51272 | 4 |
| 10.9602 | 5 |
| 6.69104 | 6 |
| 5.96879 | 7 |
| 5.44388 | 8 |
| 5.7809 | 9 |
| 6.66729 | 10 |
| 5.11536 | 11 |
| 6.3772 | 12 |
| 6.61973 | 13 |
| 8.36577 | 14 |
| 6.2774 | 15 |
| 7.51188 | 16 |
| 5.83486 | 17 |
| 4.2389 | 18 |
| 3.72427 | 19 |
| 6.45061 | 20 |
| 10.1372 | 21 |
| 12.0154 | 22 |
| 14.9492 | 23 |
| 8.69731 | 24 |
| 8.15484 | 25 |
| 13.6131 | 26 |
| 11.5459 | 27 |
| 9.10387 | 28 |
| 7.34076 | 29 |
| 9.75535 | 30 |
| 13.1528 | 31 |
| 10.5763 | 32 |
| 7.4468 | 33 |
| 6.92041 | 34 |
| 4.4553 | 35 |
| 3.96908 | 36 |
| 5.77906 | 37 |
| 14.4639 | 38 |
| 17.2088 | 39 |
| 10.2882 | 40 |
| 8.47477 | 41 |
| 7.42463 | 42 |
| 11.1262 | 43 |
| 11.0693 | 44 |
| 8.17153 | 45 |
| 8.52976 | 46 |
| 6.07042 | 47 |
| 9.3572 | 48 |
| 10.1638 | 49 |
| 6.87316 | 50 |
| 7.75576 | 51 |
| 12.435 | 52 |
| 9.30592 | 53 |
| 8.76561 | 54 |
| 11.0535 | 55 |
| 9.32241 | 56 |
| 14.2068 | 57 |
| 4.31003 | 58 |
| 7.03077 | 59 |
| 9.09971 | 60 |
| 8.92492 | 61 |
| 8.1117 | 62 |
| 8.83609 | 63 |
| 7.29411 | 64 |
| 3.76385 | 65 |
| 4.34611 | 66 |
| 4.92344 | 67 |
| 4.76089 | 68 |
| 5.77661 | 69 |
| 5.88456 | 70 |
| 8.13065 | 71 |
| 10.5414 | 72 |
| 12.9164 | 73 |
| 13.2172 | 74 |
| 9.38771 | 75 |
| 9.89361 | 76 |
| 5.8031 | 77 |
| 6.93923 | 78 |
| 6.732 | 79 |
| 6.98434 | 80 |
| 8.7809 | 81 |
| 9.60251 | 82 |
| 8.39593 | 83 |
| 7.46323 | 84 |
| 5.52487 | 85 |
| 6.98393 | 86 |
| 5.66541 | 87 |
| 6.27168 | 88 |
| 9.28542 | 89 |
| 9.63455 | 90 |
| 6.77783 | 91 |
| 7.33515 | 92 |
| 6.82399 | 93 |
| 7.95247 | 94 |
| 11.5091 | 95 |
| 6.4687 | 96 |
| 4.75942 | 97 |
| 5.96247 | 98 |
| 7.85258 | 99 |
| 6.29601 | 100 |
| 4.92179 | 101 |
| 11.3432 | 102 |
| 10.0019 | 103 |
| 8.14707 | 104 |
| 8.97379 | 105 |
| 12.0942 | 106 |
| 16.4567 | 107 |
| 9.24926 | 108 |
| 9.9692 | 109 |
| 12.0409 | 110 |
| 13.9297 | 111 |
| 12.2089 | 112 |
| 8.61456 | 113 |
| 8.20312 | 114 |
| 8.31723 | 115 |
| 4.88843 | 116 |
| 4.75803 | 117 |
| 4.34345 | 118 |
| 5.2302 | 119 |
| 10.2171 | 120 |
| 11.855 | 121 |
| 14.4811 | 122 |
| 15.6091 | 123 |
| 13.7451 | 124 |
| 5.90688 | 125 |
| 6.70444 | 126 |
| 6.29327 | 127 |
| 6.34298 | 128 |
| 15.9068 | 129 |
| 11.3794 | 130 |
| 8.10463 | 131 |
| 16.2865 | 132 |
| 17.0125 | 133 |
| 16.2464 | 134 |
| 15.0093 | 135 |
| 15.5536 | 136 |
| 5.79777 | 137 |
| 4.88623 | 138 |
| 5.66064 | 139 |
| 8.57375 | 140 |
| 13.4281 | 141 |
| 9.47827 | 142 |
| 9.08903 | 143 |
| 8.79026 | 144 |
| 10.1963 | 145 |
| 14.1335 | 146 |
| 13.0108 | 147 |
| 12.2521 | 148 |
| 10.3936 | 149 |
| 7.68349 | 150 |
| 11.8078 | 151 |
| 13.4018 | 152 |
| 15.7794 | 153 |
| 16.8283 | 154 |
| 13.0562 | 155 |
| 10.6748 | 156 |
| 8.00229 | 157 |
| 7.36916 | 158 |
| 8.31003 | 159 |
| 15.3228 | 160 |
| 14.173 | 161 |
| 16.2076 | 162 |
| 9.22198 | 163 |
| 7.8226 | 164 |
| 9.97377 | 165 |
| 11.5754 | 166 |
| 8.85694 | 167 |
| 11.161 | 168 |
| 9.43972 | 169 |
| 5.7941 | 170 |
| 4.55055 | 171 |
| 7.51496 | 172 |
| 11.9872 | 173 |
| 10.8755 | 174 |
| 17.5698 | 175 |
| 10.2205 | 176 |
| 6.03212 | 177 |
| 6.31162 | 178 |
| 5.25385 | 179 |
| 4.88249 | 180 |
| 4.03863 | 181 |
| 4.30187 | 182 |
| 5.25378 | 183 |
| 9.27051 | 184 |
| 10.9316 | 185 |
| 10.4323 | 186 |
| 7.59654 | 187 |
| 12.4092 | 188 |
| 13.6255 | 189 |
| 9.93884 | 190 |
| 4.91382 | 191 |
| 4.11499 | 192 |
| 4.03806 | 193 |
| 3.95963 | 194 |
| 3.87962 | 195 |
| 3.83899 | 196 |
| 3.91981 | 197 |
| 4.03801 | 198 |
| 4.71839 | 199 |
| 6.10853 | 200 |
| 8.5841 | 201 |
| 7.38636 | 202 |

**Non-monsoon**

|  |  |
| --- | --- |
| 2.05594 | 0 |
| 3.8504 | 5 |
| 3.8504 | 6 |
| 15.7459 | 7 |
| 19.6084 | 8 |
| 32.976 | 9 |
| 50.5589 | 10 |
| 34.6488 | 11 |
| 20.0296 | 12 |
| 17.413 | 13 |
| 7.35876 | 14 |
| 7.52788 | 15 |
| 7.71372 | 16 |
| 9.38875 | 17 |
| 18.5241 | 18 |
| 27.3478 | 19 |
| 20.2654 | 20 |
| 13.9675 | 21 |
| 28.979 | 22 |
| 28.2384 | 23 |
| 16.9177 | 24 |
| 13.7541 | 25.001 |
| 13.8834 | 26 |
| 12.4123 | 27 |
| 10.9877 | 28 |
| 11.9602 | 29 |
| 12.8715 | 30 |

|  | M30 |
| --- | --- |
| 31.1892 | 0 |
| 42.9533 | 6 |
| 44.5015 | 7 |
| 32.4871 | 8 |
| 26.4559 | 9 |
| 25.9241 | 10 |
| 14.7041 | 11 |
| 12.8045 | 12 |
| 17.525 | 13 |
| 16.3539 | 14 |
| 13.8422 | 15 |
| 17.8079 | 16 |
| 13.9091 | 17 |
| 9.18153 | 18 |
| 8.83218 | 19 |
| 14.6785 | 20 |
| 8.7782 | 21 |
| 4.30768 | 22 |
| 4.88903 | 23 |
| 3.96551 | 24 |
| 4.04403 | 25 |
| 3.71996 | 26 |
| 3.50222 | 27 |
| 3.71995 | 28 |
| 3.22156 | 29 |

|  | T30 |
| --- | --- |
| 23.5159 | 0 |
| 5.35238 | 6.048 |
| 13.7364 | 7.035 |
| 9.51821 | 8.016 |
| 10.3211 | 9.011 |
| 9.69091 | 9.98 |
| 6.04264 | 10.975 |
| 7.38955 | 12.08 |
| 5.00735 | 13.143 |
| 4.49429 | 13.978 |
| 5.00153 | 15.1 |
| 5.47823 | 16.123 |
| 5.26305 | 17.039 |
| 5.34874 | 17.963 |
| 4.11483 | 19.006 |
| 3.96389 | 19.939 |
| 4.02659 | 20.973 |
| 4.54495 | 21.983 |
| 6.10611 | 23.023 |
| 4.86683 | 23.992 |
| 6.60586 | 24.953 |
| 6.52474 | 26.002 |
| 5.26969 | 27.027 |
| 4.61885 | 27.98 |
| 5.43448 | 29.034 |
| 3.9084 | 30.03 |

|  | G600 |
| --- | --- |
| 3.96707 | 0 |
| 3.54824 | 5 |
| 3.8869 | 6 |
| 3.63585 | 7 |
| 3.45838 | 8 |
| 4.30929 | 9 |
| 3.5482 | 10 |
| 3.27128 | 11 |
| 2.69055 | 12 |
| 3.80691 | 13 |
| 3.92515 | 14 |
| 3.9271 | 15 |
| 3.41251 | 16 |
| 3.50352 | 17 |
| 3.59223 | 18 |
| 3.80496 | 19 |
| 3.27122 | 20 |
| 3.88678 | 21 |
| 3.12355 | 22 |
| 3.50348 | 23 |
| 3.50523 | 24 |
| 3.41075 | 25 |
| 3.67874 | 26 |
| 3.54809 | 27 |
| 3.80489 | 28 |
| 3.72125 | 29 |
| 3.72124 | 30 |
| 3.96685 | 31 |
| 3.88669 | 32 |
| 3.76326 | 33 |
| 3.88667 | 34 |
| 5.01766 | 35 |
| 6.14533 | 36 |
| 6.75517 | 37 |
| 7.07372 | 38 |
| 8.79859 | 39 |
| 6.61375 | 40 |
| 8.2445 | 41 |
| 8.35814 | 42 |
| 7.71186 | 43 |
| 8.20599 | 44 |
| 9.36962 | 45 |
| 6.2965 | 46 |
| 9.86031 | 47 |
| 13.5901 | 48 |
| 12.1213 | 49 |
| 14.1564 | 50 |
| 23.6302 | 51 |
| 13.6801 | 52 |
| 7.7506 | 53 |
| 11.575 | 54 |
| 15.5514 | 55 |
| 10.9889 | 56 |
| 10.7719 | 57 |
| 13.1389 | 58 |
| 18.6988 | 59 |
| 12.8963 | 60 |
| 8.14456 | 61 |
| 13.209 | 62 |
| 11.6534 | 63 |
| 8.64947 | 64 |
| 9.95065 | 65 |
| 11.4898 | 66 |
| 11.7064 | 67 |
| 11.6793 | 68 |
| 11.9451 | 69 |
| 11.6383 | 70 |
| 9.14291 | 71 |
| 6.93399 | 72 |
| 6.8427 | 73 |
| 8.48299 | 74 |
| 8.00648 | 75 |
| 6.86543 | 76 |
| 8.21937 | 77 |
| 8.50116 | 78 |
| 6.11481 | 79 |
| 7.9074 | 80 |
| 11.7846 | 81 |
| 9.47931 | 82 |
| 10.2897 | 83 |
| 10.6053 | 84 |
| 9.41233 | 85 |
| 11.0403 | 86 |
| 11.4453 | 87 |
| 11.0967 | 88 |
| 7.70499 | 89 |
| 6.70202 | 90 |
| 5.57638 | 91 |
| 6.01009 | 92 |
| 5.74281 | 93 |
| 8.0633 | 94 |
| 10.0566 | 95 |
| 9.29357 | 96 |
| 9.9464 | 97 |
| 8.27432 | 98 |
| 8.77167 | 99 |
| 8.29309 | 100 |
| 10.8229 | 101 |
| 9.34338 | 102 |
| 6.86312 | 103 |
| 6.33983 | 104 |
| 6.23994 | 105 |
| 11.5793 | 106 |
| 9.08732 | 107 |
| 6.33954 | 108 |
| 9.32588 | 109 |
| 8.62617 | 110 |
| 9.06963 | 111 |
| 7.32709 | 112 |
| 8.89468 | 113 |
| 9.2747 | 114 |
| 9.10376 | 115 |
| 12.3899 | 116 |
| 8.34794 | 117 |
| 5.48951 | 118 |
| 4.94816 | 119 |
| 13.3122 | 120 |
| 7.89936 | 121 |
| 6.90717 | 122 |
| 6.38773 | 123 |
| 7.82329 | 124 |
| 7.30454 | 125 |
| 6.03384 | 126 |
| 5.65799 | 127 |
| 5.63016 | 128 |
| 9.7887 | 129 |
| 8.60182 | 130 |
| 7.43195 | 131 |
| 6.00747 | 132 |
| 5.65488 | 133 |
| 7.2212 | 134 |
| 5.98118 | 135 |
| 12.8232 | 136 |
| 6.41146 | 137 |
| 4.26611 | 138 |
| 4.1543 | 139 |
| 4.26609 | 140 |
| 5.73997 | 141 |
| 5.19471 | 142 |
| 5.76718 | 143 |
| 6.74516 | 144 |
| 5.43087 | 145 |
| 9.65343 | 146 |
| 7.28547 | 147 |
| 7.89774 | 148 |
| 6.21215 | 149 |
| 6.62751 | 150 |
| 7.7613 | 151 |
| 11.5743 | 152 |
| 9.71748 | 153 |
| 7.92099 | 154 |
| 4.78543 | 155 |
| 4.61863 | 156 |
| 5.1333 | 157 |
| 5.25408 | 158 |
| 8.92627 | 159 |
| 6.79058 | 160 |
| 5.07168 | 161 |
| 5.48755 | 162 |
| 4.81789 | 163 |
| 5.45885 | 164 |
| 9.78109 | 165 |
| 6.79031 | 166 |
| 7.51388 | 167 |
| 7.76023 | 168 |
| 5.34245 | 169 |
| 6.90462 | 170 |
| 5.19355 | 171 |
| 4.91426 | 172 |
| 5.9268 | 173 |
| 5.68366 | 174 |
| 4.65191 | 175 |
| 4.48018 | 176 |
| 4.94597 | 177 |
| 3.87798 | 178 |
| 3.84121 | 179 |
| 4.68543 | 180 |
| 4.30155 | 181 |
| 4.44496 | 182 |
| 5.65583 | 183 |
| 5.76564 | 184 |
| 6.38504 | 185 |
| 5.9529 | 186 |
| 5.071 | 187 |
| 6.83563 | 188 |
| 5.34199 | 189 |
| 6.74317 | 190 |
| 7.8797 | 191 |
| 7.21484 | 192 |
| 5.84632 | 193 |
| 4.65148 | 194 |
| 5.03974 | 195 |
| 4.2646 | 196 |
| 4.40919 | 197 |
| 4.68501 | 198 |
| 4.61989 | 199 |
| 8.20655 | 200 |
| 7.32237 | 201 |
| 4.15271 | 202 |
| 4.40905 | 203 |

|  |  |
| --- | --- |
| 1.12425 | 0 |
| 6.57948 | 5 |
| 16.4558 | 6 |
| 28.3231 | 7 |
| 57.9513 | 8 |
| 68.1481 | 9 |
| 30.1658 | 10 |
| 4.00814 | 11 |
| 7.3821 | 12 |
| 7.55079 | 13 |
| 5.96607 | 14 |
| 11.2947 | 15 |
| 8.2292 | 16 |
| 14.6025 | 17 |
| 14.9537 | 18 |
| 11.878 | 19 |
| 12.1529 | 20 |
| 20.9921 | 21 |
| 10.9758 | 21.999 |
| 4.79412 | 23 |
| 17.8675 | 24 |
| 25.6076 | 25 |
| 23.5362 | 26 |
| 18.3242 | 27 |
| 13.2506 | 28 |
| 12.2509 | 29 |
| 18.0367 | 30 |
| 24.0855 | 31 |
| 22.9601 | 32 |
| 17.4297 | 33 |
| 21.4781 | 34 |
| 16.1446 | 35 |
| 16.5478 | 36 |
| 13.768 | 37 |
| 9.05777 | 38 |
| 6.21796 | 39 |
| 5.98612 | 40 |
| 6.31818 | 41 |
| 9.87117 | 42 |
| 8.46542 | 43 |
| 6.63346 | 44 |
| 7.94825 | 45 |
| 5.95948 | 46 |
| 6.84321 | 47 |
| 9.4143 | 48 |
| 5.57796 | 49 |
| 4.48482 | 50 |
| 6.79693 | 51 |
| 5.74441 | 52 |
| 5.49268 | 53 |
| 5.57782 | 54 |
| 3.54548 | 55 |
| 4.41407 | 56 |
| 4.72358 | 57 |
| 6.58537 | 58 |
| 3.92406 | 59 |
| 3.84314 | 60 |
| 4.91905 | 61 |
| 5.13776 | 62 |
| 5.04516 | 63 |
| 5.82563 | 64 |
| 5.52096 | 65 |
| 4.48453 | 66 |
| 6.14067 | 67 |
| 6.62933 | 68 |
| 7.18222 | 69.001 |
| 5.68899 | 70 |
| 4.68989 | 71 |
| 5.34728 | 72 |
| 5.4922 | 73 |
| 5.6612 | 74 |
| 4.88667 | 75 |
| 4.65618 | 76 |
| 5.57726 | 77 |
| 4.00301 | 78 |
| 3.88348 | 79 |
| 5.57721 | 80 |
| 6.14027 | 81 |
| 8.42653 | 82 |
| 6.5607 | 83 |
| 4.65598 | 84 |
| 5.52041 | 85 |
| 7.7261 | 86 |
| 10.0109 | 87 |
| 12.464 | 88 |
| 11.5195 | 89.001 |
| 6.90995 | 90 |
| 9.56085 | 91 |
| 4.98147 | 92 |
| 4.34128 | 93 |
| 4.48364 | 94 |
| 6.67857 | 95 |
| 7.66403 | 96 |
| 7.06693 | 97 |
| 6.84104 | 98 |
| 6.77177 | 99 |
| 8.08266 | 100 |
| 6.72511 | 101 |
| 8.73612 | 102 |
| 6.77153 | 103 |
| 6.51139 | 104 |
| 7.66346 | 105 |
| 7.13276 | 106 |
| 8.29314 | 107 |
| 9.00123 | 108 |
| 5.57576 | 109 |
| 4.98079 | 110 |
| 7.30282 | 111 |
| 7.41678 | 112.001 |
| 9.15654 | 113 |
| 5.1663 | 114 |
| 5.60361 | 115 |
| 9.05287 | 116 |
| 10.5727 | 117 |
| 8.00328 | 118 |
| 6.70065 | 119 |
| 9.65659 | 120 |
| 11.8466 | 121 |
| 6.08655 | 122 |
| 8.40463 | 123 |
| 4.30377 | 124 |
| 5.13524 | 125 |
| 5.4323 | 126 |
| 4.37605 | 127 |
| 5.63089 | 128 |
| 5.92957 | 129 |
| 3.67406 | 130 |
| 5.25596 | 131 |
| 9.84891 | 132 |
| 9.42531 | 133 |
| 10.3156 | 134 |
| 7.43269 | 135 |
| 4.88419 | 136 |
| 5.19557 | 137 |
| 9.03377 | 138 |
| 9.6225 | 139 |
| 8.34732 | 140 |
| 6.72275 | 141 |
| 5.51758 | 142 |
| 4.68716 | 143 |
| 6.65225 | 144 |
| 10.3602 | 145 |
| 8.80435 | 146 |
| 18.1353 | 147 |
| 13.1491 | 148 |
| 8.69598 | 149 |
| 7.2856 | 150 |
| 10.504 | 150.999 |
| 8.76752 | 152 |
| 12.98 | 153 |
| 12.4113 | 154 |
| 13.0757 | 155 |
| 11.2157 | 156 |
| 8.36372 | 157 |
| 5.54441 | 158 |
| 5.28366 | 159 |
| 5.16354 | 160 |
| 5.90062 | 161 |
| 4.94631 | 162 |
| 3.49756 | 163 |
| 6.10644 | 164 |
| 8.73473 | 165.001 |
| 4.40981 | 166 |
| 7.3875 | 167 |
| 10.758 | 168 |
| 8.65785 | 169 |
| 5.95313 | 170 |
| 7.36592 | 171 |
| 6.26116 | 172 |
| 6.67331 | 173 |
| 6.67327 | 174 |
| 6.60235 | 175 |
| 7.47134 | 176 |
| 5.95284 | 177 |
| 4.15302 | 178 |
| 4.40937 | 179 |
| 4.07678 | 180 |
| 5.97904 | 181 |
| 6.33551 | 182 |
| 7.25819 | 183 |
| 5.73805 | 184 |
| 5.19298 | 185 |
| 5.19296 | 186 |
| 5.25298 | 187 |
| 5.40012 | 188 |
| 4.97707 | 189 |
| 5.1626 | 190 |
| 4.61755 | 191 |
| 4.61753 | 192 |
| 5.16254 | 193 |
| 5.16252 | 194 |
| 5.19278 | 195 |
| 7.10496 | 196 |
| 4.9135 | 197 |
| 7.38615 | 198 |
| 7.17429 | 199 |
| 5.15975 | 199.999 |
| 5.39978 | 201 |
| 3.99869 | 202 |
| 3.87929 | 203 |

|  |  |
| --- | --- |
| 1.53803 | 0 |
| 5.32788 | 4 |
| 5.32786 | 5 |
| 4.35016 | 6 |
| 6.40324 | 7 |
| 15.1695 | 8 |
| 47.8193 | 9.441 |
| 39.3701 | 9.952 |
| 27.333 | 10.999 |
| 18.8479 | 12.001 |
| 10.3834 | 13.031 |
| 8.48039 | 13.945 |
| 11.6932 | 15.004 |
| 15.1271 | 16.081 |
| 11.3161 | 16.927 |
| 5.67811 | 18.031 |
| 7.54354 | 19.028 |
| 7.80868 | 20.008 |
| 10.3551 | 21.001 |
| 8.72774 | 22.002 |
| 11.1811 | 23.001 |
| 10.8149 | 24.009 |
| 6.25044 | 25.02 |
| 5.86257 | 25.967 |
| 5.62307 | 26.994 |
| 5.84563 | 28.029 |
| 8.23674 | 28.988 |
| 11.8841 | 30.062 |
| 12.791 | 30.912 |
| 6.41946 | 32.023 |
| 6.00337 | 32.988 |
| 9.80463 | 33.98 |
| 10.406 | 35.001 |
| 11.3762 | 36.066 |
| 12.9329 | 36.912 |
| 13.1509 | 38.038 |
| 11.5367 | 39.015 |
| 16.7008 | 40.031 |
| 15.6226 | 41.021 |
| 17.9526 | 41.986 |
| 14.9468 | 43.004 |
| 17.3979 | 44.008 |
| 9.07747 | 44.984 |
| 7.52182 | 45.966 |
| 11.5468 | 46.974 |
| 16.312 | 48.015 |
| 14.7013 | 49.095 |
| 9.45116 | 49.93 |
| 6.80274 | 50.993 |
| 8.17981 | 52.007 |
| 6.87428 | 53.008 |
| 8.29411 | 54.017 |
| 6.21039 | 55.056 |
| 6.90113 | 55.946 |
| 7.63396 | 56.965 |
| 12.1654 | 58.003 |
| 9.22656 | 59.046 |
| 6.35822 | 59.951 |
| 6.1213 | 60.965 |
| 7.37016 | 62.005 |
| 6.31555 | 63.11 |
| 6.22951 | 63.924 |
| 6.41463 | 64.969 |
| 7.65147 | 66.026 |
| 8.75526 | 66.985 |
| 7.0839 | 67.956 |
| 6.80485 | 68.984 |
| 7.59551 | 70.054 |
| 11.005 | 70.963 |
| 9.55967 | 71.962 |
| 4.83708 | 73.009 |
| 5.42928 | 74.005 |
| 7.84951 | 74.987 |
| 8.53375 | 76.016 |
| 6.80337 | 77.008 |
| 5.94675 | 78.01 |
| 6.6446 | 79.034 |
| 6.6446 | 80.012 |
| 6.6446 | 81.032 |
| 5.0295 | 81.952 |
| 5.73346 | 83.015 |
| 7.669 | 84.019 |
| 8.15879 | 85.08 |
| 6.11347 | 85.945 |
| 6.29521 | 87.013 |
| 5.76614 | 87.963 |
| 5.06311 | 88.985 |
| 5.46897 | 89.999 |
| 7.22537 | 91.132 |
| 8.27319 | 91.932 |
| 8.06626 | 92.985 |
| 8.82233 | 94.008 |
| 9.39114 | 95.112 |
| 13.6436 | 95.926 |
| 13.561 | 96.989 |
| 6.41817 | 97.993 |
| 5.47087 | 99.069 |
| 5.24674 | 99.957 |
| 6.16896 | 100.996 |
| 10.5785 | 102.014 |
| 10.528 | 103.09 |
| 9.47862 | 103.958 |
| 9.15839 | 104.982 |
| 6.30817 | 105.989 |
| 7.74656 | 107.007 |
| 8.53892 | 108.012 |
| 11.7055 | 108.995 |
| 11.7423 | 109.982 |
| 5.90389 | 111.025 |
| 4.64175 | 111.988 |
| 4.86874 | 112.969 |
| 6.28372 | 113.994 |
| 9.65497 | 115.025 |
| 8.36291 | 116.002 |
| 7.21892 | 116.981 |
| 10.1324 | 118.004 |
| 8.50957 | 119.034 |
| 11.2505 | 119.995 |
| 6.76971 | 121.009 |
| 4.364 | 122.018 |
| 9.90212 | 123.049 |
| 11.533 | 123.955 |
| 14.2163 | 124.986 |
| 13.5272 | 125.988 |
| 12.0746 | 127.006 |
| 11.953 | 128.004 |
| 9.11253 | 129.089 |
| 8.15855 | 129.916 |
| 7.6931 | 130.957 |
| 8.57632 | 131.978 |
| 12.8107 | 133.018 |
| 7.91883 | 134.104 |
| 7.97013 | 134.938 |
| 12.9974 | 135.982 |
| 16.4268 | 136.994 |
| 8.80662 | 138.011 |
| 8.34427 | 139.043 |
| 13.1324 | 139.966 |
| 18.5966 | 140.998 |
| 10.1853 | 142.058 |
| 6.00548 | 142.928 |
| 6.31873 | 143.998 |
| 8.19611 | 145.003 |
| 5.15344 | 146.017 |
| 4.57476 | 147.001 |
| 5.25146 | 148.054 |
| 7.41962 | 148.974 |
| 5.40859 | 149.996 |
| 7.36867 | 151.003 |
| 7.51743 | 151.989 |
| 6.48579 | 153.006 |
| 5.05012 | 153.97 |
| 7.56643 | 154.972 |
| 14.8106 | 155.991 |
| 15.1572 | 157.013 |
| 10.089 | 158.042 |
| 4.56342 | 159.02 |
| 5.31976 | 159.964 |
| 5.54696 | 160.99 |
| 6.46494 | 162.022 |
| 7.17188 | 162.991 |
| 7.10501 | 163.964 |
| 6.00129 | 164.98 |
| 5.15926 | 166.017 |
| 5.20541 | 167.097 |
| 4.53659 | 167.937 |
| 5.71356 | 168.985 |
| 4.0302 | 170.013 |
| 3.98581 | 171.016 |
| 5.50737 | 171.961 |
| 7.50604 | 172.97 |
| 9.51574 | 173.995 |
| 9.36435 | 175.016 |
| 7.67393 | 176.051 |
| 7.32828 | 176.957 |
| 6.25322 | 177.989 |
| 5.46674 | 178.991 |
| 6.24917 | 179.977 |
| 10.3407 | 181.022 |
| 11.167 | 182.033 |
| 10.1872 | 182.968 |
| 5.35829 | 183.992 |
| 6.92398 | 184.992 |
| 6.23165 | 186.006 |
| 6.35244 | 187.059 |
| 5.28544 | 187.965 |
| 7.14611 | 188.995 |
| 9.26206 | 190.007 |
| 6.08174 | 191.018 |
| 4.51078 | 192.033 |
| 4.21362 | 192.969 |
| 5.1095 | 193.971 |
| 6.12242 | 195.009 |
| 7.40331 | 196.036 |
| 7.21844 | 197.024 |
| 5.24657 | 197.988 |
| 6.079 | 198.992 |
| 5.53726 | 200 |
| 5.97376 | 200.994 |
| 7.76594 | 201.98 |

|  | G600 |
| --- | --- |
| 1023.16 | 0 |
| 1023.17 | 5 |
| 1023.17 | 6 |
| 1023.18 | 7 |
| 1023.18 | 8 |
| 1023.19 | 9 |
| 1023.19 | 10 |
| 1023.2 | 11 |
| 1023.2 | 12 |
| 1023.2 | 13 |
| 1023.21 | 14 |
| 1023.21 | 15 |
| 1023.22 | 16 |
| 1023.22 | 17 |
| 1023.22 | 18 |
| 1023.23 | 19 |
| 1023.23 | 20 |
| 1023.24 | 21 |
| 1023.24 | 22 |
| 1023.24 | 23 |
| 1023.25 | 24 |
| 1023.25 | 25 |
| 1023.26 | 26 |
| 1023.26 | 27 |
| 1023.26 | 28 |
| 1023.27 | 29 |
| 1023.27 | 30 |
| 1023.28 | 31 |
| 1023.28 | 32 |
| 1023.29 | 33 |
| 1023.29 | 34 |
| 1023.3 | 35 |
| 1023.3 | 36 |
| 1023.32 | 37 |
| 1023.33 | 38 |
| 1023.35 | 39 |
| 1023.37 | 40 |
| 1023.39 | 41 |
| 1023.41 | 42 |
| 1023.43 | 43 |
| 1023.45 | 44 |
| 1023.47 | 45 |
| 1023.5 | 46 |
| 1023.51 | 47 |
| 1023.54 | 48 |
| 1023.6 | 49 |
| 1023.65 | 50 |
| 1023.71 | 51 |
| 1023.89 | 52 |
| 1023.95 | 53 |
| 1023.97 | 54 |
| 1024.01 | 55 |
| 1024.09 | 56 |
| 1024.12 | 57 |
| 1024.16 | 58 |
| 1024.22 | 59 |
| 1024.33 | 60 |
| 1024.38 | 61 |
| 1024.4 | 62 |
| 1024.46 | 63 |
| 1024.5 | 64 |
| 1024.52 | 65 |
| 1024.55 | 66 |
| 1024.6 | 67 |
| 1024.64 | 68 |
| 1024.68 | 69 |
| 1024.73 | 70 |
| 1024.77 | 71 |
| 1024.8 | 72 |
| 1024.81 | 73 |
| 1024.83 | 74 |
| 1024.85 | 75 |
| 1024.87 | 76 |
| 1024.89 | 77 |
| 1024.91 | 78 |
| 1024.93 | 79 |
| 1024.94 | 80 |
| 1024.96 | 81 |
| 1025.01 | 82 |
| 1025.04 | 83 |
| 1025.07 | 84 |
| 1025.11 | 85 |
| 1025.13 | 86 |
| 1025.17 | 87 |
| 1025.21 | 88 |
| 1025.25 | 89 |
| 1025.27 | 90 |
| 1025.29 | 91 |
| 1025.3 | 92 |
| 1025.31 | 93 |
| 1025.32 | 94 |
| 1025.34 | 95 |
| 1025.37 | 96 |
| 1025.4 | 97 |
| 1025.43 | 98 |
| 1025.45 | 99 |
| 1025.48 | 100 |
| 1025.5 | 101 |
| 1025.54 | 102 |
| 1025.56 | 103 |
| 1025.58 | 104 |
| 1025.59 | 105 |
| 1025.6 | 106 |
| 1025.65 | 107 |
| 1025.67 | 108 |
| 1025.69 | 109 |
| 1025.71 | 110 |
| 1025.74 | 111 |
| 1025.76 | 112 |
| 1025.78 | 113 |
| 1025.81 | 114 |
| 1025.83 | 115 |
| 1025.86 | 116 |
| 1025.91 | 117 |
| 1025.93 | 118 |
| 1025.94 | 119 |
| 1025.95 | 120 |
| 1026 | 121 |
| 1026.02 | 122 |
| 1026.04 | 123 |
| 1026.05 | 124 |
| 1026.07 | 125 |
| 1026.09 | 126 |
| 1026.1 | 127 |
| 1026.11 | 128 |
| 1026.12 | 129 |
| 1026.15 | 130 |
| 1026.18 | 131 |
| 1026.19 | 132 |
| 1026.2 | 133 |
| 1026.21 | 134 |
| 1026.23 | 135 |
| 1026.24 | 136 |
| 1026.3 | 137 |
| 1026.31 | 138 |
| 1026.31 | 139 |
| 1026.32 | 140 |
| 1026.33 | 141 |
| 1026.34 | 142 |
| 1026.34 | 143 |
| 1026.36 | 144 |
| 1026.37 | 145 |
| 1026.38 | 146 |
| 1026.41 | 147 |
| 1026.43 | 148 |
| 1026.45 | 149 |
| 1026.46 | 150 |
| 1026.47 | 151 |
| 1026.49 | 152 |
| 1026.53 | 153 |
| 1026.56 | 154 |
| 1026.58 | 155 |
| 1026.59 | 156 |
| 1026.6 | 157 |
| 1026.61 | 158 |
| 1026.62 | 159 |
| 1026.64 | 160 |
| 1026.66 | 161 |
| 1026.66 | 162 |
| 1026.67 | 163 |
| 1026.68 | 164 |
| 1026.69 | 165 |
| 1026.72 | 166 |
| 1026.73 | 167 |
| 1026.75 | 168 |
| 1026.77 | 169 |
| 1026.78 | 170 |
| 1026.8 | 171 |
| 1026.81 | 172 |
| 1026.81 | 173 |
| 1026.82 | 174 |
| 1026.83 | 175 |
| 1026.84 | 176 |
| 1026.85 | 177 |
| 1026.86 | 178 |
| 1026.86 | 179 |
| 1026.86 | 180 |
| 1026.87 | 181 |
| 1026.88 | 182 |
| 1026.88 | 183 |
| 1026.89 | 184 |
| 1026.9 | 185 |
| 1026.92 | 186 |
| 1026.93 | 187 |
| 1026.94 | 188 |
| 1026.95 | 189 |
| 1026.96 | 190 |
| 1026.98 | 191 |
| 1027 | 192 |
| 1027.01 | 193 |
| 1027.02 | 194 |
| 1027.03 | 195 |
| 1027.04 | 196 |
| 1027.04 | 197 |
| 1027.05 | 198 |
| 1027.06 | 199 |
| 1027.06 | 200 |
| 1027.09 | 201 |
| 1027.1 | 202 |
| 1027.11 | 203 |

|  |  |
| --- | --- |
| 1019.18 | 0 |
| 1019.18 | 5 |
| 1019.2 | 6 |
| 1019.28 | 7 |
| 1019.54 | 8 |
| 1020.6 | 9 |
| 1022.07 | 10 |
| 1022.36 | 11 |
| 1022.35 | 12 |
| 1022.37 | 13 |
| 1022.39 | 14 |
| 1022.4 | 15 |
| 1022.44 | 16 |
| 1022.46 | 17 |
| 1022.53 | 18 |
| 1022.6 | 19 |
| 1022.64 | 20 |
| 1022.69 | 21 |
| 1022.83 | 21.999 |
| 1022.87 | 23 |
| 1022.88 | 24 |
| 1022.98 | 25 |
| 1023.19 | 26 |
| 1023.36 | 27 |
| 1023.47 | 28 |
| 1023.52 | 29 |
| 1023.57 | 30 |
| 1023.68 | 31 |
| 1023.86 | 32 |
| 1024.03 | 33 |
| 1024.12 | 34 |
| 1024.27 | 35 |
| 1024.35 | 36 |
| 1024.44 | 37 |
| 1024.5 | 38 |
| 1024.53 | 39 |
| 1024.54 | 40 |
| 1024.55 | 41 |
| 1024.56 | 42 |
| 1024.59 | 43 |
| 1024.62 | 44 |
| 1024.63 | 45 |
| 1024.65 | 46 |
| 1024.66 | 47 |
| 1024.68 | 48 |
| 1024.71 | 49 |
| 1024.72 | 50 |
| 1024.72 | 51 |
| 1024.74 | 52 |
| 1024.75 | 53 |
| 1024.76 | 54 |
| 1024.77 | 55 |
| 1024.77 | 56 |
| 1024.78 | 57 |
| 1024.78 | 58 |
| 1024.8 | 59 |
| 1024.8 | 60 |
| 1024.81 | 61 |
| 1024.82 | 62 |
| 1024.82 | 63 |
| 1024.83 | 64 |
| 1024.84 | 65 |
| 1024.85 | 66 |
| 1024.86 | 67 |
| 1024.87 | 68 |
| 1024.88 | 69.001 |
| 1024.9 | 70 |
| 1024.91 | 71 |
| 1024.92 | 72 |
| 1024.93 | 73 |
| 1024.94 | 74 |
| 1024.95 | 75 |
| 1024.95 | 76 |
| 1024.96 | 77 |
| 1024.97 | 78 |
| 1024.98 | 79 |
| 1024.98 | 80 |
| 1024.99 | 81 |
| 1025 | 82 |
| 1025.03 | 83 |
| 1025.04 | 84 |
| 1025.05 | 85 |
| 1025.06 | 86 |
| 1025.08 | 87 |
| 1025.11 | 88 |
| 1025.16 | 89.001 |
| 1025.2 | 90 |
| 1025.21 | 91 |
| 1025.24 | 92 |
| 1025.25 | 93 |
| 1025.26 | 94 |
| 1025.26 | 95 |
| 1025.28 | 96 |
| 1025.3 | 97 |
| 1025.31 | 98 |
| 1025.33 | 99 |
| 1025.34 | 100 |
| 1025.36 | 101 |
| 1025.38 | 102 |
| 1025.4 | 103 |
| 1025.42 | 104 |
| 1025.43 | 105 |
| 1025.45 | 106 |
| 1025.46 | 107 |
| 1025.49 | 108 |
| 1025.51 | 109 |
| 1025.52 | 110 |
| 1025.53 | 111 |
| 1025.55 | 112.001 |
| 1025.56 | 113 |
| 1025.59 | 114 |
| 1025.6 | 115 |
| 1025.61 | 116 |
| 1025.64 | 117 |
| 1025.67 | 118 |
| 1025.69 | 119 |
| 1025.71 | 120 |
| 1025.74 | 121 |
| 1025.78 | 122 |
| 1025.79 | 123 |
| 1025.81 | 124 |
| 1025.82 | 125 |
| 1025.83 | 126 |
| 1025.84 | 127 |
| 1025.84 | 128 |
| 1025.85 | 129 |
| 1025.87 | 130 |
| 1025.87 | 131 |
| 1025.88 | 132 |
| 1025.91 | 133 |
| 1025.94 | 134 |
| 1025.97 | 135 |
| 1025.99 | 136 |
| 1026 | 137 |
| 1026.01 | 138 |
| 1026.03 | 139 |
| 1026.06 | 140 |
| 1026.08 | 141 |
| 1026.1 | 142 |
| 1026.11 | 143 |
| 1026.11 | 144 |
| 1026.13 | 145 |
| 1026.16 | 146 |
| 1026.19 | 147 |
| 1026.29 | 148 |
| 1026.35 | 149 |
| 1026.37 | 150 |
| 1026.39 | 150.999 |
| 1026.42 | 152 |
| 1026.45 | 153 |
| 1026.5 | 154 |
| 1026.55 | 155 |
| 1026.61 | 156 |
| 1026.65 | 157 |
| 1026.67 | 158 |
| 1026.68 | 159 |
| 1026.69 | 160 |
| 1026.69 | 161 |
| 1026.71 | 162 |
| 1026.71 | 163 |
| 1026.72 | 164 |
| 1026.73 | 165.001 |
| 1026.75 | 166 |
| 1026.76 | 167 |
| 1026.78 | 168 |
| 1026.81 | 169 |
| 1026.84 | 170 |
| 1026.85 | 171 |
| 1026.87 | 172 |
| 1026.88 | 173 |
| 1026.89 | 174 |
| 1026.91 | 175 |
| 1026.92 | 176 |
| 1026.94 | 177 |
| 1026.95 | 178 |
| 1026.96 | 179 |
| 1026.96 | 180 |
| 1026.97 | 181 |
| 1026.98 | 182 |
| 1026.99 | 183 |
| 1027.01 | 184 |
| 1027.02 | 185 |
| 1027.03 | 186 |
| 1027.04 | 187 |
| 1027.05 | 188 |
| 1027.05 | 189 |
| 1027.06 | 190 |
| 1027.07 | 191 |
| 1027.08 | 192 |
| 1027.08 | 193 |
| 1027.09 | 194 |
| 1027.1 | 195 |
| 1027.11 | 196 |
| 1027.13 | 197 |
| 1027.13 | 198 |
| 1027.15 | 199 |
| 1027.17 | 199.999 |
| 1027.18 | 201 |
| 1027.19 | 202 |
| 1027.19 | 203 |

|  |  |
| --- | --- |
| 1021.05 | 0 |
| 1021.05 | 4 |
| 1021.06 | 5 |
| 1021.07 | 6 |
| 1021.08 | 7 |
| 1021.09 | 8 |
| 1021.19 | 9.441 |
| 1021.56 | 9.952 |
| 1022.08 | 10.999 |
| 1022.32 | 12.001 |
| 1022.43 | 13.031 |
| 1022.46 | 13.945 |
| 1022.49 | 15.004 |
| 1022.54 | 16.081 |
| 1022.6 | 16.927 |
| 1022.64 | 18.031 |
| 1022.65 | 19.028 |
| 1022.67 | 20.008 |
| 1022.69 | 21.001 |
| 1022.72 | 22.002 |
| 1022.75 | 23.001 |
| 1022.79 | 24.009 |
| 1022.82 | 25.02 |
| 1022.84 | 25.967 |
| 1022.85 | 26.994 |
| 1022.86 | 28.029 |
| 1022.87 | 28.988 |
| 1022.89 | 30.062 |
| 1022.93 | 30.912 |
| 1022.99 | 32.023 |
| 1023 | 32.988 |
| 1023.01 | 33.98 |
| 1023.04 | 35.001 |
| 1023.08 | 36.066 |
| 1023.11 | 36.912 |
| 1023.17 | 38.038 |
| 1023.23 | 39.015 |
| 1023.27 | 40.031 |
| 1023.36 | 41.021 |
| 1023.43 | 41.986 |
| 1023.54 | 43.004 |
| 1023.61 | 44.008 |
| 1023.7 | 44.984 |
| 1023.73 | 45.966 |
| 1023.75 | 46.974 |
| 1023.79 | 48.015 |
| 1023.88 | 49.095 |
| 1023.94 | 49.93 |
| 1023.97 | 50.993 |
| 1023.98 | 52.007 |
| 1024.01 | 53.008 |
| 1024.02 | 54.017 |
| 1024.04 | 55.056 |
| 1024.05 | 55.946 |
| 1024.07 | 56.965 |
| 1024.09 | 58.003 |
| 1024.14 | 59.046 |
| 1024.16 | 59.951 |
| 1024.18 | 60.965 |
| 1024.19 | 62.005 |
| 1024.21 | 63.11 |
| 1024.22 | 63.924 |
| 1024.23 | 64.969 |
| 1024.24 | 66.026 |
| 1024.26 | 66.985 |
| 1024.29 | 67.956 |
| 1024.3 | 68.984 |
| 1024.32 | 70.054 |
| 1024.33 | 70.963 |
| 1024.37 | 71.962 |
| 1024.4 | 73.009 |
| 1024.41 | 74.005 |
| 1024.42 | 74.987 |
| 1024.44 | 76.016 |
| 1024.46 | 77.008 |
| 1024.48 | 78.01 |
| 1024.49 | 79.034 |
| 1024.5 | 80.012 |
| 1024.5 | 81.032 |
| 1024.53 | 81.952 |
| 1024.54 | 83.015 |
| 1024.55 | 84.019 |
| 1024.57 | 85.08 |
| 1024.58 | 85.945 |
| 1024.6 | 87.013 |
| 1024.61 | 87.963 |
| 1024.62 | 88.985 |
| 1024.63 | 89.999 |
| 1024.64 | 91.132 |
| 1024.65 | 91.932 |
| 1024.68 | 92.985 |
| 1024.7 | 94.008 |
| 1024.72 | 95.112 |
| 1024.75 | 95.926 |
| 1024.81 | 96.989 |
| 1024.87 | 97.993 |
| 1024.88 | 99.069 |
| 1024.89 | 99.957 |
| 1024.9 | 100.996 |
| 1024.91 | 102.014 |
| 1024.95 | 103.09 |
| 1024.98 | 103.958 |
| 1025.01 | 104.982 |
| 1025.04 | 105.989 |
| 1025.05 | 107.007 |
| 1025.07 | 108.012 |
| 1025.09 | 108.995 |
| 1025.14 | 109.982 |
| 1025.18 | 111.025 |
| 1025.19 | 111.988 |
| 1025.2 | 112.969 |
| 1025.21 | 113.994 |
| 1025.22 | 115.025 |
| 1025.25 | 116.002 |
| 1025.27 | 116.981 |
| 1025.29 | 118.004 |
| 1025.32 | 119.034 |
| 1025.34 | 119.995 |
| 1025.38 | 121.009 |
| 1025.4 | 122.018 |
| 1025.4 | 123.049 |
| 1025.43 | 123.955 |
| 1025.48 | 124.986 |
| 1025.54 | 125.988 |
| 1025.6 | 127.006 |
| 1025.65 | 128.004 |
| 1025.7 | 129.089 |
| 1025.72 | 129.916 |
| 1025.74 | 130.957 |
| 1025.76 | 131.978 |
| 1025.78 | 133.018 |
| 1025.84 | 134.104 |
| 1025.86 | 134.938 |
| 1025.88 | 135.982 |
| 1025.93 | 136.994 |
| 1026.02 | 138.011 |
| 1026.05 | 139.043 |
| 1026.07 | 139.966 |
| 1026.12 | 140.998 |
| 1026.24 | 142.058 |
| 1026.27 | 142.928 |
| 1026.28 | 143.998 |
| 1026.29 | 145.003 |
| 1026.32 | 146.017 |
| 1026.32 | 147.001 |
| 1026.33 | 148.054 |
| 1026.34 | 148.974 |
| 1026.36 | 149.996 |
| 1026.37 | 151.003 |
| 1026.38 | 151.989 |
| 1026.4 | 153.006 |
| 1026.41 | 153.97 |
| 1026.42 | 154.972 |
| 1026.44 | 155.991 |
| 1026.51 | 157.013 |
| 1026.59 | 158.042 |
| 1026.62 | 159.02 |
| 1026.63 | 159.964 |
| 1026.63 | 160.99 |
| 1026.65 | 162.022 |
| 1026.66 | 162.991 |
| 1026.67 | 163.964 |
| 1026.69 | 164.98 |
| 1026.7 | 166.017 |
| 1026.71 | 167.097 |
| 1026.72 | 167.937 |
| 1026.73 | 168.985 |
| 1026.74 | 170.013 |
| 1026.74 | 171.016 |
| 1026.75 | 171.961 |
| 1026.76 | 172.97 |
| 1026.77 | 173.995 |
| 1026.8 | 175.016 |
| 1026.83 | 176.051 |
| 1026.85 | 176.957 |
| 1026.87 | 177.989 |
| 1026.88 | 178.991 |
| 1026.89 | 179.977 |
| 1026.9 | 181.022 |
| 1026.94 | 182.033 |
| 1026.97 | 182.968 |
| 1027.01 | 183.992 |
| 1027.02 | 184.992 |
| 1027.03 | 186.006 |
| 1027.05 | 187.059 |
| 1027.06 | 187.965 |
| 1027.07 | 188.995 |
| 1027.08 | 190.007 |
| 1027.11 | 191.018 |
| 1027.12 | 192.033 |
| 1027.13 | 192.969 |
| 1027.13 | 193.971 |
| 1027.14 | 195.009 |
| 1027.16 | 196.036 |
| 1027.17 | 197.024 |
| 1027.19 | 197.988 |
| 1027.2 | 198.992 |
| 1027.21 | 200 |
| 1027.22 | 200.994 |
| 1027.23 | 201.98 |

|  | Depth (m) |
| --- | --- |
| 1020.94 | 0 |
| 1020.94 | 5 |
| 1020.95 | 6 |
| 1020.94 | 7 |
| 1021.02 | 8 |
| 1021.14 | 9 |
| 1021.49 | 10 |
| 1022.3 | 11 |
| 1022.68 | 12 |
| 1022.8 | 13 |
| 1022.9 | 14 |
| 1022.92 | 15 |
| 1022.94 | 16 |
| 1022.96 | 17 |
| 1022.98 | 18 |
| 1023.09 | 19 |
| 1023.33 | 20 |
| 1023.46 | 21 |
| 1023.52 | 22 |
| 1023.79 | 23 |
| 1024.04 | 24 |
| 1024.13 | 25.001 |
| 1024.19 | 26 |
| 1024.26 | 27 |
| 1024.3 | 28 |
| 1024.34 | 29 |
| 1024.39 | 30 |

|  | M30 |
| --- | --- |
| 1020.99 | 0 |
| 1021.3 | 6 |
| 1021.88 | 7 |
| 1022.51 | 8 |
| 1022.85 | 9 |
| 1023.07 | 10 |
| 1023.28 | 11 |
| 1023.35 | 12 |
| 1023.4 | 13 |
| 1023.5 | 14 |
| 1023.59 | 15 |
| 1023.65 | 16 |
| 1023.75 | 17 |
| 1023.81 | 18 |
| 1023.84 | 19 |
| 1023.86 | 20 |
| 1023.93 | 21 |
| 1023.95 | 22 |
| 1023.96 | 23 |
| 1023.97 | 24 |
| 1023.97 | 25 |
| 1023.98 | 26 |
| 1023.98 | 27 |
| 1023.99 | 28 |
| 1023.99 | 29 |

|  | Depth (m) |
| --- | --- |
| 1023 | 0 |
| 1022.92 | 5.083 |
| 1023.09 | 6.048 |
| 1023.1 | 7.035 |
| 1023.16 | 8.016 |
| 1023.19 | 9.011 |
| 1023.22 | 9.98 |
| 1023.25 | 10.975 |
| 1023.27 | 12.08 |
| 1023.28 | 13.143 |
| 1023.29 | 13.978 |
| 1023.3 | 15.1 |
| 1023.31 | 16.123 |
| 1023.32 | 17.039 |
| 1023.32 | 17.963 |
| 1023.33 | 19.006 |
| 1023.34 | 19.939 |
| 1023.34 | 20.973 |
| 1023.35 | 21.983 |
| 1023.36 | 23.023 |
| 1023.37 | 23.992 |
| 1023.37 | 24.953 |
| 1023.39 | 26.002 |
| 1023.4 | 27.027 |
| 1023.41 | 27.98 |
| 1023.42 | 29.034 |
| 1023.43 | 30.03 |

**Monsoon**

**Goa	Mangalore	Trivandrum**

Fig. S2. Depth profiles of density and Brunt-Vaisala frequency^[1]^ (BVF) in the water column of the coastal and off-shore stations during non-monsoon and monsoon season.­­­­

**(a)**

Observwd features

|  | G30 | G600 | M30 | M600 | T30 | T600 |
| --- | --- | --- | --- | --- | --- | --- |
| 1 | 1 | 1 | 1 | 1 | 1 | 1 |
| 1 | 1 | 1 | 1 | 1 | 1 | 1 |
| 1 | 1 | 1 | 1 | 1 | 1 | 1 |
| 1 | 1 | 1 | 1 | 1 | 1 | 1 |
| 1 | 1 | 1 | 1 | 1 | 1 | 1 |
| 1 | 1 | 1 | 1 | 1 | 1 | 1 |
| 1 | 1 | 1 | 1 | 1 | 1 | 1 |
| 1 | 1 | 1 | 1 | 1 | 1 | 1 |
| 1 | 1 | 1 | 1 | 1 | 1 | 1 |
| 10 | 1 | 1 | 1 | 1 | 1 | 1 |
| 6161 | 445.2 | 341 | 518.1 | 472.7 | 339.7 | 321.1 |
| 12322 | 538.3 | 453.2 | 619.7 | 604.9 | 434.8 | 384.2 |
| 18482 | 597.8 | 504.8 | 670.1 | 668 | 476.7 | 413 |
| 24643 | 631.4 | 535.7 | 704.2 | 718.7 | 507.4 | 436.5 |
| 30803 | 660.8 | 557.9 | 731.7 | 749.1 | 525.1 | 450.8 |
| 36964 | 683.5 | 576.1 | 753.1 | 774 | 540.4 | 458.3 |
| 43124 | 699.7 | 594.9 | 767 | 792.7 | 548.8 | 471 |
| 49285 | 714.5 | 602.6 | 776.2 | 807.4 | 556.3 | 475 |
| 55446 | 723.9 | 612.5 | 782 | 820.9 | 560.7 | 483.6 |

**(b)**

Observwd features

| Sequencing depth | G30 | G600 | M30 | M600 | T30 | T600 |
| --- | --- | --- | --- | --- | --- | --- |
| 1 | 1 | 1 | 1 | 1 | 1 | 1 |
| 1 | 1 | 1 | 1 | 1 | 1 | 1 |
| 1 | 1 | 1 | 1 | 1 | 1 | 1 |
| 1 | 1 | 1 | 1 | 1 | 1 | 1 |
| 1 | 1 | 1 | 1 | 1 | 1 | 1 |
| 1 | 1 | 1 | 1 | 1 | 1 | 1 |
| 1 | 1 | 1 | 1 | 1 | 1 | 1 |
| 1 | 1 | 1 | 1 | 1 | 1 | 1 |
| 1 | 1 | 1 | 1 | 1 | 1 | 1 |
| 10 | 1 | 1 | 1 | 1 | 1 | 1 |
| 6161 | 760.4 | 1120.2 | 1084 | 964.2 | 1726.2 | 1402.9 |
| 12322 | 1028.9 | 1516.5 | 1457.7 | 1268.7 | 2156.6 | 1803.2 |
| 18482 | 1178.5 | 1715.5 | 1633.9 | 1415.9 | 2355.2 | 2003.6 |
| 24643 | 1278.4 | 1862.4 | 1745.7 | 1507.5 | 2479.9 | 2139.3 |
| 30803 | 1350.1 | 1959.3 | 1826.3 | 1577.2 | 2569.3 | 2235 |
| 36964 | 1407.3 | 2040 | 1873.3 | 1624.8 | 2631 | 2315 |
| 43124 | 1455.1 | 2097.7 | 1923.8 | 1661.5 | 2687.8 | 2385.5 |
| 49285 | 1497.6 | 2150.7 | 1956.1 | 1695.8 | 2730.9 | 2435.8 |
| 55446 | 1533.7 | 2197.7 | 1985.4 | 1725 | 2759.6 | 2484.8 |

**Fig. S3.** Rarefactions curve at various sequencing depths for (a) non-monsoon and (b) monsoon seasons.

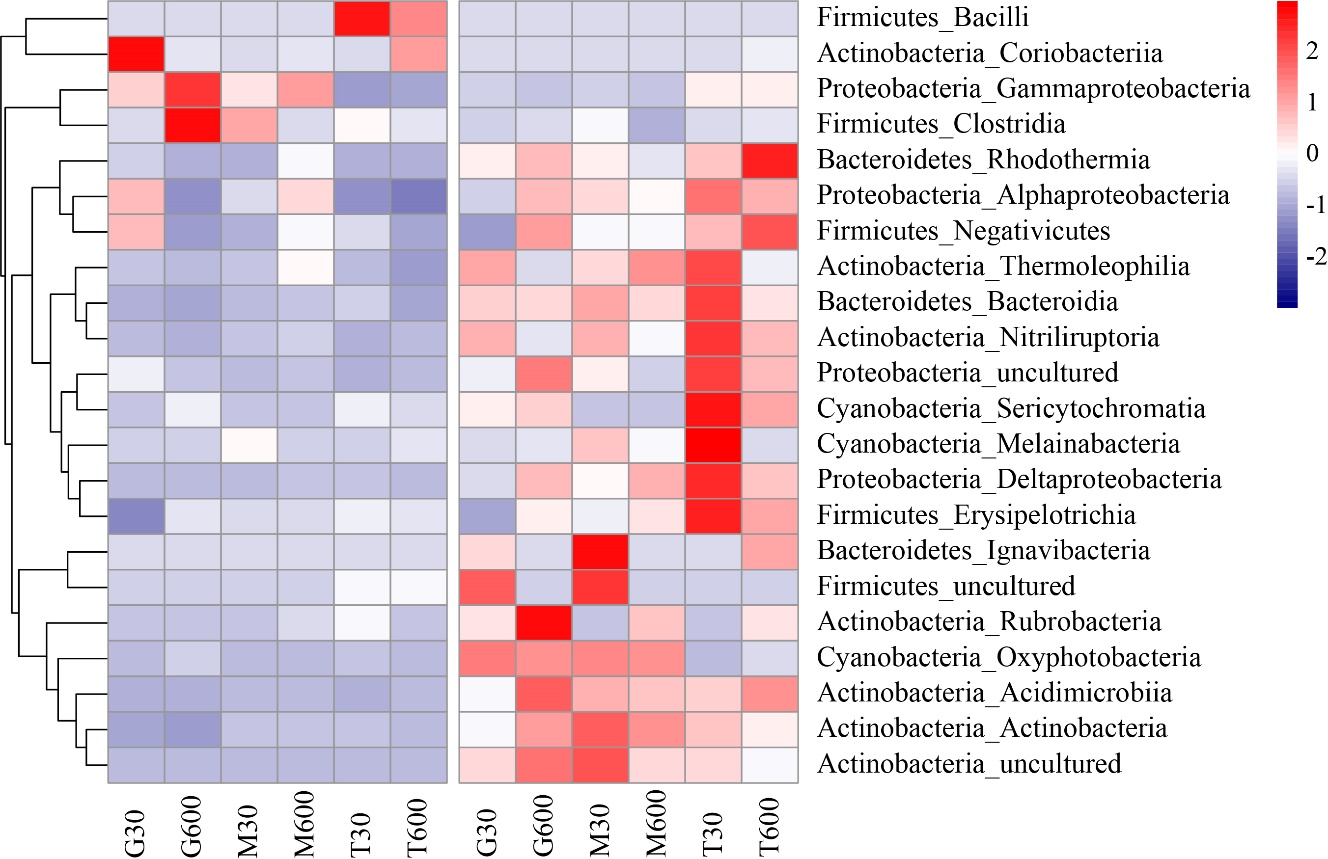

**(a)**

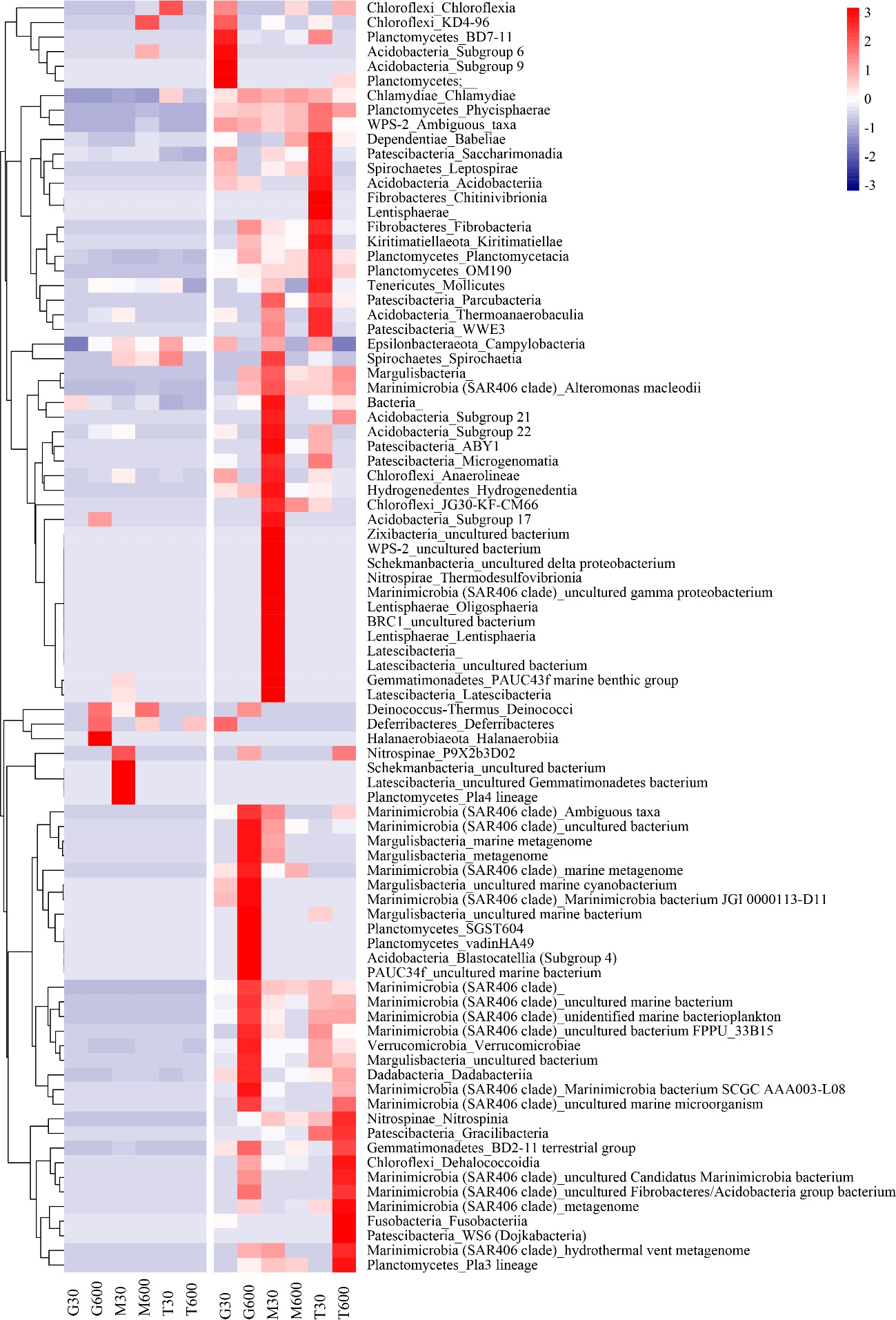

**(b)**

**Fig. S4.** Hierarchal clustering heatmap based on the Euclidian similarity index, showing the relative abundance of bacteria at class level (a) from top five phyla and (b) from other than top five phyla during non-monsoon and monsoon season

**Table S1**

Depth wise values of chlorophyll-a (Chl-a), primary productivity (PP), bacterial productivity (BP), bacterial abundance (BA), bacterial carbon (BC), and total plate count (TPC) in the water column of the coastal and off-shore stations during non-monsoon (NMS) and monsoon season (MS).

| **Stations** | **Depth** | **Chl-a**  **(mg m^-3^)** | | **PP**  **(mg C m^-3^ d^-1^)** | | **BP**  **(mg C m^-3^ d^-1^)** | | **BA**  **(Cells × 10^9^ L^-1^ )** | | **BC**  **(mg C m^-3^)** | | **TPC**  **(CFU mL^-1^)** | |
| --- | --- | --- | --- | --- | --- | --- | --- | --- | --- | --- | --- | --- | --- |
|  |  | NMS | MS | NMS | MS | NMS | MS | NMS | MS | NMS | MS | NMS | MS |
| G30 | Surface | 0.12 | 0.53 | 18.70 | 266.24 | 0.66 | 13.10 | 8.90 | 59.89 | 178.09 | 1197.85 | 152 | 5200 |
|  | C-Max | 0.11 | 1.05 | 10.77 | 83.01 | 2.01 | 13.88 | 13.73 | 62.39 | 274.51 | 1247.76 | 85 | 90000 |
|  | 30 | 0.08 | 0.07 | 8.08 | 29.62 | 10.25 | 14.68 | 9.30 | 36.53 | 186.03 | 730.51 | 1090 | 7200 |
| G600 | Surface | 0.07 | 0.06 | 11.82 | 73.44 | 3.56 | 11.38 | 13.78 | 129.54 | 275.64 | 671.52 | 500 | 6850 |
|  | C-Max | 0.09 | 0.07 | 8.68 | 59.68 | 2.77 | 7.35 | 17.92 | 39.70 | 358.45 | 2590.8 | 2240 | 1600 |
|  | 100 | 0.02 | 0.004 | 6.58 | 36.20 | 4.15 | 5.48 | 11.80 | 57.40 | 235.94 | 794.03 | 80 | 13400 |
|  | 200 | ND | ND | 5.38 | 59.83 | 4.37 | 5.62 | 11.23 | 68.97 | 224.6 | 1147.94 | 1770 | 6500 |
|  | 400 | ND | ND | ND | ND | 1.00 | 5.95 | 6.92 | 17.70 | 138.39 | 1379.34 | 580 | 36900 |
|  | 600 | ND | ND | ND | ND | 1.87 | 8.25 | 6.18 | 29.27 | 123.64 | 353.91 | 3560 | 15300 |
| M30 | Surface | 0.14 | 0.12 | 145.38 | 75.98 | 0.82 | 6.80 | 13.95 | 55.13 | 279.04 | 585.31 | 3600 | 7550 |
|  | C-Max | 0.14 | 0.35 | 48.01 | 50.11 | 6.15 | 6.96 | 12.76 | 66.02 | 255.22 | 353.91 | 4480 | 2300 |
|  | 30 | 0.12 | 0.36 | 18.85 | 33.06 | 4.41 | 7.63 | 14.24 | 45.37 | 284.72 | 1102.56 | 4020 | 17400 |
| M600 | Surface | 0.02 | 0.05 | 5.24 | 65.51 | 3.74 | 10.76 | 7.37 | 52.86 | 147.46 | 907.46 | 170 | 750 |
|  | C-Max | 0.16 | 0.23 | 8.53 | 31.56 | 29.93 | 5.28 | 8.96 | 50.14 | 179.22 | 744.12 | 3080 | 35000 |
|  | 100 | 0.01 | 0.02 | 3.14 | 30.66 | 32.42 | 6.38 | 7.15 | 51.27 | 142.92 | 694.21 | 3335 | 25000 |
|  | 200 | ND | ND | 7.48 | 29.76 | 31.39 | 9.61 | 6.81 | 31.99 | 136.12 | 1057.19 | 7640 | 11500 |
|  | 400 | ND | ND | ND | ND | 27.55 | 8.20 | 4.59 | 31.08 | 91.88 | 1002.74 | 1055 | 3550 |
|  | 600 | ND | ND | ND | ND | 30.48 | 7.21 | 3.29 | 30.17 | 65.79 | 1025.43 | 6080 | 15000 |
| T30 | Surface | 0.05 | 0.87 | 8.23 | 94.83 | 6.73 | 16.93 | 11.06 | 84.85 | 221.19 | 639.76 | 6640 | 1500 |
|  | C-Max | 0.18 | 0.40 | 11.37 | 55.64 | 7.69 | 14.38 | 10.89 | 43.78 | 217.79 | 621.61 | 475 | 250 |
|  | 30 | 0.29 | 0.29 | 10.47 | 37.99 | 5.83 | 12.23 | 10.95 | 34.71 | 218.92 | 603.46 | 17840 | 100 |
| T600 | Surface | 0.03 | 0.03 | 7.63 | 76.58 | 2.09 | 11.07 | 11.23 | 73.50 | 224.6 | 1329.43 | 18080 | 1280 |
|  | C-Max | 0.82 | 0.05 | 14.21 | 37.39 | 15.00 | 8.41 | 16.79 | 80.08 | 335.76 | 766.8 | 9320 | 830 |
|  | 100 | 0.06 | 0.04 | 1.79 | 29.91 | 20.00 | 9.65 | 9.70 | 42.65 | 193.97 | 650.35 | 3080 | 430 |
|  | 200 | ND | ND | 1.20 | 32.61 | 15.00 | 12.99 | 10.10 | 27.22 | 201.91 | 635.22 | 5080 | 860 |
|  | 400 | ND | ND | ND | ND | 25.90 | 9.26 | 8.17 | 22.69 | 163.34 | 281.31 | 3900 | 680 |
|  | 600 | ND | ND | ND | ND | 28.82 | 14.19 | 4.25 | 18.15 | 85.07 | 1470.09 | 4450 | 670 |
| ^ND: Not determined^ | | | | | | | | | | | | | |

**Table S2**

Read statistics of the amplicon sequence variants (ASVs) and operational taxonomic units (OTUs) generated.

| **Station** | **ASVs** | | | **OTUs** |
| --- | --- | --- | --- | --- |
|  | **Input** | **Merged** | **Non-chimeric** |  |
| **Non-monsoon season** | | | | |
| G30 | 1160519 | 842896 | 69470 | 740 |
| G600 | 1043626 | 795324 | 108281 | 657 |
| M30 | 871125 | 633751 | 55446 | 740 |
| M600 | 1056229 | 786561 | 82877 | 657 |
| T30 | 1211616 | 913814 | 107047 | 590 |
| T600 | 854647 | 510738 | 66070 | 492 |
| **Monsoon season** | | | | |
| G30 | 1179010 | 857178 | 138751 | 1737 |
| G600 | 1204483 | 850357 | 148939 | 2444 |
| M30 | 1255518 | 914848 | 151014 | 2172 |
| M600 | 1033245 | 736013 | 135516 | 1869 |
| T30 | 1120905 | 790040 | 100403 | 2878 |
| T600 | 1121972 | 758076 | 96196 | 2619 |

**Table S3**

Analysis of variance (ANOVA: single factor) assessing the seasonal difference in the bacterial OTUs generated and diversity indices between non-monsoon and monsoon season.

|  | **df** | **MS** | **F** | **F crit** | **P-value** |
| --- | --- | --- | --- | --- | --- |
| OTUs | 1 | 7691204 | 72.45 | 4.96 | < 0.001^**^ |
| Simpson | 1 | 0.005 | 5.83 | 4.96 | 0.036 ^*^ |
| Shannon | 1 | 8.240 | 15.22 | 4.96 | 0.003^**^ |
| Chao-1 | 1 | 7680800 | 72.75 | 4.96 | < 0.001^**^ |
| ACE | 1 | 7830975 | 73.05 | 4.96 | < 0.001^**^ |
| ^Df: degree of freedom, F value greater than F critical value indicates statistical significance. **: significant at 0.1%, *: significant at 1%.^ | | | | | |
